# Large language models complement established workflows for mechanistic model construction and refinement in systems biology

**DOI:** 10.64898/2026.11.11.766014

**Authors:** Domagoj Dorešić, Yuhong Liu, Vincent Wieland, Paul Jonas Jost, Manuel Huth, Simon Merkt, Carolina Alvarez-Garavito, Paula Rocio Moore Dillems, Christoph Thiele, Lea Seep, Dilan Pathirana, Jan Hasenauer

## Abstract

Mechanistic models provide a framework for describing, understanding, and predicting biological processes, but their development requires an interdisciplinary team with expertise in biology, mathematics, and computation. Here we present Octomod, a multi-agent workflow based on large language models (LLMs) that integrates biological knowledge retrieval, model construction, parameter estimation, and hypothesis testing. Built on established software tools, including AMICI and pyPESTO, and community standards, including SBML and PEtab, Octomod supports model development and refinement through natural-language interaction with a user. The agents’ proposals must pass defined checks and the user’s decision before they are used. Every input, model, check, and decision is recorded, so that model building remains transparent and reproducible. We applied Octomod to four problems. Three used published data and differ in model size and in how much the user guided the run: Octomod constructed a model that matched a published model without user guidance, supported the interactive refinement of a gene-regulatory model, and built models of cell–cell communication that explained the responses of five immune cell types to different stimuli. In the fourth application, on newly generated and unpublished lipidomics data, Octomod helped to determine which processes the measurements could inform, identify unresolved label fluxes, and reduce a supplied model to those processes. Together, these results show how LLMs can strengthen and complement established workflows for constructing and refining biologically informed models.

## Introduction

Mechanistic mathematical models of biological processes describe the interactions among biochemical species based on first principles. These models are used to study signal transduction, gene regulation, and metabolism (Keating et al., 2020; Gyori et al., 2017; Schmidt et al., 2011), as well as cell–cell interactions (Collier et al., 1996; Hart et al., 2014). Knowledge and hypotheses about the underlying processes are formulated in terms of reaction networks and kinetic laws, linking molecular mechanisms to temporal changes in measurable quantities. Today, researchers use different mathematical frameworks, ranging from qualitative Boolean models to complex stochastic and multiscale simulation frameworks (Hasenauer et al., 2015). Ordinary differential equation (ODE) models are among the most frequently used modelling approaches, as they provide a computationally efficient framework that is suitable for describing many cellular processes.

Model development is an iterative process that requires repeated rounds of model (re-)formulation, parameter estimation, quality-of-fit analysis, uncertainty quantification, and related tasks (Villaverde et al., 2022). Community standards and computational tools provide the foundation for this iterative modelling process. The Systems Biology Markup Language (SBML) supports the exchange and reuse of biological models (Keating et al., 2020), while Antimony provides a human-readable language for constructing models that can be translated into SBML (Smith et al., 2009). PEtab complements model descriptions with specifications of experimental conditions, measurements, observation and noise models, and parameters to be estimated (Schmiester et al., 2021). An ecosystem of tools supports numerical simulation, parameter estimation, and model analysis, including COPASI (Hoops et al., 2006), Data2Dynamics (Raue et al., 2015), AMICI and pyPESTO (Fröhlich et al., 2021; Schälte et al., 2023), and PEtab.jl (Persson et al., 2025). PEtab Select further provides a common representation of candidate model spaces and interfaces for model selection (Pathirana et al., 2026). Together, these standards provide explicit, machine-readable descriptions of models and inference problems, which the tools of this ecosystem exchange to perform their individual tasks.

Within this computational infrastructure, translating biological knowledge into model structures remains time-consuming. The relevant biochemical species and processes are often not known *a priori*. Even when they are known in principle, appropriate kinetic laws must still be selected. Furthermore, the mapping between model variables and experimental readouts may need to be identified. Knowledge-based assembly approaches have addressed parts of this task by introducing intermediate representations between biological statements and executable models. INDRA, for example, assembles mechanistic statements extracted from natural language and databases into models of signalling networks (Gyori et al., 2017). More recently, BioMASS has connected pathway knowledge and large language model (LLM) outputs to executable models through its Text2Model representation (Arakane et al., 2024). Context-dependent gene regulatory networks have also been combined with LLM-assisted integration of existing reaction equations to construct a signalling model for breast cancer (Tsutsui et al., 2026). These studies establish routes for incorporating textual knowledge and reusable model components into quantitative biological hypotheses.

When prior knowledge does not suffice for a full mechanistic description, complementary approaches infer the governing equations directly from experimental trajectories. Symbolic regression has been used to construct and refine kinetic models of metabolism, including correcting initially misspecified models (Schmidt et al., 2011), while sparse identification of nonlinear dynamics selects governing terms from a predefined library of candidate functions (Brunton et al., 2016). ODEFormer learns to infer symbolic ODE systems from trajectories using a specialised transformer (d’Ascoli et al., 2024). General-purpose LLMs provide an additional source of model proposals: LLM-SR combines generated equation programs with numerical parameter optimisation and evolutionary search, while LLM-ODE uses language-model-generated variations to guide the search over dynamical equations (Shojaee et al., 2025; Ziaei Bideh and Gryak, 2026). Recent work further combines LLM agents with constrained symbolic regression for biological ODE discovery (Krongauz et al., 2026). These approaches demonstrate different ways to search over model structures, with assumptions about the observed variables, available trajectories, and admissible equations that must be considered when applying them to biological measurements.

LLMs are also being used to coordinate broader mechanistic modelling workflows by connecting knowledge interpretation, model generation, and computational analysis. QSP-Copilot is designed to assist model development in quantitative systems pharmacology, including literature-based mechanism extraction and model formulation (Saini and Farnoud, 2025). LLM-based tool orchestration has additionally connected network construction, Boolean simulation, and multicellular modelling through natural-language interactions (Ruscone et al., 2026). LLM-based model construction has also been evaluated against human modelling efforts using the PEtab benchmark collection (Schmiester et al., 2021; Lakrisenko et al., 2026; de Pomereu et al., 2026). This comparison suggests that frontier models can outperform human modellers with respect to model selection criteria, prediction accuracy on held-out data, and the time required for model construction. Together, these developments suggest that LLMs can contribute to both the implementation and the scientific development of mechanistic models. These developments may ultimately lead to biomedical world models: representations that simulate how biological systems evolve under alternative interventions and potentially connect molecular, cellular, tissue, and clinical scales (Noori et al., 2026). Yet, building such a world model by hand is still beyond reach, because of the vast and fragmented knowledge it would have to integrate and the scarcity of time-resolved quantitative data to calibrate it. In practice, mechanistic models are therefore built to answer specific questions, at a scale the available data can support. Even this question-driven model development is slow and laborious. LLM-based agents may accelerate it by taking over steps such as gathering prior knowledge and building and calibrating candidate models, while the modeller, who owns the question, keeps the decisions.

Here, we present an LLM-based workflow that expands PEtab-based modelling, simulation, and inference using AMICI and pyPESTO, by adding capabilities for model construction and iterative refinement. The workflow translates biological descriptions and proposed revisions into SBML models and PEtab estimation problems, checks them before they are estimated and compared, and records every step in one run record. It supports both model construction from biological descriptions and the refinement of supplied models, with the evaluation of every fit informing the next proposals and the user deciding on the scope and the revisions at defined interaction points. This integration provides a proof of principle for augmenting an established systems biology workflow with LLM-assisted model development, and a proposal for documenting such development with every input, model, check, and decision.

## Methods

### Overview of the workflow

We consider the LLM-assisted construction of mechanistic ODE models to address a research question using prior knowledge and experimental data. To this end, we implement Octomod, a workflow in which users and LLM agents collaborate across eight modules coordinated by a central engine (Fig. 1A). The modules are knowledge (M0), model building (M1), model expansion (M2), data extraction (M3), PEtab assembly (M4), compilation and probing (M5), parameter estimation (M6) and evaluation (M7), followed by a revision step between rounds. Each module has a dedicated module engine that invokes agents, executes computational procedures, validates outputs, and records results (Fig. 1B). The workflow uses SBML and PEtab for model and parameter estimation problem specification, and AMICI and pyPESTO for simulation and parameter estimation (Keating et al., 2020; Schmiester et al., 2021; Fröhlich et al., 2021; Schälte et al., 2023). The agents propose elements not yet specified by the user or available information, for instance, unknown components of the base model (e.g., biochemical species and reactions) and unknown mappings between model quantities and experimental measurements. They also suggest possible explanations for model misfit and potential model revisions. Each agent is implemented as a single LLM call with a defined context (Supplementary Note S1.1), and the module engine validates the agent’s response before use (Supplementary Note S1.2). Verification agents audit the outputs of M0, M2 and M4 in separate calls that receive the output and the project description. An orchestrating agent provides the conversational interface, presenting questions and proposed choices to the user and requesting decisions or approval at interaction points (Supplementary Note S1.3). A run may comprise several rounds of model construction, fitting, comparison, and revision. All inputs, agent calls, checks, and decisions of a single run are stored in a single file called the run record. Fig. 2 shows the complete workflow: every module with its agents, checks, and interaction points.

**Figure 1:**
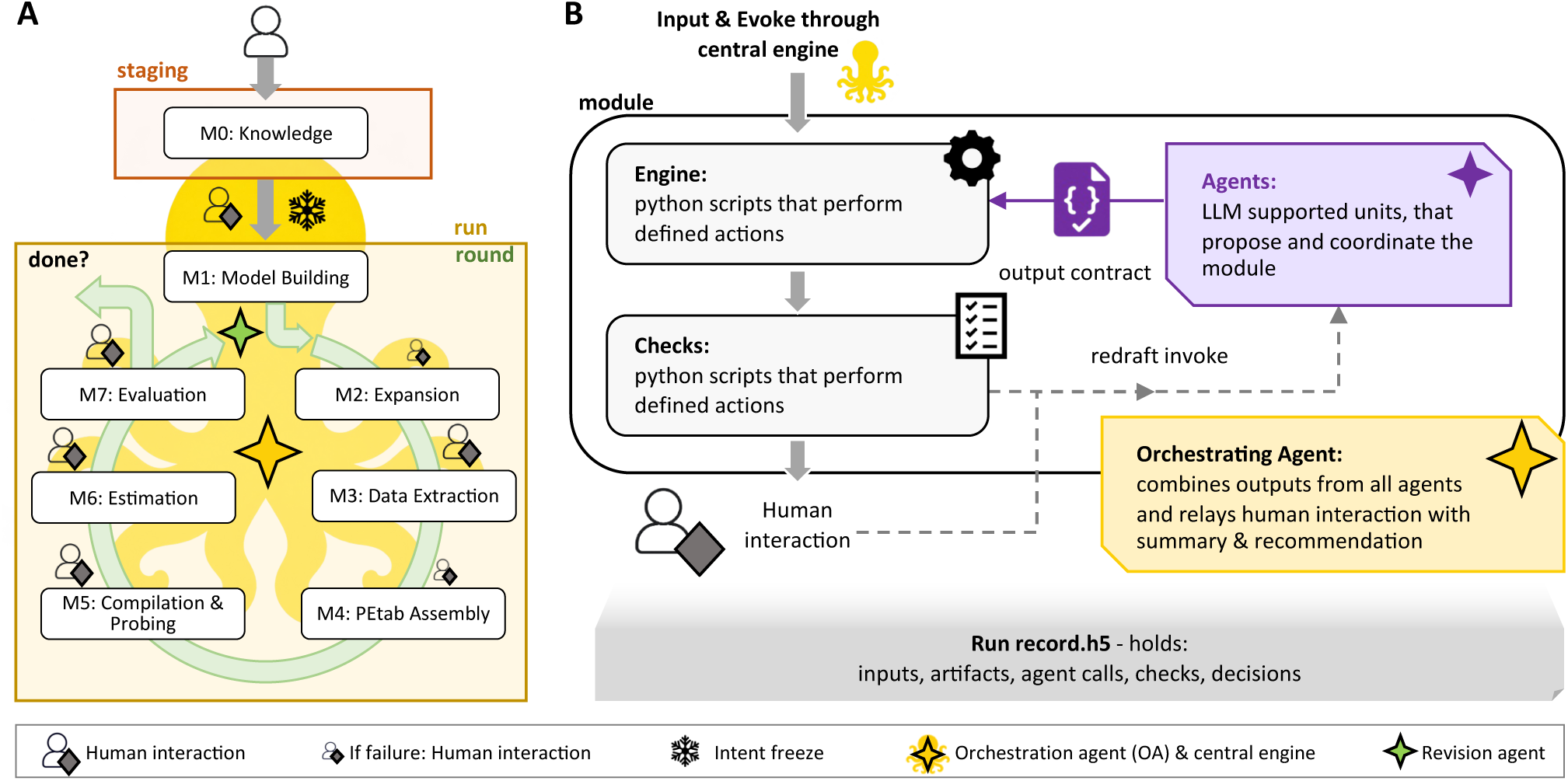
Overview of Octomod. (A) Workflow. The user supplies measurement data and a biological question and refines an intent in the so-called staging phase before freezing the intent. Then a run starts with the first round which includes module 1 (M1). Each following round goes directly to M2. (B) Architecture of a module. Each module has engine parts, agent parts, and output contracts. All parts are coordinated by the orchestrating agent and the central engine. Each step, decision and result is recorded in the run record.

**Figure 2:**
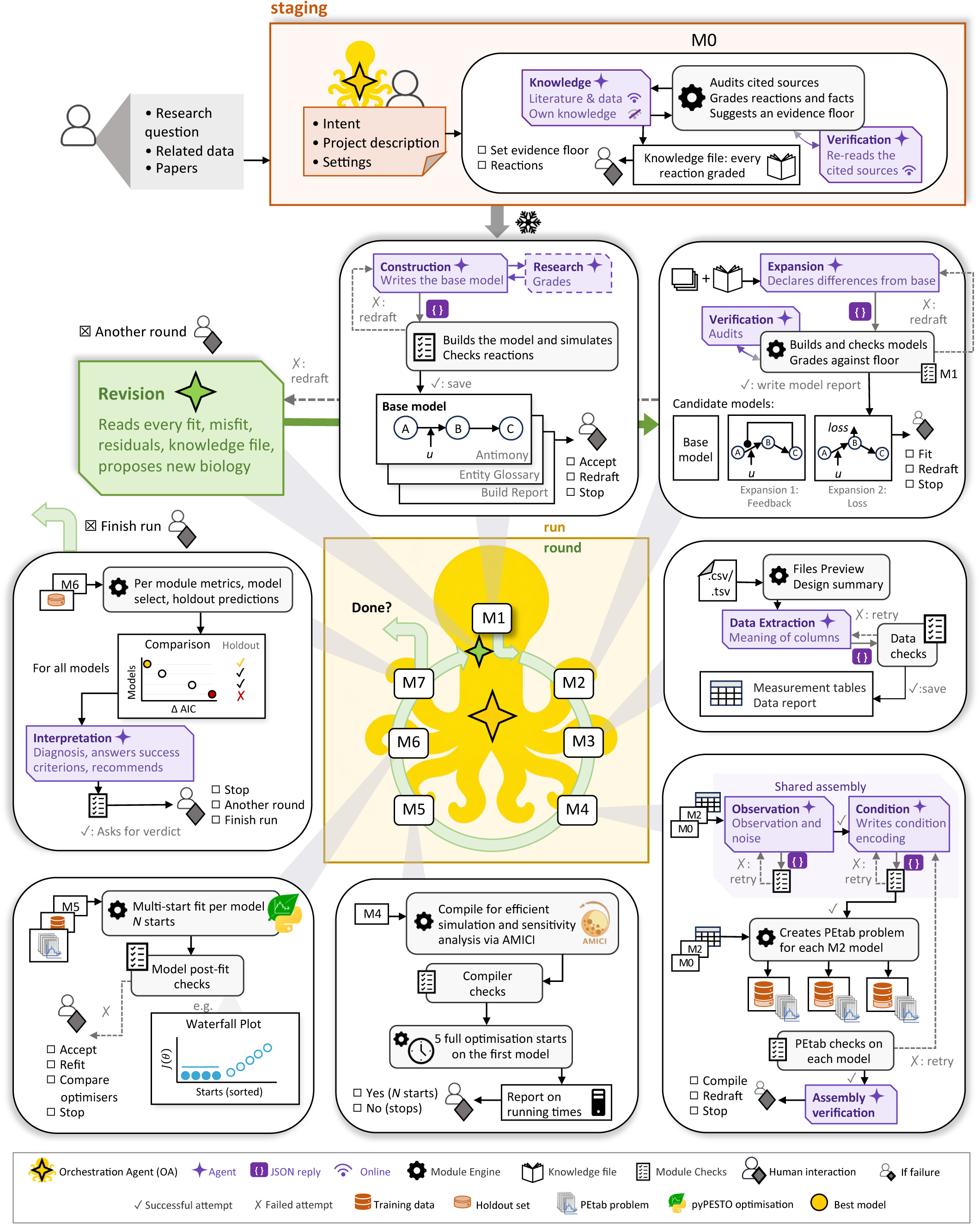
The Octomod workflow in detail. Staging (orange, top) precedes the run; the snowflake marks where the inputs are frozen. The run (yellow, centre) consists of rounds (green ring) coordinated by the orchestrating agent and the central engine. The first round starts at M1, every later round at M2. Each module is drawn as one card beside the ring, in order. Within a card, the boxes next to a person list the user’s choices at that interaction point, and a dashed arrow returns a failed check to its agent. After M7, the revision agent (green) proposes the changes for the next round or the run ends. Models, candidates and plots are illustrative. The symbols are in the legend.

### Staging and knowledge (M0): intent and knowledge file construction

The user and the orchestrating agent, an LLM session, stage the research questions, including a description of the biological systems, the experimental data, and the modelling task, along with expected outputs and success criteria (Fig. 2). Two files called intent and project description are created to contain all this information for shared use in later modules. The data files are pre-processed and the holdout data are set at this point by the user and are never chosen or accessed by an agent.

The orchestrating agent hands the intent and the project description to the knowledge agent in M0. This knowledge agent then conducts an online literature search or, in an offline mode, uses only its own pre-trained knowledge on the biological process and the experimental technique. The knowledge agent returns potentially relevant species and reactions with their sources attached. The module engine retrieves these sources, the knowledge verification agent re-reads them, and the module engine assigns each reaction an evidence grade using a fixed rule table (Supplementary Note S1.4). The grade (high, medium, low, weak or none) follows from the kind of evidence and its context. The suggested floor is the lowest grade among the essential reactions the literature supports. The user confirms or changes the evidence floor suggested by the module engine, and chooses between two modes: in strict mode, a reaction can only be added to a model when its grade is at or above the floor and in advisory mode, it can always be added, but a reaction below the floor is flagged to the user. A knowledge file is produced by the module engine.

Model building and expansion (M1, M2): construction of model candidates. We consider biochemical reaction networks, modelled as ODEs of the form

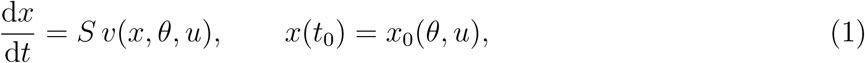

in which *x*(*t*) denotes the concentrations of the biochemical species, *S* the stoichiometric matrix, *v* the vector of reaction rates, *θ* the unknown parameters, *u* the experimental condition, and *x*(*t*_0_) is the initial state. In M1, the construction agent proposes the base model, i.e. the structure that all model candidates in all rounds are based upon, implemented in the human-readable format Antimony (Smith et al., 2009). The construction agent receives the intent, the project description, the knowledge above the evidence floor, and a verbal description of the shape of the data, but no measurement values. If the construction agent cannot answer a question using the knowledge file, then a research agent is employed to search for additional literature. The module engine converts the model from Antimony to SBML and checks by simulation, without any LLM, that the model integrates without error and that no state becomes negative. In M2, the expansion agent proposes model expansions, which can be one or more additional mechanisms, from which the module engine builds the model candidates. The construction verification agent audits them before they are presented to the user for their decision.

### Data and PEtab assembly (M3, M4): linking the model to the data

In M3, the data extraction agent determines which columns of each data file hold the observables, the conditions, and the time points according to their column names, the intent, and the project description. These are then passed to the module engine, which performs the actual conversion from data file to PEtab file in an agent-free manner, to avoid the possibility of an agent altering a measurement. In M4, the observation agent and the condition agent link the model to the data. For each observable *i*, the model observable is *y_i_*(*t, θ*) = *a_i_ h_i_*(*x*(*t, θ*)*, θ*) + *b_i_*, in which *h_i_* describes the dependence of the observable on the model state and the parameters, and *a_i_* and *b_i_* are optional scaling and offset parameters that can be hierarchically estimated from data. Each measurement *ŷ_i,j,k_* of observable *i* and replicate *j* at time *t_k_* is distributed as

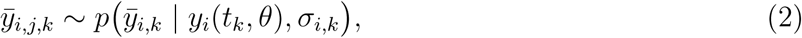

in which *p* is a normal, log-normal, Laplace or log-Laplace density and *σ_i,k_*(*x*(*t_k_, θ*)*, θ*) the noise scale, which is either fixed from the replicate spread or parameterised and estimated together with *θ*. For additive normally distributed measurement noise it holds that *ŷ_i,j,k_* = *y_i_*(*t_k_, θ*) + *ε_i,j,k_* with 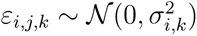. The observation agent specifies the observable functions and the noise model, and the condition agent encodes each experimental condition as the model parameters and initial states it affects. These two agents receive statistical moments and the experimental design, but no individual measurement values, such that the observation model cannot be tuned to the data. The module engine assembles and validates one PEtab problem per model candidate. The assembly verification agent audits the observable, condition and parameter tables against the experimental design and project description. The user approves the PEtab problems before any heavy computations (e.g. parameter estimation) are performed.

### Compilation and estimation (M5, M6): fitting the model candidates

Model compilation and parameter estimation are performed without any agent calls. In M5, the module engine compiles each model candidate with AMICI and projects the computation time from a sample of five optimization starts (Supplementary Note S1.5 and Table S2). After approval by the user, in M6, it estimates the parameters by minimizing the negative log-likelihood, which for (2) is given by

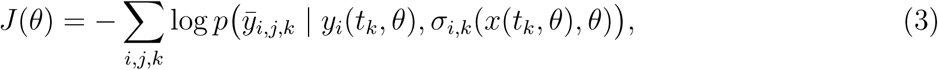

in which the sum runs over all observables and their training measurements. For normal noise, 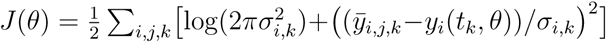. Global optimization is performed by running multi-start local optimization (Raue et al., 2013; Villaverde et al., 2022) with pyPESTO (Schalte et al., 2023), by default with 100 starts and the optimizer Fides (Fröhlich and Sorger, 2022). To evaluate optimization, the module engine checks that several starts reach the best objective function value, that the gradient is small and that few simulations failed; if a check fails, the user decides how to proceed – one option provided by Octomod is an optimizer study, which compares the best-performing optimizers reported by Grein et al. (2026) to evaluate whether there is a better optimizer for the current model.

### Evaluation (M7): comparing and diagnosing the model candidates

In M7, the module engine compares the model candidates using the Akaike information criterion (Akaike, 1974), AIC = 2*J* (*θ*^^^) + 2*n_θ_*, cross-checked with PEtab Select (Pathirana et al., 2026), which also provides the AICc and the BIC. Model candidates within two units of the best one are reported as indistinguishable (Burnham and Anderson, 2002). If holdout data are declared, normalized RMSE is computed with these data for every model candidate along with the AIC computed with the training data. For the selected model, the module engine characterizes the misfit by fitting a constant, a linear, an exponential, and a combined-exponentials functions to each time course of residuals to check for missing structure in the residuals. Lag-1 autocorrelation is also checked (Supplementary Note S1.6). Additionally, the Fisher information matrix is computed at the optimum to identify the parameters and parameter combinations that the data do not constrain. On request, it assesses parameter uncertainty using the profile likelihood, PL(*θ_l_*) = min*_θl′ ,l̸_*_=_*_l′_ J* (*θ*), from which confidence intervals are derived (Raue et al., 2009). The interpretation agent receives the fits and residuals of all model candidates, their comparison and these diagnostics of the selected model, and provides a diagnosis, hypotheses for the misfit and an answer to each success criterion. The user then gives the verdict and accepts the result or requests a revision in a further round.

### Revision: proposing the model candidates of the next round

For a further round, the revision agent receives the fitted model candidates, the misfit evidence, and the knowledge file, and proposes model revisions. It can add or remove a reaction, change a rate law, fix or free a parameter, or change the observation model. The agent is instructed to propose biological mechanisms rather than terms that merely improve the fit. The module engine checks each revision and flags those below the evidence floor, or withholds them in strict mode, and the user decides which new model candidates are fitted. The module engine builds the new model candidates from the selected model and the chosen revisions and repeats the checks of M1 to M4 without calling their agents again, as the data do not change. M5 to M7 are then run again. Only the new model candidates are estimated and compared to the selected model; thus, the search is local and does not cover all combinations of the proposed mechanisms.

### Implementation: agent calls, checks and redrafts

Each agent call is a separate commandline call of Claude Code (Anthropic, 2026) without memory of previous calls, and only the knowledge agent and the research agent can access the web. Each agent receives only a declared set of inputs, listed per agent in Supplementary Note S1.1, which the module engine enforces. A reply that violates its output contract or fails a check is returned to the agent for a redraft (Fig. 1B), and after three failed attempts the user is asked how to proceed (Supplementary Note S1.2). Such interaction is achieved through the session with the orchestrating agent. The user interacts with the session, which relays the questions at the interaction points to the user and keeps the conversation history. The session is instructed not to answer any questions itself. The settings of the workflow and of each application run are given in Supplementary Tables S2 and S3. The resulting SBML and PEtab files can be used independently of the workflow.

### Recording: one run record per run

To facilitate validation, verification, reproducibility, and dissemination of results, relevant information is stored in Run Record files; one file per run in HDF5 format (Supplementary Note S1.7). For example, all inputs, outputs, steps, and decisions during a run are saved to a run record, which can then be inspected interactively with Claude or the provided command-line utility. Other recorded information includes prompts, replies, AI model versions and monetary costs, and detailed CPU and wall timing. Octomod also provides functions to trim the run record for dissemination, or generate HTML reports of the run. In addition, the run record itself is structured according to a schema and is automatically validated, which includes checks for structure, type, and semantics.

### Agent-only analyses

To establish a baseline for comparison with the newly developed ODE-modelling workflow, we evaluated in two agent-only analyses without Octomod whether frontier LLMs (from OpenAI and Anthropic, tested separately) could independently solve the modelling task when used as orchestrators. This was done on one use case, the signalling data of peripheral blood mononuclear cells (PBMCs) from Bodenmiller et al. (2012), the third application of the Results, with internet access and without user intervention. For each case, an execution prompt was first generated from the project description using a high-capability model (OpenAI’s GPT6-Astra at ultra effort; Anthropic’s Fable 5.1) and subsequently passed to an independent orchestrator model (OpenAI’s GPT6-Sol at medium effort; Anthropic’s Opus 5.5). The orchestrator was instructed to iteratively decompose the task across multiple agents, with strict separation between implementation and verification such that no agent could evaluate its own work. The setup and the comparison are given in Supplementary Note S4.6, and the two execution prompts are in the Zenodo record (agent only analyses/).

### Cell culture

SH-SY5Y human neuroblastoma cells (ATCC CRL-2266) were cultured in DMEM/F12 and supplemented with 10% fetal calf serum, without antibiotics. For passaging, cells were washed once with PBS, detached with pre-warmed trypsin for 5 min in an incubator, and the transferred into fresh medium. For labelling experiments, cells grown to near confluence in T-75/T-175 flasks were seeded at 50% confluence into 12-well plates and labelled the following day.

### Dual-tracer pulse-chase labelling

Cells were co-labelled with alkyne-sphinganine (aSA; final concentration 1 µM) and ¹*C-labeled alkyne-palmitate (aPal; final concentration 50 µM). Both tracers were added sequentially to pre-warmed growth medium while vortexing continuously at low speed to ensure even dispersion, yielding a single combined pulse medium. For the pulse, the culture medium was replaced with 1 mL pulse medium per well, and cells were labelled for 60 min. The cells were then washed once with unlabeled growth medium and chased in unlabeled growth medium for 0, 15, 30, 60, 120, or 180 min; at 0 min, cells were harvested immediately after the chase. Each plate included one unlabeled negative control well. The experiment was done as three biological replicates, each with three technical replicates.

### Lipid extraction

At the end of the chase, the medium was removed, cells were washed with ice-cold PBS containing 1% (w/v) delipidated BSA and with ice-cold 150 mM ammonium acetate. The remaining liquid was removed. Lipids were extracted by adding 1 mL extraction mix per well: 900 µL methanol/chloroform (5:1, v/v) plus 100 µL internal standard mix. Plates were sonicated in an ultrasonic bath for 15 s. Extracts were cleared by centrifugation (20,000 × g, 5 min), and the supernatants were transferred into new tubes. Phase separation was induced with 300 µL chloroform and 600 µL water, followed by shaking and centrifugation (20,000 × g, 5 min). The lower organic phase was dried in a vacuum concentrator (45 °C, 20 min). All solvents were of MS grade.

### Click reaction and sample multiplexing

Dried lipids were dissolved in 10 µL chloroform, and subsequently 45 µL click mix was added, yielding lipids to be clicked to one of three isotopically coded alkyne-reactive reporters for multiplexing, as described (Thiele et al., 2019). The click mix was prepared from 10 µL of 100 mM reporter in 50% methanol, 200 µL of 5 mM Cu(I)(CH_3_CN)_4_BF_4_ in acetonitrile and 800 µL ethanol. Samples were sonicated for 5 min and incubated overnight at 42 *^◦^*C. Then 100 µL chloroform was added to each sample, and samples with different reporters were pooled. After adding 600 µL water to each pool and centrifuging (20,000 × g, 2 min), the lower phase was dried and redissolved in 1,250 µL spray buffer (2-propanol/methanol/water 8:5:1, v/v/v, with 10 mM ammonium acetate), sonicated for 5 min and stored at −20 °C until analysis.

### Mass spectrometry

Samples were analysed by direct infusion on a Q Exactive Plus mass spectrometer (Thermo Fisher Scientific) with a heated electrospray ionisation source. A Dionex AS-AP autosampler and an AXP-MS pump delivered the samples at 10 µL min*^−^*1, under control of Xcalibur 4.7. Source settings: sheath gas 6, auxiliary gas 2, sweep gas 0, no gas heating, spray voltage +4.1 kV / −3.2 kV, capillary temperature 280 *^◦^*C. The acquisition followed Thiele et al. (2019).

### Lipid identification

Raw files were converted to mzML with MSConvert (v 3.0.23233-c72ce16) (Adusumilli and Mallick, 2017) and analysed with LipidXplorer (v 1.2.8.1) (Herzog et al., 2012), using custom MFQL queries for the clicked, alkyne-labelled lipids. Each attached reporter adds one positive charge, so lipids carrying one, two or three labels were detected as singly, doubly or triply charged ions. For label separation, the palmitate tracer was ¹*C_6_-labeled, making each palmitate-derived alkyne heavier than a sphinganine-derived one. Labelled species were therefore assigned to the sphinganine- or palmitate-derived pool by their MS1 precursor composition. Species carrying both label types were classed as double-labelled. Where a label sits within the molecule (e.g. N-acyl chain vs. sphingoid base) was not resolved. We detected doubly labelled sphingolipids (double or triple charge), but no matching internal standards were available, so we excluded them from quantification.

### Data processing and statistics

Lipid intensities were converted to absolute amounts (pmol per well) by normalising to the class-matched internal standard. Species were grouped into lipid classes. The free fatty acid pool includes all labelled free fatty acids, not only the palmitate tracer. Within each sample, the amounts of all species in a pool were summed over both tracers. For each pool, biological replicate, and chase time, we combined the three technical replicates by median, which is robust to single outlying wells. Each biological replicate was then normalised to its own total labelled lipid amount at the end of the pulse (t = 0 min, summed over all measured classes). The resulting values are the fraction of the initial label in each pool. This normalisation removes differences in total tracer uptake between biological replicates, but not differences in the time courses. Biological replicates (n = 3, independent experiments, each with three technical replicates) were kept as separate observations for parameter estimation. Where summary statistics are given, they are mean ± SD across the three biological replicates.

## Results

### Octomod is able to construct coherent models without user guidance

To assess what Octomod achieves without user guidance, we considered a published modelling problem with an expert-built model. Callenbach et al. (2026) investigated how the kinase P-TEFb phosphorylates the eight repeats of a C-terminal domain (CTD) substrate (Fig. 3A), using mass spectrometry data that resolve the number of phosphorylated repeats at four time points (36 measurements) (Czudnochowski et al., 2012). The authors compared five hypotheses for how the kinase proceeds along the repeats: Uniform distributive, Processive, Neighbouring-effect, Global-effect and Directional (Fig. 3B). Model selection favoured the Neighbouring-effect model (7 parameters, AIC *−*166.0), indicating distributive phosphorylation with local cooperativity: phosphorylation of a neighbouring repeat accelerates phosphorylation by 2.3-fold. We supplied Octomod with the same data, an experimental description and the study question (Supplementary Note S2.1), with its knowledge module disabled (M0). No additional information or web access was provided, to avoid Search-Time Data contamination (Han et al., 2025). At each interaction point we approved the proposal (the intent, all model candidates, the default estimation settings and one revision round) without adding information or steering the choice, so that the run reflects the workflow’s own construction.

**Figure 3:**
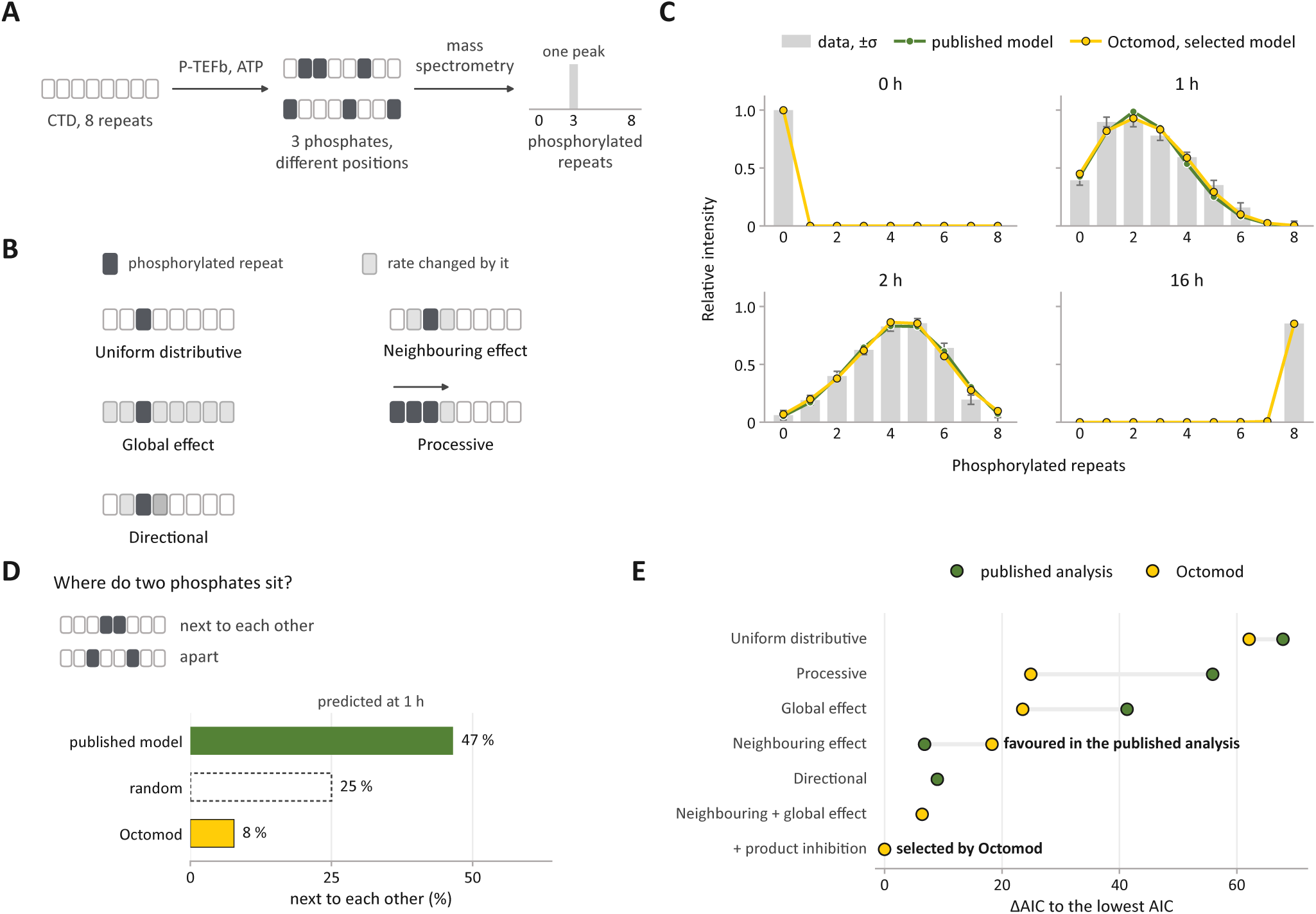
CTD data, model fits and model selection. **A** The experiment: P-TEFb phosphorylates the eight repeats of a CTD substrate. Mass spectrometry resolves the number of phosphorylated repeats, not their positions. **B** The five hypotheses of the published analysis: a phosphorylated repeat (dark), the repeats whose rate it changes (grey), the kinase proceeds along the substrate (arrow). **C** Data at the four time points (bars, *±σ* as estimated in the published analysis) with the fits of the published Neighbouring-effect model (green) and the model selected by Octomod (yellow). **D** Among CTDs with two phosphorylated repeats at 1 h, the predicted percentage with the two on adjacent repeats, for the published model, random placement and the Octomod model. **E** ΔAIC to the lowest AIC for every model of both analyses. Lines join the models that test the same hypothesis in the two analyses. The hypotheses are the same, but the rate laws differ in detail (Supplementary Note S2.4).

In M1, the construction agent proposed a base model with one species for each number of phosphorylated repeats. Unlike the published models, in which the rate is proportional to the ATP concentration, it assumes that the kinase is saturated with ATP. In M2, the expansion agent proposed five model candidates: four models correspond to different phosphorylation mechanisms (Uniform distributive, Processive, Neighbouring-effect and Global-effect model), and one combines the neighbouring and the global effect, which was not among the published hypotheses. Further-more, the neighbouring effect implementation differs: a repeat between two phosphorylated neigh-bours receives the neighbour-factor twice, whereas the published model applies the factor once. Interestingly, for the two candidates with a neighbouring effect, the expansion agent grouped the 256 phosphorylation patterns into 74 classes with identical rates, providing an exact reduction that was not used in the published study (Supplementary Notes S2.2 and S2.4, Table S5). M3 extracted the data and provided the measurement table.

In M4, the condition agent encoded the single experimental condition, and the observation agent translated the description of the measurement into an observation model. As the description states that the intensities are not comparable between time points and that the measurement error is assumed to be additive and normally distributed, with one standard deviation that scales with intensity at each time point, the observation agent defined a time-point-specific observable and noise formula:

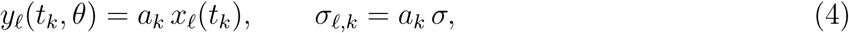

in which *x_ℓ_*denotes the concentration of CTDs with *ℓ* phosphorylated repeats, *a_k_* the scaling parameter of time point *t_k_*and *σ* a global noise parameter. This matches the observation and noise model of the published analysis. The module engine assembled one PEtab problem per model candidate, and the assembly verification agent audited them against the description and reported one concern: at 0 and 16 h only one species is populated, so with a scaling parameter per time point these spectra are fitted exactly by every candidate and contribute only zero residuals to the estimate of *σ*. The interpretation agent took this into account when judging the fits: it attributed about half of the AIC margins between the candidates to the shared *σ* rather than to the mechanisms.

Octomod then ran through M5 to M7, estimating all five model candidates with 100 starts each.

M7 selected the model combining neighbouring and global effects (8 parameters, AIC = *−*166.4), whereas the published analysis favoured the Neighbouring-effect model (7 parameters, AIC = *−*166.0). As the observation and noise models are the same, the Octomod and published analyses can be compared directly. Accordingly, Octomod identified without user feedback a model on par with the published model.

The interpretation agent reported that the selected model lagged behind the data at 1 h and ran ahead at 2 h, and recommended revision. The revision agent identified their shared assumption of constant kinase activity and proposed product inhibition. The revised model was fitted in a second round and selected over the round-one model (9 parameters, AIC = *−*172.8; Fig. 3E). Product inhibition slows the reaction, as ATP dependence does in the published model. The resulting model achieves an AIC lower by 6.8 than the published model and describes the data similarly well (Fig. 3C). Both models indicate cooperative phosphorylation, but differ in where the cooperativity acts. In the published model cooperativity is local: a phosphate on a neighbouring repeat increases the phosphorylation rate 2.3-fold. In the Octomod model the effects are globally positive and locally negative: every phosphate on the CTD increases the rate at every repeat 1.6-fold, while a neighbouring phosphate additionally scales the rate by 0.31, approximately halving it overall. As the run report notes, the mass spectrometry data do not resolve the positions of the phosphates and therefore cannot distinguish these mechanisms. The two models consequently predict different phosphate arrangements: among CTDs with two phosphorylated repeats at 1 h, the published model places them next to each other in 47 % of the CTDs, the Octomod model in 8 % (Fig. 3D).

In conclusion, from a description of the experiment, the workflow arrived at the published observation model, an exact reduction of the state space and a model that describes the data as well as the published one, with a different form of the cooperativity, which, given the data, cannot be resolved. Importantly, it even highlights limitations of the data concerning the identifiability of different mechanism.

### Octomod facilitates knowledge-guided model refinement

Having shown that Octomod can develop a model comparable to a published one without human guidance, we next explored how it can support model development in an interactive setup, where a user communicates with Octomod through the orchestrating agent. For this purpose, we consider the published bulk transcriptomic dataset from Xue et al. (2014), in particular, the data for GM-CSF-differentiated macrophages stimulated with IFN-*γ* or left untreated. We focused on four key response genes: the transcription factor IRF1, the signalling inhibitor SOCS1 and the chemokines CXCL9 and CXCL10. The curated panel comprised 324 measurements spanning baseline to 72 h. The initial research question asked: can a shared activating signal, together with gene-specific mRNA production and degradation, describe the dynamics of transcriptional response of human macrophages to IFN-*γ*, or are additional regulatory mechanisms required? We supplied this question and an initial set of proposed species and reactions to Octomod. The online literature search of the staging module (M0) was excluded and no additional online research by Octomod was allowed (Supplementary Note S3.2). The initial questions, resulting hypotheses, success criteria and development steps are detailed in the Supplementary Information (Supplementary Note S3).

In M1, Octomod constructed a five-state base model comprising a latent activator *A*, as well as transcripts for the four key response genes, each with basal and activator-induced production and first-order degradation (model 0, Fig. 4A; equations of models 0–5 in Supplementary Note S3.4.

**Figure 4:**
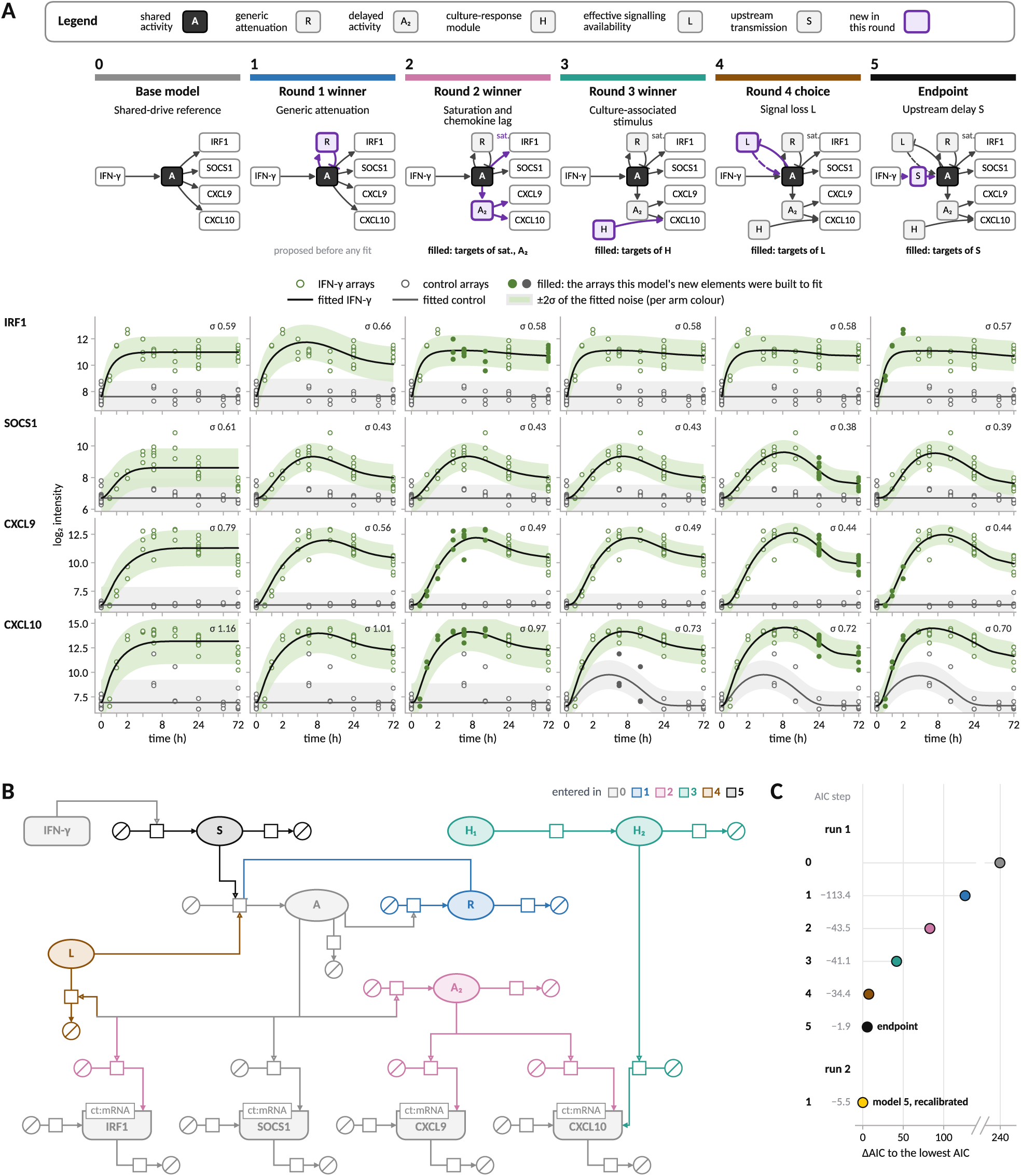
**Interactive development of a model for macrophage IFN-***γ* **response. A** The models along the refinement, from left to right: a base structure without feedback (model 0); generic activity-induced attenuation (model 1); saturating IRF1 induction and a delayed chemokine-driving activity (model 2); a CXCL10 culture-associated component (model 3); activity-dependent loss of signalling availability (model 4); and the endpoint with upstream signal-transmission stage (model 5). Purple outlines mark elements new relative to the preceding model. Below, stimulated (green) and untreated (grey) observations with the fitted means (black, stimulated; grey, untreated) and the fitted mean *±*2*σ_i_* (shading), where *σ_i_* is the estimated residual standard deviation of transcript *i*, shared across treatment arms. Filled circles mark the observations Octomod identified as guiding the extension from the last model. **B** SBGN process description of the endpoint model (model 5); colours indicate the model in which each element was introduced. **C** AIC of all models relative to the lowest AIC, including model 5 after reparameterisation in a second run; numbers give the AIC change relative to the preceding model.

Although *A* was not assigned a molecular identity, the construction agent suggested a possible link to STAT1, which transmits IFN-*γ* signals through phosphorylation-dependent nuclear translocation (Haspel et al., 1996).

The base model provided the starting point for iterative model development, with agents proposing revisions based on residual patterns and users guiding their selection. In the first round, the expansion agent proposed SOCS1-linked feedback, generic attenuation, or stimulus loss as possible extensions of the base model (M2). After data extraction (M3), PEtab assembly (M4) and model fitting (M5-M6), the evaluation (M7) favoured generic activity-induced attenuation represented by the repressor *R* based on goodness-of-fit, residual drifts and an AIC improvement of 113.4 relative to the base model (model 1, Fig. 4A; all model candidates in Supplementary Table S8. However, the success criteria were not yet met and a next round was recommended and agreed upon by us. The revision agent then identified persistent IRF1 misfit despite a bound-limited degradation rate and early overprediction followed by underprediction of chemokine expression. It proposed saturating IRF1 induction and a delayed chemokine-driving activity, *A*_2_, which we requested to evaluate jointly as well as separately. Their combination outperformed either extension alone, improving AIC by 43.5 (model 2, Fig. 4A), although the early IRF1 peak remained unresolved (Supplementary Fig. S2). Next, the revision agent highlighted the untreated CXCL10 increase at 6–12 h (Supplementary Fig. S2), which contradicted the assumption that the control is at steady-state, and proposed extensions to explain these measurements. From these, we selected an additional component driving CXCL10 production in both conditions for the next round’s expansion module. Evaluation showed an improvement in AIC by 41.1, mainly through a better fit to the controls (model 3, Fig. 4A). Remaining late SOCS1 and chemokine residuals and bound-limited attenuation led the orchestrating agent to suggest reopening the attenuation comparison. We decided to reopen this question and the revision agent proposed SOCS1-protein feedback and activity-dependent loss of signalling availability. Although an additional candidate with a condition-specific CXCL9 residual scale achieved the lowest AIC, the evaluation module attributed its gain primarily to tightly clustered control observations rather than improved time-dependent predictions (Supplementary Note S3.3). Informed by this assessment, we selected the activity-dependent model for further development (model 4, Fig. 4A). A final round introducing an upstream delay of IFN-*γ* signalling yielded only a marginal gain in AIC (model 5, Fig. 4A). After five rounds, residual variation was consistent with the observed variability among biological replicates. However, several parameters remained at their search bounds (Supplementary Table S9), leading the orchestrating agent to recommend a new Octomod run focusing on revising parameter bounds and reaction parameterisations rather than further structural extensions.

The second run supplied the model with a rescaled attenuator, revised saturation and amplitude parameterisation, fixed limiting missing fractions and wider selected bounds improving the AIC of the final model to 533.5 (Supplementary Note S3.5). The resulting mechanistic ODE model (Fig. 4B) consists of 11 state variables and 25 parameters. While most residual patterns were resolved by the workflow, Octomod also reported that the early IRF1 peak and donor-associated residuals suggested a poor fit, for a few datapoints (Supplementary Fig. S2). Since the revision agent could not identify further significant improvements, we decided to stop the model development. Thus, the cycle produced both interpretable extensions and explicit limits.

Although the retained components carry no molecular identity, each has a plausible counterpart in interferon biology. The activity-induced attenuator (model 1) is consistent with negative feedback on JAK–STAT signalling, for instance by SOCS1 (Liau et al., 2018; House et al., 2023), whose transcript may poorly reflect its inhibitory activity because SOCS1 is regulated post-transcriptionally (Gregorieff et al., 2000). The saturating IRF1 response, and the delayed chemokine drive (model 2), match the higher IFN-*γ* sensitivity of IRF1 compared with CXCL9 and CXCL10 (Naigles et al., 2023), and the declining signalling availability (model 4) resembles the IFN-*γ*-induced loss of IFNGR1 expression and responsiveness in myeloid cells (Crisler et al., 2019). The CXCL10 component acting in both arms (model 3) most likely reflects a technical culture response, and keeps it apart from IFN-*γ*-driven regulation.

In conclusion, Octomod supports interactive model development through substantive exchanges with the orchestrating agent. Its comprehensive assessment of model fits and residuals informed our discussions, allowing us to connect discrepancies with biological hypotheses and shape subsequent comparisons. These exchanges, conducted in natural language, prompted joint evaluation of extensions and renewed comparison of attenuation mechanisms, while the workflow generated model code and performed numerical evaluation. The complete process took a few hours (Supplementary Table S3), yielded a biologically interpretable model that can explain the data, including differences in response timing and dynamics unrelated to stimulation.

### Octomod constructs and compares models of cell–cell communication across cell types and stimuli

Having demonstrated Octomod on small models, we next assessed its scalability on a larger modelling problem involving intercellular communication. We considered data from Bodenmiller et al. (2012), who used mass cytometry to measure signalling dynamics in human peripheral blood mononuclear cells (PBMCs; Fig. 5A). Their study aimed, in part, to investigate communication between cell types. One example is the STAT response of lymphocytes to lipopolysaccharide (LPS). The lymphocytes’ response to IFN-*α* starts within minutes, but their response to LPS only after about an hour (Fig. 5B). Because lymphocytes express little or no LPS receptor, the authors proposed that this response could be mediated by IL-6 or other factors released by monocytes. This raises a mechanistic question: can the observed responses be explained by cell-intrinsic signalling alone, or do they require communication between cell types and, if so, which secreted factors are consistent with the data? To address this question, we selected six stimuli and an unstimulated reference. Considering multiple stimuli allows us to test whether a single model can coherently explain responses to different inputs, rather than fitting the LPS response in isolation. The other stimuli may also provide information about cell-type-specific signalling and crosstalk, helping constrain which mechanisms are shared across conditions. We gated five cell types (B cells, CD4^+^ and CD4*^−^* T cells, NK cells and monocytes) and summarised phosphorylated STAT1, STAT3 and STAT5 by their medians within each cell type, yielding 840 measurements. We modelled the five cell types in a shared medium, so that candidate models can represent communication through secreted factors, and compared their timing and cell-type-specific STAT responses. Together with the orchestrating agent, we formulated the project description and intent based on the dataset and original publication. We specified five candidate explanations from the literature, including models with monocyte-secreted factors and a delayed intracellular lymphocyte response without secreted factors. We asked the workflow to evaluate these candidates and identify additional hypotheses, then compared its analysis with agent-only analyses of the same problem.

**Figure 5:**
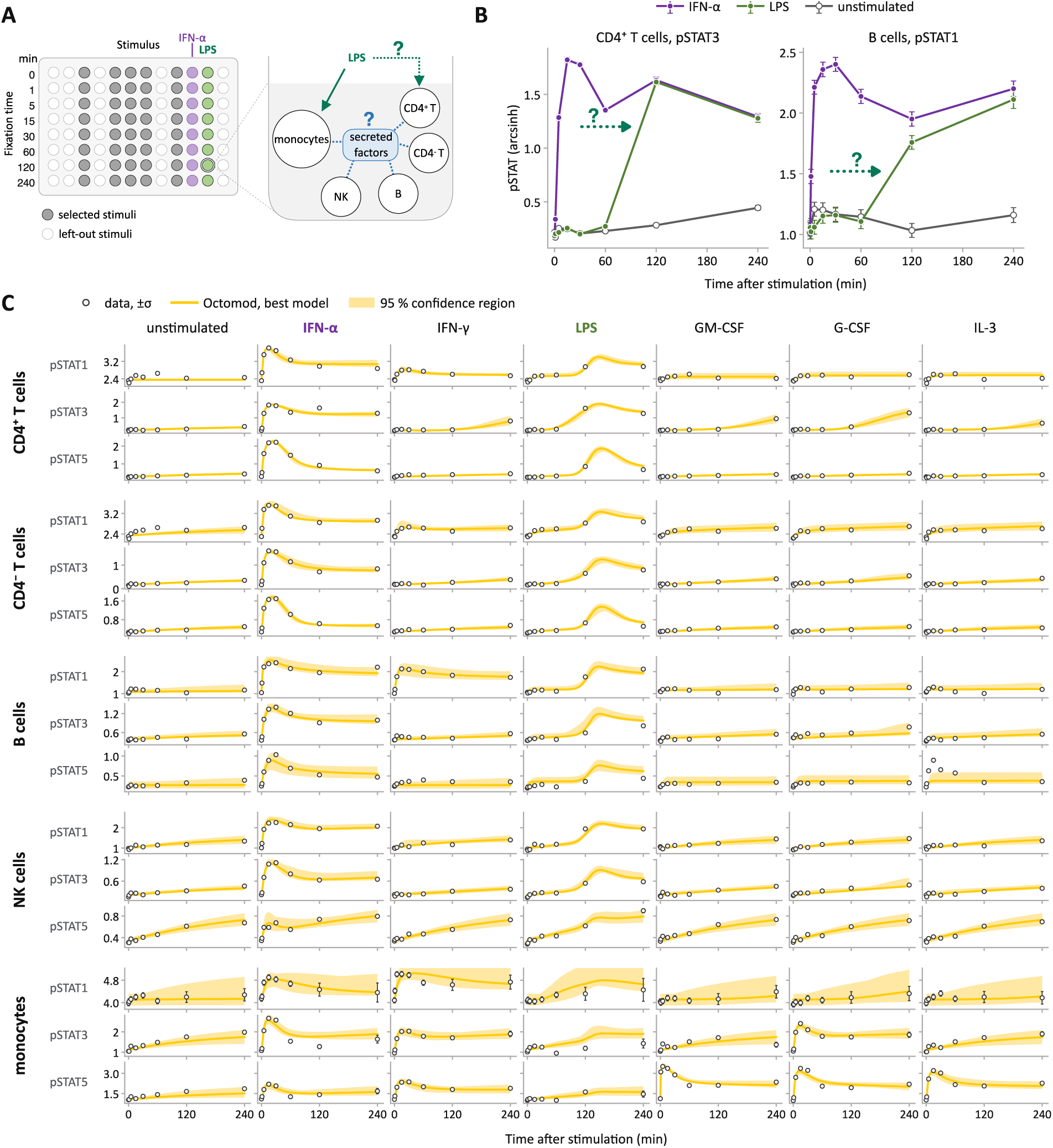
PBMC data and the fit of the Octomod best model. **A** One plate with the stimuli in columns and the fixation times in rows, and one well with the five cell types in a shared medium. Under LPS, lymphocytes may respond to factors secreted by the monocytes (blue) or through a delayed response of their own (green dotted). **B** pSTAT3 in CD4^+^ T cells and pSTAT1 in B cells under IFN-*α*, LPS and unstimulated; error bars show the noise level *σ*. The arrow marks the delay of the LPS response relative to the IFN-*α* response. **C** All 840 measurements (circles, *±σ*) with the simulation of the Octomod best model (yellow) and the envelope of its simulations over the joint 95 % confidence region of the parameters (shading).

In M0, the knowledge agent surveyed the literature in two calls and gathered evidence from 44 sources for the reactions and facts it proposed, which the knowledge verification agent checked. The resulting knowledge file held 73 reactions and 25 facts, each with its sources and evidence grade (Supplementary Tables S12 and S13). The list of facts included information that (1) LPS, GM-CSF, G-CSF and IL-3 receptors are restricted to myeloid cells, (2) type-I interferon receptors are expressed on all cell types, and (3) the IFN-*γ* receptor is high on monocytes, moderate on B cells and low on T cells. We accepted the suggested floor, “low”. The suggestion was set by the lowest-graded essential reaction, the inactivation of the type-I interferon receptor, for which the knowledge agent found evidence only from human cell lines (Kumar et al., 2003).

In M1, the construction agent proposed a base model covering all five cell types in the shared-medium experiment (64 species, 104 reactions and 109 estimated parameters; Fig. 6A, grey). The base model included cell-type-specific cellular processes but no explicit communication between cell types. Each cell type had receptors for type-I interferon, IFN-*γ* and IL-6, as well as STAT1, STAT3 and STAT5, their phosphorylated forms, and SOCS feedback. The presence of receptors varied by cell type according to the literature information on receptor expression levels: receptors for LPS, GM-CSF, G-CSF, and IL-3 were included only for monocytes, as specified in the knowledge file. A drift term represented the slow rise in all readouts over the four-hour experiment, a rise also seen in the unstimulated well (Supplementary Note S4.3, Fig. S4). This absence of communication applied only to the base model; the expanded candidates introduced communication through factors secreted by monocytes.

**Figure 6:**
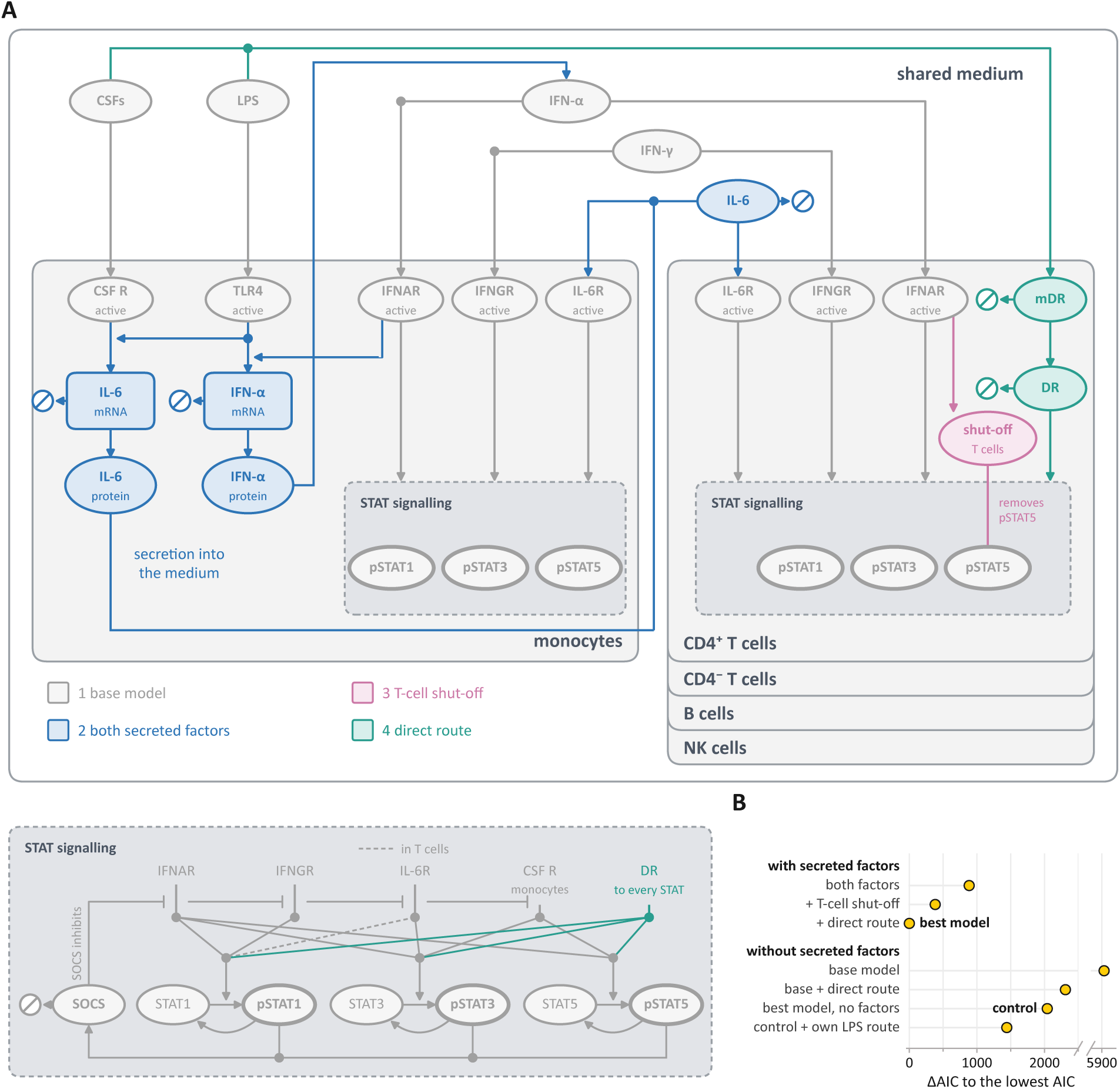
The best model and the model selection of the PBMC case. **A** The best model, drawn as one well. The shared medium at the top carries the stimuli and the two secreted factors, the monocytes and the four lymphocyte cell types below it carry their receptors, and the STAT signalling that is the same in every cell is drawn once at the bottom. The monocytes secrete a type-I interferon, drawn as IFN-*α*, and IL-6. The direct route induces in every lymphocyte cell type a hypothetical transcript (mDR) and protein (DR), and the T-cell shut-off removes pSTAT5. Each element is coloured by the model that added it (legend). Dashed couplings exist in T cells only, the measured pSTATs are in bold, and Ø marks degradation or clearance. CSFs are GM-CSF, G-CSF and IL-3. **B** ΔAIC of the seven Octomod models to the lowest AIC, the best model, grouped by whether the model contains secreted factors. Each “+” adds one mechanism to the row above, and the control is the best model with the secreted factors removed.

In M2, the expansion agent proposed four expansions to the base model: monocyte secretion of type-I interferon, secretion of IL-6, secretion of both factors, or both factors with basal secretion in place of the drift. The construction verification agent audited the five candidates and flagged one concern: the candidate with IL-6 secretion alone had no route to STAT1 in B cells and therefore could not produce their late pSTAT1 response. At the interaction point that followed, we chose all five candidates for estimation, since the concern affected only how the two candidates with one secreted factor were compared, not what was fitted.

After data extraction (M3), PEtab assembly (M4), and model fitting (M5, M6) with 100 starts per candidate, evaluation (M7) selected the model with both secreted factors over all other model candidates. This model described most time courses, but systematic misfits remained (Supplementary Note S4.4). In two further rounds, the revision agent proposed mechanisms for these misfits, among them a STAT-specific shut-off of the response in CD4^+^ and CD4*^−^* T cells, production of IL-6 triggered by type-I interferon, a direct response of B cells to IL-3 and a direct route from LPS and the colony-stimulating factors (CSFs; GM-CSF, G-CSF and IL-3) into the lymphocytes. We chose nine of them for estimation, in addition to the five candidates of the first round; each round’s candidates were fitted and evaluated before the next round was proposed. The model with the lowest AIC, called the best model in the following, contained both secreted factors from the monocytes, the shut-off in the T cells and the direct route (Fig. 6A).

To test whether communication is required, we requested two further model candidates in two more rounds: a control, the best model with the production of both secreted factors switched off, and a version of it in which LPS acts on each lymphocyte type through a route of its own, a hypothetical receptor with its own induced transcript and protein. In total, 17 models were built and fitted (Supplementary Note S4.4, Table S14). Of these, four models did not contain secreted factors: the base model, the base model with the direct route, the control and the control with its own LPS route (Fig. 6B). Among them, the control with its own LPS route fitted best, but its AIC was still much higher than that of the best model (ΔAIC = 1441). The models with secreted factors reproduced the absence of a response during the first hour and the rise between 60 and 120 minutes (Fig. 5C, LPS), which suggests that the delay is caused by the production of the secreted factors. In contrast, the models without secreted factors, in which the lymphocytes respond through a direct route, responded from the start and reached the late level too slowly (Supplementary Fig. S5). Thus, all Octomod model candidates without communication between the cell types are rejected by the data. Overall, the best model described the time courses of all readouts under all seven conditions (Fig. 5C).

To assess the benefits of using Octomod for biologically grounded model development, we compared its analysis with independent runs of Claude Code and Codex on the same description and data, with online literature access but without user intervention (Supplementary Note S4.6, Table S15). Codex did not produce a model that described the data, due to an unprompted choice by the agent to strictly limit the number of parameter estimation starts and steps to save computational resources. Claude Code fitted 32 models in 15 h, using its own code rather than SBML and PEtab, and selected a model with an AIC 770 lower than that of the best AIC Octomod model, despite having 18 additional parameters (Supplementary Fig. S7). We verified its objective values and independently re-implemented the selected model (Supplementary Fig. S8). The two analyses agreed on most findings but differed on whether communication is required (Supplementary Note S4.6, Table S16). However, Claude Code’s selected model represented lymphocyte responses to LPS and the CSFs through free delayed chains, without explicit STAT species, secreted factors or receptors for these stimuli (Supplementary Fig. S6). Because it did not account for the biological basis of these responses, its lower AIC is unsurprising: the free delayed chains could fit the observed delays without having to explain them through biologically specified mechanisms. Thus, the comparison illustrates a trade-off between fit and biological grounding, rather than evidence that the agent-only model provides a better mechanistic explanation.

Together, these results show that Octomod can construct and compare mechanistically specified models of cell–cell communication across multiple cell types. In this case, the best-supported model linked monocyte signalling to lymphocyte responses through secreted factors, while the alternative of a free delayed response could reproduce the data without identifying a biological mechanism. The comparison therefore highlights how explicit biological structure can make a model useful for investigating communication, beyond achieving a close fit. Next, we tested Octomod on unpublished lipid data, where few established models were available.

### Literature-informed hypothesis testing refines a hand-built model of new, unpublished lipid data

The previous applications demonstrated the versatility of Octomod across different modelling problems. However, although tailored models were unavailable for the second and third applications, their datasets had already been published and might have been represented in LLM training data. We therefore next evaluated Octomod on a newly generated, unpublished dataset that could not have been encountered during training. This application is also used to evaluate Octomod’s ability to assess how well the characteristics of biological processes can be informed based on measurements, how strongly they constrain those processes, and which measurements contribute useful information to the analysis.

We generated pulse-chase lipidomics data on sphingolipid metabolism in human SH-SY5Y neuroblastoma cells. Cells were pulsed for 60 min with alkyne-labelled sphinganine (Sph) and palmitate (Pal), then chased for 180 min. The fraction of the initial label in 11 lipid classes was measured at six time points, with three replicates per time point (see Methods). Sphingolipids are characterized by a sphingoid-base backbone and have structural and signalling roles, with links to processes including apoptosis and autophagy (Hannun and Obeid, 2018; Quinville et al., 2021). Although mechanistic models of sphingolipid metabolism have been published (Gupta et al., 2011; Wronowska et al., 2015; Savoglidis et al., 2016), each is tailored to the species, inputs and organism of its original study and could not be directly applied to these data.

We provided Octomod with an SBML draft model (14 species, 21 reactions, 11 measured species), the data, and descriptions of the experiment and modelling objective. We asked whether the model captured the processes relevant to interpreting the labelled time courses, whether its reactions were supported by the literature, the data or both, and whether relevant processes were missing. We also asked which reactions and species could be removed without compromising the model’s mechanistic scope, and which measurements are not informative (e.g., because the species are located at the boundary of the modelled pathway and unobserved outflows could compensate for their dynamics). We addressed these questions over six consecutive runs. In each run, Octomod evaluated one or more structural hypotheses against the data, and the modeller reviewed the results and revised the model with the workflow. Each run began from the model refined in the preceding run, allowing the analysis to build on earlier findings. We compared alternative model structures using the Akaike information criterion (AIC).

The orchestrating agent provided a conversational interface for examining the model’s scope and discussing how well the available data could inform its processes. Across the six runs, we used this interaction to formulate and test structural hypotheses, and to remove observables. The following five cases illustrate these decisions (Fig. 7); together, they span all six runs.

**Figure 7:**
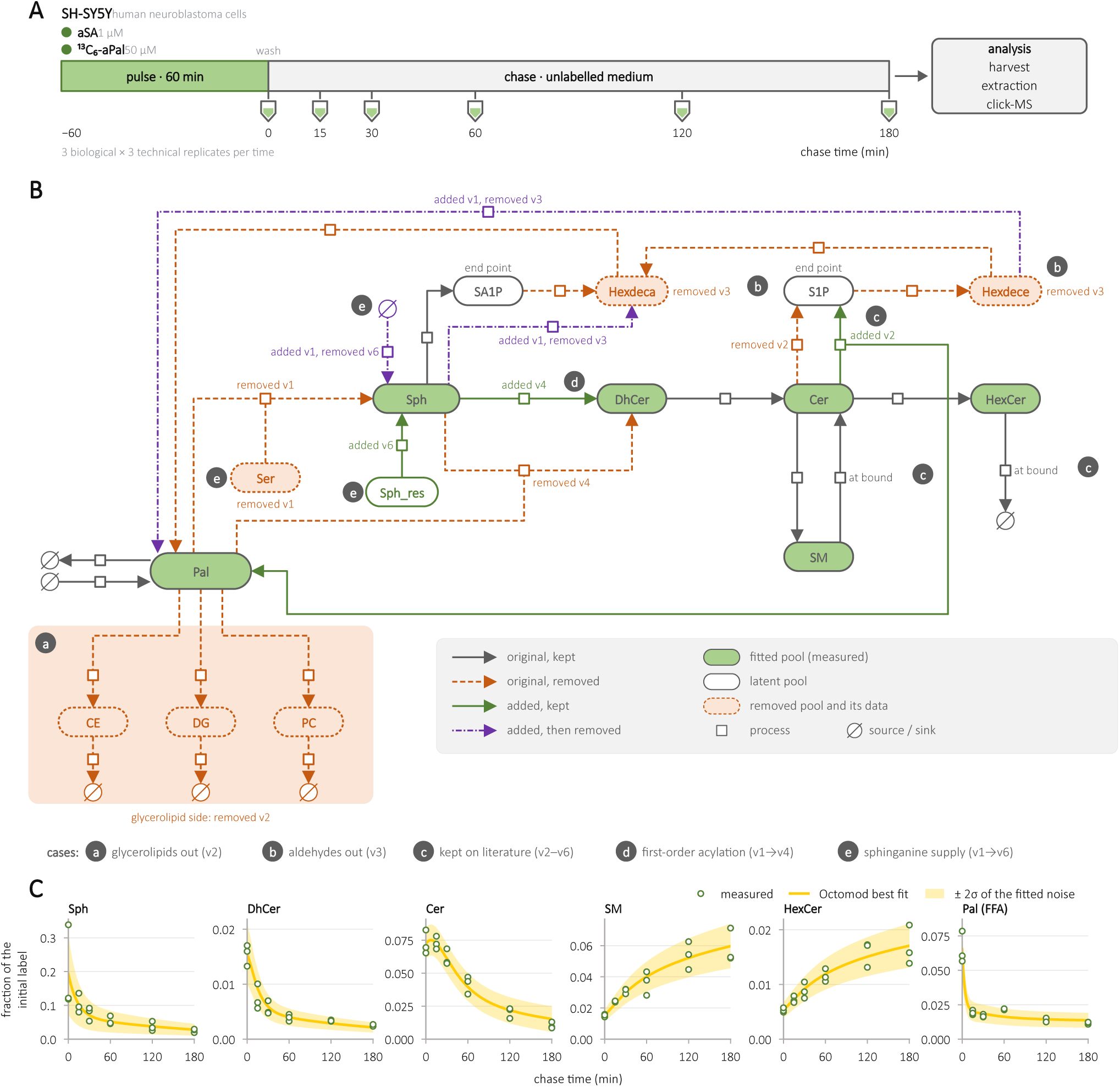
Development of the sphingolipid model across six consecutive runs (v1–v6). (A) Pulse-chase experiment: SH-SY5Y cells were labelled for 60 min with alkyne sphinganine (aSA) and ^13^C_6_-alkyne palmitate (^13^C_6_-aPal) and harvested at six chase times. (B) Process description map (SBGN) of the supplied hand-built model (14 species, 21 reactions) and of every reaction added during the series; the final model (v6) has 9 species and 11 reactions. Arc styles give the fate of each reaction (key), labels the run in which it was added or removed. “At bound”: rate estimated at its lower bound, so that only an upper limit is determined. Green nodes are the six fitted lipid classes, white nodes unmeasured pools. Badges (a)–(e) mark the five cases discussed in the text. (C) Final model (v6): measured fraction of the initial label (circles; three biological replicates), best fit (line) and *±*2*σ* of the fitted noise model (band).

#### (a) Identification of measurements for limited or no information

Initially, phosphatidyl-choline (PC), diacylglycerol (DG) and cholesteryl ester (CE), all of which were measured, were included in the supplied model. Each was represented by the same pathway structure, Pal *→* Pool *→ Ø*. Although these reactions link palmitate (Pal) to the three pools, they provide only a generic representation of glycerolipid metabolism, with parallel in- and outflows. Modelling the underlying processes in detail would have made the model overly complex and difficult to identify from the available data. The orchestrating agent concluded that measurements did not provide information about the sphingolipid metabolism. We absorbed the pools into the model’s existing Pal outflow, Pal *→ Ø*, and set the measurements aside. Because the workflow requires all supplied measurements to be included in a run, excluding measurements required us to restage the problem and start a new run, documenting this decision.

#### (b) Identifying an unresolved label flux

The experiment used distinguishable labels for sphinganine (Sph) and palmitate (Pal). The low-abundance fatty aldehydes hexadecenal (Hexdece) and hexadecanal (Hexdeca) carried predominantly the Pal label (Supplementary Fig. S9B), which could not be explained by sphingolipid degradation: this process preserves the sphingoid back-bone derived from Sph, and the Sph carried no Pal label. Neither the knowledge module nor a literature search by the orchestrating agent identified a plausible route within the model’s scope. As possible explanations involved processes outside that scope, uncertainty in assigning low-abundance species, or separate modelling of the two labels, we set the aldehyde measurements aside and removed the aldehydes from the model. This focused the analysis on processes that could be assessed using the Sph label, while leaving the route by which Pal label enters the alde-hydes unresolved. Sphingosine-1-phosphate (S1P) and sphinganine-1-phosphate (SA1P) therefore remained as model endpoints: they represent the next known steps of ceramide degradation and sphinganine phosphorylation, but function as sinks in the model.

#### (c) Literature evidence informs decision in case of initially inconclusive statistical test

The knowledge module suggested modifying the reaction Cer *→* S1P to include the release of palmitate (Pal), based on a curated Reactome entry graded high (i.e., Cer *→* S1P + Pal). The first hypothesis test was inconclusive (ΔAIC = *−*1.0), but we retained the reaction because of its literature support. In a later run, we tested its removal again; this was disfavoured (ΔAIC = +5.6). Thus, literature evidence complemented the data in model selection, and repeated hypothesis testing strengthened support for including the Pal release.

#### (d) Revising acylation to match the observed label pattern

The supplied model represented acylation as the bimolecular reaction Sph + Pal *→* DhCer. With the two labels represented together, this formulation couples the formation of labelled dihydroceramide (DhCer) to consumption of labelled Pal, implying double-labelled DhCer. Yet the measured sphingolipids carried almost exclusively the Sph label, and double-labelled species could not be observed. The construction agent flagged this issue in the first run and proposed an unlabelled acyl-CoA donor; we deferred that change. Later, the interpretation agent identified this coupling as a possible contributor to the need for constant Sph and Pal influxes (removal: ΔAIC = +40.1). We then tested a first-order acylation, Sph *→* DhCer, which consumes no labelled Pal. This change was supported (ΔAIC = *−*6.5), almost entirely through an improved fit to the Pal data, while both influxes remained necessary (removal: ΔAIC = +39.7). The test therefore favoured the representation more consistent with the observed label pattern, although it was biochemically less complete.

#### (e) Replacing constant sphinganine import with a depleting reservoir

The final model also changed how Sph was supplied. The supplied model included de novo synthesis (Pal + Ser *→* Sph), which we replaced with a constant Sph import because the Sph carried no Pal label (Supplementary Fig. S9A). The import was required (removal: ΔAIC = +10.3), but its constant rate maintained Sph above the measured levels late in the chase and consequently kept Cer elevated. The interpretation agent ranked declining Sph supply as the most likely explanation for this misfit. Replacing the constant import with a depleting, unmeasured Sph reservoir was supported (ΔAIC = *−*7.5), although Cer remained overestimated late in the chase.

Cases (d) and (e) both arose from representing the Sph and Pal labels as a single label, a simplification fixed in the modelling objective from the start. The workflow worked within this scope, but repeatedly identified separate modelling of the two labels as the natural scope extension of it, which would also allow testing one of the possible explanations in case (b).

In summary, on newly generated, unpublished data, Octomod supported the iterative evaluation of biological processes and model scope. Across six runs, we refined a supplied model with 14 species and 21 reactions, yielding a model with 9 species and 11 reactions. The final model was fitted to 6 of the 11 measured lipid classes (108 data points): the glycerolipid pools (case a) and the fatty aldehydes (case b) were set aside. The six runs took about 43 h of wall time, including about 21 h spent at human interaction points.

## Discussion

Developing mechanistic ODE models is not linear; it is an iterative process that spans defining the model structure, assessing the scope, integrating data, estimating parameters, and analysing and refining the model and potentially its scope. Each step has its own requirements and degrees of freedom, and decisions made at one point propagate to the others and to the final model (Kitano, 2002). To inform, guide and accelerate this modelling process and its associated decisions, we present Octomod, a proof-of-principle that couples trusted software modules with LLM-based assistance.

We showed that Octomod constructs plausible models from small to large scale. Without access to the published model, it built a model from the same data and setting that was comparable to the human-constructed counterpart. For data without an existing model, it proposed new models that provided a plausible description for the potentially underlying biological mechanisms. Our application to lipid dynamics and metabolism illustrates how valuable knowledge accumulates along this iterative process: Octomod identified measurements that did not inform the modelled pathway. Because the workflow does not let measurement data be silently ignored, they were set aside only in a restaged run with the remaining data. Yet, when the model was simplified, two named species became sinks of the model, a subtle change in interpretation that the interactive review steps of Octomod make explicit. Such cases show that the human remains essential to understand, question and redirect the system, and that iterative refinement is central to the approach.

In designing Octomod, we deliberately constrained the LLM’s freedom. Deterministic module engines enforce established, validated software and standards (Schmiester et al., 2021; Keating et al., 2020; Fröhlich et al., 2021; Schalte et al., 2023), engineered agents and checks operate within these modules, and the user acts as a human gate who decides and steers. The iterative workflow thus gains trust through constraints: the LLM contributes flexibility and domain reasoning at massive scope, while critical computations are performed by single tools with reproducible behaviour. Recent work suggests that such constraints may not be required for frontier LLMs to contribute to model building (de Pomereu et al., 2026; Shojaee et al., 2025). We therefore compared Octomod with two agent-only analyses, by Codex and by Claude Code, given structured prompting and the same project description and data. Codex, although comparing models in a reasonable way, limited itself unprompted to very few iterations during model fitting and produced no model that described the data. Claude Code, judged by goodness of fit, yielded a better model, albeit one that represented the lymphocyte responses by free delayed chains without a biological mechanism. Moreover, reconstructing what the agent-only analyses had done required substantial post-hoc inspection. Both the Codex and the Claude Code analyses, however, were run only once and were not instructed to use the standards and tools that would have helped them. The comparison is there-fore not exhaustive, and benchmarking across levels of structure is needed. Even so, constraining the search space improves traceability, possibly at the cost of excluding solutions outside it, but it also keeps the models mechanistic, with every reaction standing for a biological process and carrying its origin and evidence grade. Octomod is organised into modules whose deterministic engines can be exchanged and versioned independently, so every result can be traced to the software that produced it, and the variation of the LLM enters only through the proposals of the agents and the decisions of the user, both held in the run record. Conversely, the agent-only approach could be refined, for instance by prescribing an optimiser, requiring PEtab output, or enforcing specific checks. Each such refinement adds structure and gradually turns the agent-only approach into a workflow. GEMOT (de Pomereu et al., 2026) is such an intermediate point: it builds on the same standards and tools as Octomod (SBML, PEtab, AMICI, and pyPESTO), keeps the fitting and scoring of every hypothesis outside the LLM, and records every model version and the full trace of the run. The loop itself, however, is run by an LLM lead that decides what to query and which hypotheses to build and revise, without a human in the loop. The question is therefore not workflow versus no workflow, but how much structure is needed, and where, to obtain trustworthy results.

Although our comparison of Octomod with the agent-only analyses does not fully answer this question, it highlights several relevant factors and practical difficulties in evaluating the benefits of structure. For example, given identical prompts, the two agent-only analyses produced different results and used different tools and routes, indicating high variance, which has also been quantified for other use cases (Cui and Alexander, 2026). Assessing this variance systematically would require repeated runs and varied settings, and assessing bias would require problems no LLM has seen. Both were beyond the scope of this study. Still, we expect the workflow to reduce variance, although the human in the loop remains a source of variation and bias in itself. Exact reproducibility is out of reach for any LLM-based method: LLM outputs differ between runs, and commercial frontier models change behaviour even within the same version, exemplified by varying enforced restrictions. Further, it is unclear how much an LLM agent draws on information it was trained on but was not explicitly given, even when its context and environment are carefully controlled. Thus, alongside search-time data contamination (Han et al., 2025), this is a second avenue for evaluation bias when comparing against published models as ground truth. Hence, the traceability of modelling decisions, including, e.g., the choice of observable functions, the evaluated model versions and the parameter estimation setup, remains the basis for judging their suitability and validity.

Consequently, describing the data alongside software versions and open-source code is not sufficient. Standards such as SBML and PEtab capture the final model and estimation problem, but not how they came about. Until recently, no modelling tool or workflow recorded this derivation. GEMOT (de Pomereu et al., 2026) does so in its trace, and so does the run record introduced here. Still, the field needs agreed-upon objects that record every step of model development to enable exchangeability, reproducibility, and integrity. The run record, which captures the complete model and its derivation, can serve as a blueprint for such a standard. Together with trusted APIs between agents and established tools, this would let the field harness the power of agentic modelling demonstrated here while keeping the process verifiable.

## Data availability

The CTD phosphorylation data are from Callenbach et al. (2026), the macrophage transcript data from GEO (GSE46903, Xue et al. 2014), and the PBMC mass cytometry data from Bodenmiller et al. (2012). The lipidomics data were generated for this study. They are available in a Zenodo record (https://doi.org/10.5281/zenodo.24147167), together with the processed data and the run records of all applications, the models in SBML and PEtab, and the re-estimation results.

## Code availability

Octomod is available at https://github.com/ICB-DCM/Octomod under the BSD-3-Clause licence. The analysis scripts and the scripts behind the figures are in the Zenodo record given under Data availability.

## Acknowledgements

This work was supported by the Deutsche Forschungsgemeinschaft (DFG, German Research Foundation) under Germany’s Excellence Strategy (EXC 2047—390685813, EXC 2151—390873048) and under the project IDs 432325352 – SFB 1454 and 450149205 – TRR 333, by the European Union via ERC grant INTEGRATE (GA 101126146), the German Federal Ministry of Research, Technology and Space (BMFTR) under the CompLS program (InterpretTME, grant no. 031L0308B), and by the University of Bonn (via the Schlegel Professorship of J.H.). The funders played no role in study design, data collection, analysis and interpretation of data, or the writing of this manuscript. The authors gratefully acknowledge access to the Marvin cluster of the University of Bonn.

## Author contributions

JH, DD, YL, and DP conceptualised the project. JH acquired financial support. DD and YL designed and implemented Octomod, with contributions from DP, VW, and PJJ. DD conducted the CTD and PBMC applications, VW the macrophage application, PJJ the sphingolipid application, and MH and SM the agent-only analyses. CT and LS evaluated the biological plausibility of the models. PRMD and CT generated the lipidomics data. DD, YL, VW, PJJ, SM, LS, CAG, DP, and JH wrote the manuscript and prepared the figures. All authors reviewed and commented on the manuscript and approved the final version.

## AI use statement

Generative AI was used in the evaluated system and in preparing this work. In the reported applications, the agents of Octomod are calls of Claude Code (Anthropic), with the models and settings given in the Methods and Supplementary Table S3, and the agent-only analyses were run with Claude Code and Codex (OpenAI) as described in the Methods. During the research, Claude Code helped implement Octomod and the analysis and figure scripts, and was used to read the run records and to check the reported values against them. Claude drafted the Supplementary Notes and figure captions from the run records and suggested edits to the main text, which the authors revised. AI was not used to generate measurement data. The authors checked all code and values, directed the work, and retain responsibility for its claims, references, code, and final text.

## References

Adusumilli, R. and Mallick, P. (2017). Data conversion with ProteoWizard msConvert. In Comai, L., Katz, J. E., and Mallick, P., editors, Proteomics: Methods and Protocols, volume 1550 of Methods in Molecular Biology, pages 339–368. Humana Press, New York, NY.

Akaike, H. (1974). A new look at the statistical model identification. IEEE Transactions on Automatic Control, 19(6):716–723.

Anthropic (2026). Claude Code. https://claude.com/claude-code. Software.

Arakane, K., Imoto, H., Ormersbach, F., and Okada, M. (2024). Extending BioMASS to construct mathematical models from external knowledge. Bioinformatics Advances, 4(1):vbae042.

Bodenmiller, B., Zunder, E. R., Finck, R., Chen, T. J., Savig, E. S., Bruggner, R. V., Simonds, E. F., Bendall, S. C., Sachs, K., Krutzik, P. O., and Nolan, G. P. (2012). Multiplexed mass cytometry profiling of cellular states perturbed by small-molecule regulators. Nature Biotechnology, 30(9):858–867.

Brunton, S. L., Proctor, J. L., and Kutz, J. N. (2016). Discovering governing equations from data by sparse identification of nonlinear dynamical systems. Proceedings of the National Academy of Sciences, 113(15):3932–3937.

Burnham, K. P. and Anderson, D. R. (2002). Model selection and multimodel inference: A practical information-theoretic approach. Springer, New York, 2 edition.

Callenbach, A., Dorešić, D., Düster, R., Nakonecnij, V., Dudkin, E., Geyer, M., and Hasenauer, J. (2026). Quantitative modelling of P-TEFb mediated CTD phosphorylation identifies local cooperativity. PLOS Computational Biology, 22(7):e1014531.

Collier, J. R., Monk, N. A. M., Maini, P. K., and Lewis, J. H. (1996). Pattern formation by lateral inhibition with feedback: a mathematical model of Delta-Notch intercellular signalling. Journal of Theoretical Biology, 183(4):429–446.

Crisler, W. J., Eshleman, E. M., and Lenz, L. L. (2019). Ligand-induced IFNGR1 down-regulation calibrates myeloid cell IFN*γ* responsiveness. Life Science Alliance, 2(5):e201900447.

Cui, J. and Alexander, R. (2026). Same prompt, different outcomes: Evaluating the reproducibility of data analysis by LLMs. *arXiv*. Preprint, arXiv:2602.14349.

Czudnochowski, N., Bösken, C. A., and Geyer, M. (2012). Serine-7 but not serine-5 phosphorylation primes RNA polymerase II CTD for P-TEFb recognition. Nature Communications, 3(1):842.

d’Ascoli, S., Becker, S., Schwaller, P., Mathis, A., and Kilbertus, N. (2024). ODEFormer: Symbolic regression of dynamical systems with transformers. In International Conference on Learning Representations.

de Pomereu, T., Snelling, B., and Fröhlich, F. (2026). GEMOT: Towards mechanistic world models for biology. *bioRxiv*. Preprint, doi:10.64898/2026.09.26.754619.

Fröhlich, F. and Sorger, P. K. (2022). Fides: Reliable trust-region optimization for parameter estimation of ordinary differential equation models. PLOS Computational Biology, 18(7):e1010322.

Fröhlich, F., Weindl, D., Schälte, Y., Pathirana, D., Paszkowski, L., Lines, G. T., Stapor, P., and Hasenauer, J. (2021). AMICI: high-performance sensitivity analysis for large ordinary differential equation models. Bioinformatics, 37(20):3676–3677.

Gregorieff, A., Pyronnet, S., Sonenberg, N., and Veillette, A. (2000). Regulation of SOCS-1 expression by translational repression. Journal of Biological Chemistry, 275(28):21596–21604.

Grein, S., Penas, D. R., Weindl, D., Lakrisenko, P., Banga, J. R., and Hasenauer, J. (2026). Largescale analysis of optimisation methods for parameter estimation problems in the life sciences. *bioRxiv*. Preprint, doi:10.64898/2026.07.11.737731.

Gupta, S., Maurya, M. R., Merrill, Jr., A. H., Glass, C. K., and Subramaniam, S. (2011). Integration of lipidomics and transcriptomics data towards a systems biology model of sphingolipid metabolism. BMC Systems Biology, 5(1):26.

Gyori, B. M., Bachman, J. A., Subramanian, K., Muhlich, J. L., Galescu, L., and Sorger, P. K. (2017). From word models to executable models of signaling networks using automated assembly. Molecular Systems Biology, 13(11):954.

Han, Z., Mankikar, M., Michael, J., and Wang, Z. (2025). Search-time data contamination. *arXiv*. Preprint, arXiv:2508.13180.

Hannun, Y. A. and Obeid, L. M. (2018). Sphingolipids and their metabolism in physiology and disease. Nature Reviews Molecular Cell Biology, 19(3):175–191.

Hart, Y., Reich-Zeliger, S., Antebi, Y. E., Zaretsky, I., Mayo, A. E., Alon, U., and Friedman, N. (2014). Paradoxical signaling by a secreted molecule leads to homeostasis of cell levels. Cell, 158(5):1022–1032.

Hasenauer, J., Jagiella, N., Hross, S., and Theis, F. J. (2015). Data-driven modelling of biological multi-scale processes. Journal of Coupled Systems and Multiscale Dynamics, 3(2):101–121.

Haspel, R. L., Salditt-Georgieff, M., and Darnell, Jr., J. E. (1996). The rapid inactivation of nuclear tyrosine phosphorylated Stat1 depends upon a protein tyrosine phosphatase. The EMBO Journal, 15(22):6262–6268.

Herzog, R., Schuhmann, K., Schwudke, D., Sampaio, J. L., Bornstein, S. R., Schroeder, M., and Shevchenko, A. (2012). LipidXplorer: A software for consensual cross-platform lipidomics. PLOS ONE, 7(1):e29851.

Hoops, S., Sahle, S., Gauges, R., Lee, C., Pahle, J., Simus, N., Singhal, M., Xu, L., Mendes, P., and Kummer, U. (2006). COPASI—a COmplex PAthway SImulator. Bioinformatics, 22(24):3067– 3074.

House, I. G., Derrick, E. B., Sek, K., Chen, A. X. Y., Li, J., Lai, J., Todd, K. L., Munoz, I., Michie, J., Chan, C. W., Huang, Y.-K., Chan, J. D., Petley, E. V., Tong, J., Nguyen, D., Engel, S., Savas, P., Hogg, S. J., Vervoort, S. J., Kearney, C. J., Burr, M. L., Lam, E. Y. N., Gilan, O., Bedoui, S., Johnstone, R. W., Dawson, M. A., Loi, S., Darcy, P. K., and Beavis, P. A. (2023). CRISPR-Cas9 screening identifies an IRF1-SOCS1-mediated negative feedback loop that limits CXCL9 expression and antitumor immunity. Cell Reports, 42(8):113014.

Keating, S. M., Waltemath, D., König, M., et al. (2020). SBML Level 3: an extensible format for the exchange and reuse of biological models. Molecular Systems Biology, 16(8):e9110.

Kitano, H. (2002). Systems biology: a brief overview. Science, 295(5560):1662–1664.

Krongauz, D., Zulti, A., Segal, E., and Lazebnik, T. (2026). Automatic ordinary differential equations discovery for biological systems using large language model powered agentic system. *arXiv*. Preprint, arXiv:2607.13608.

Kumar, K. G. S., Tang, W., Ravindranath, A. K., Clark, W. A., Croze, E., and Fuchs, S. Y. (2003). SCF^HOS^ ubiquitin ligase mediates the ligand-induced down-regulation of the interferon-*α* receptor. The EMBO Journal, 22(20):5480–5490.

Lakrisenko, P., Pathirana, D., Philipps, M., et al. (2026). Benchmarking-Initiative/Benchmark-Models-PEtab: Benchmark Models PEtab v2026.01.24. Zenodo, doi:10.5281/zenodo.18356607.

Liau, N. P. D., Laktyushin, A., Lucet, I. S., Murphy, J. M., Yao, S., Whitlock, E., Callaghan, K., Nicola, N. A., Kershaw, N. J., and Babon, J. J. (2018). The molecular basis of JAK/STAT inhibition by SOCS1. Nature Communications, 9(1):1558.

Naigles, B., Narla, A. V., Soroczynski, J., Tsimring, L. S., and Hao, N. (2023). Quantifying dynamic pro-inflammatory gene expression and heterogeneity in single macrophage cells. Journal of Biological Chemistry, 299(10):105230.

Noori, A., Fishman, N., Fang, A., Fesser, L., and Zitnik, M. (2026). World models for biomedicine. Cell, 189(19):5845–5869.

Pathirana, D., Bergmann, F. T., Doresic, D., Lakrisenko, P., Persson, S., Neubrand, N., Timmer, J., Kreutz, C., Binder, H., Cvijovic, M., Weindl, D., Hasenauer, J., and the PEtab Select community (2026). PEtab Select: Specification standard and supporting software for automated model selection. PLOS Computational Biology, 22(9):e1014774.

Persson, S., Fröhlich, F., Grein, S., Loman, T., Ognissanti, D., Hasselgren, V., Hasenauer, J., and Cvijovic, M. (2025). PEtab.jl: advancing the efficiency and utility of dynamic modelling. Bioinformatics, 41(9):btaf497.

Quinville, B. M., Deschenes, N. M., Ryckman, A. E., and Walia, J. S. (2021). A comprehensive review: Sphingolipid metabolism and implications of disruption in sphingolipid homeostasis. International Journal of Molecular Sciences, 22(11):5793.

Raue, A., Kreutz, C., Maiwald, T., Bachmann, J., Schilling, M., Klingmüller, U., and Timmer, J. (2009). Structural and practical identifiability analysis of partially observed dynamical models by exploiting the profile likelihood. Bioinformatics, 25(15):1923–1929.

Raue, A., Schilling, M., Bachmann, J., Matteson, A., Schelker, M., Kaschek, D., Hug, S., Kreutz, C., Harms, B. D., Theis, F. J., Klingmüller, U., and Timmer, J. (2013). Lessons learned from quantitative dynamical modeling in systems biology. PLOS ONE, 8(9):e74335.

Raue, A., Steiert, B., Schelker, M., Kreutz, C., Maiwald, T., Hass, H., Vanlier, J., Tönsing, C., Adlung, L., Engesser, R., Mader, W., Heinemann, T., Hasenauer, J., Schilling, M., Höfer, T., Klipp, E., Theis, F., Klingmüller, U., Schöberl, B., and Timmer, J. (2015). Data2Dynamics: a modeling environment tailored to parameter estimation in dynamical systems. Bioinformatics, 31(21):3558–3560.

Ruscone, M., Vazquez, M., and Valencia, A. (2026). Intelligent tool orchestration for rapid mechanistic model prototyping: MCP servers as AI-biology interfaces. npj Systems Biology and Applications, 12(1):145.

Saini, A. and Farnoud, A. (2025). QSP-Copilot: An AI-augmented platform for accelerating quantitative systems pharmacology model development. CPT: Pharmacometrics & Systems Pharmacology, 14(11):1775–1786.

Savoglidis, G., da Silveira dos Santos, A. X., Riezman, I., Angelino, P., Riezman, H., and Hatzimanikatis, V. (2016). A method for analysis and design of metabolism using metabolomics data and kinetic models: Application on lipidomics using a novel kinetic model of sphingolipid metabolism. Metabolic Engineering, 37:46–62.

Schälte, Y., Fröhlich, F., Jost, P. J., Vanhoefer, J., Pathirana, D., Stapor, P., Lakrisenko, P., Wang, D., Raimúndez, E., Merkt, S., Schmiester, L., Städter, P., Grein, S., Dudkin, E., Doresic, D., Weindl, D., and Hasenauer, J. (2023). pyPESTO: a modular and scalable tool for parameter estimation for dynamic models. Bioinformatics, 39(11):btad711.

Schmidt, M. D., Vallabhajosyula, R. R., Jenkins, J. W., Hood, J. E., Soni, A. S., Wikswo, J. P., and Lipson, H. (2011). Automated refinement and inference of analytical models for metabolic networks. Physical Biology, 8(5):055011.

Schmiester, L., Schälte, Y., Bergmann, F. T., et al. (2021). PEtab—Interoperable specification of parameter estimation problems in systems biology. PLOS Computational Biology, 17(1):e1008646.

Shojaee, P., Meidani, K., Gupta, S., Farimani, A. B., and Reddy, C. K. (2025). LLM-SR: Scientific equation discovery via programming with large language models. In International Conference on Learning Representations.

Smith, L. P., Bergmann, F. T., Chandran, D., and Sauro, H. M. (2009). Antimony: a modular model definition language. Bioinformatics, 25(18):2452–2454.

Thiele, C., Wunderling, K., and Leyendecker, P. (2019). Multiplexed and single cell tracing of lipid metabolism. Nature Methods, 16(11):1123–1130.

Tsutsui, M., Arakane, K., and Okada, M. (2026). Literature-derived, context-aware gene regulatory networks improve biological predictions and mathematical modeling. Bioinformatics, 42(8):btag101.

Villaverde, A. F., Pathirana, D., Fröhlich, F., Hasenauer, J., and Banga, J. R. (2022). A protocol for dynamic model calibration. Briefings in Bioinformatics, 23(1):bbab387.

Wronowska, W., Charzyńska, A., Niena-ltowski, K., and Gambin, A. (2015). Computational modeling of sphingolipid metabolism. BMC Systems Biology, 9(1):47.

Xue, J., Schmidt, S. V., Sander, J., Draffehn, A., Krebs, W., Quester, I., De Nardo, D., Gohel, T. D., Emde, M., Schmidleithner, L., Ganesan, H., Nino-Castro, A., Mallmann, M. R., Labzin, L., Theis, H., Kraut, M., Beyer, M., Latz, E., Freeman, T. C., Ulas, T., and Schultze, J. L. (2014). Transcriptome-based network analysis reveals a spectrum model of human macrophage activation. Immunity, 40(2):274–288.

Ziaei Bideh, A. and Gryak, J. (2026). LLM-ODE: Data-driven discovery of dynamical systems with large language models. In Proceedings of the Genetic and Evolutionary Computation Conference, pages 853–860, New York, NY, USA. ACM.

