## Supplementary Information for "Large language models complement established workflows for mechanistic model construction and refinement in systems biology"

**S5 Results 4: Hypothesis testing on unpublished sphingolipid data 53**

### S1 Details of the workflow

The terms used in this and the following notes are collected in a glossary at the end of this note (S1.10).

#### S1.1 What each agent receives and returns

Every `agent` is one call of Claude Code with a defined context and no memory of earlier calls. The `module engine` assembles the context from the run record, so an `agent` receives exactly the inputs listed below and nothing else, and it returns one structured reply shaped by its output contract. Only the `knowledge agent` and the `research agent` may reach the web. No `agent` that proposes structure sees a measurement value, and the held-out conditions reach no `agent`.

| **agent** (module) | **receives** | **returns** |
|---|---|---|
| `knowledge agent` (M0, before the run) | the intent, the project description, the required reactions, the user's literature, the search horizon and the excluded sources; web access if online | species with identifiers, reactions marked essential or not, facts tagged per `agent`, every source classified by kind, context, and direction |
| `knowledge verification agent` (M0, online only) | one quoted passage with the cached text around it and the claim it is cited for; nothing else | a second classification of that source; never blocks |
| `research agent` (M1, revision) | one question from the `construction agent` or the `revision agent`, the search horizon and the excluded sources; web access | sources for that question, graded by the `module engine` and merged into the knowledge file |
| `construction agent` (M1) | the intent, the project description, the domain packs (short texts on the kind of system and the measurement technique), the user's literature as claims with their quotes, the essential reactions with grades and sources, the facts meant for construction, in advisory mode every other reaction as prior knowledge, and the shape of the data in words (curve shapes and timescales per observable, level against the control, monotonicity in dose); no measurement value, no held-out condition | the base model as Antimony text with a parameter table (nominal value, bounds, and source per parameter, each tested feature as a switch whose base value turns it off), a build report with one entry per modelling decision (rationale, alternatives, evidence, confidence), numbered questions for the user, an entity glossary, and optionally expansions it could not settle, or a research request instead of a guess |
| `expansion agent` (M2) | what the `construction agent` received, the tested base model with the decisions of its build report, the `construction agent`'s suggested expansions, and every reaction of the knowledge file with its grade at the expansion floor (the base model's floor lowered by one band) | two to five expansions, each a declared difference from the base model that tests one mechanism (a switch turned on, a value changed, a reaction replaced or added), with every new parameter's nominal value, bounds, and source |

| **agent** (module) | **receives** | **returns** |
|---|---|---|
| `construction verification agent` (M2) | the intent, the project description, the base model and every candidate, the build report, and the facts that parameters cite, in a fresh context; not the data, not the user's literature | findings per aspect: whether the models do what the intent asks, whether the build report's decisions are justified, whether the cited facts were applied correctly; never blocks |
| `data extraction agent` (M3, once per run) | the intent, the project description, a preview of each data file (header and first rows), and a summary of the experimental design (design columns with their levels, replicates per condition, absent combinations); never the model, never a measurement value | the meaning of each column (time, value, condition, replicate, error column and its type, units), optional processing steps from a closed vocabulary (normalise by a group's sum, subtract a reference, scale units), and flagged rows; exclusion is the user's decision |
| `observation agent` (M4, once per round) | the intent, the project description, the domain packs, the model's entity glossary and a summary of its species, parameters, and kinetics with the magnitudes every species reaches at the nominal parameters, the observable labels with units and error type, the noise statistics computed from the replicates (how the spread grows with the signal, zeros and negatives, the range), and the knowledge file's facts about the assay; no measurement value | for every observable the formula on the model's species, the transformation (linear or logarithmic), the noise distribution and its shape (constant, proportional, additive and proportional, or scaled with a gain), the scaling and offset parameters (per observable, condition, or time point, shared or not), and one rationale; an alternative it cannot settle is recorded as an expansion for a later round |
| `condition agent` (M4, once per round) | the same context without the noise statistics, the condition labels, the experimental design summary, the observation draft, the basal state the `construction agent` declared, and the facts about how a perturbation enters the system | for every condition what it sets (a dose, an input, a parameter that varies by condition) and whether the model is first run to steady state with the stimulus removed |
| `assembly verification agent` (M4) | the observables, conditions, and parameters tables and the project description, in a fresh context | findings on four aspects: the noise model against the readout technique, the conditions against the protocol, scaling and offset against the units, the observable mapping against what the assay measures; never blocks |
| `interpretation agent` (M7) | the intent, the project description, the computed evaluation, the residual diagnostics, and the three computed lists (what the data could not distinguish, what they nonetheless determined, what is finite but poorly determined); never the user's literature; it may request further diagnostics twice (the residuals of a series, a profile path, the model at given parameter values, the residuals grouped by a data column) | a diagnosis, hypotheses for the misfit, its own three lists, an answer to each success criterion, and a recommendation to accept, revise, or stop; it decides nothing |

| **agent** (module) | **receives** | **returns** |
|---|---|---|
| `revision agent` (between rounds) | the selected model and the selection table, every model fitted in the run, the expansions declared but never fitted, the misfit evidence and residual diagnostics, the evaluation report, the pressure of the fit on the parameter bounds, the drift of quantities the project description calls constant, the `interpretation agent`'s hypotheses, the best model's assembly, the intent, the project description, the domain packs, the user's literature, and the knowledge file's reactions with their grades at the next round's floor; it may request diagnostics twice and one targeted search | revisions, each one mechanism with one change (of the reactions, of the observation model or the condition encoding, or a split of one parameter into one copy per group) with its added parameters, the question it tests, and the data that would separate it from the base; it accounts for every untested expansion and names what the data cannot settle and which experiment would |
| `orchestrating agent` (an interactive session, unlike the other `agents`, which are single calls) | the interaction requests of the `module engine`, each with the question, the evidence, the consequences of every answer, and the answer forms, and the conversation with the user | the user's answer as given, or an answer on the user's standing instructions, marked as such in the record |

#### S1.2 Checks per module

The `module engine` checks every `agent` reply and every artefact without an LLM. A blocking check that fails sends the work back to the `agent` that wrote it, with the check in words and the history of its earlier attempts, three attempts in all, after which the user is asked. A malformed reply (one that violates its output contract) is sent back with the error quoted and counts as an attempt. Advisory checks report and never block. A redraft the user asks for in their own words does not count against the three attempts.

| **module** | **blocking checks** | **advisory** |
|---|---|---|
| M1, base model | the Antimony text builds to valid SBML; every symbol resolves; every estimated parameter is read by the model; the model integrates at the nominal values and across the parameter bounds; no state becomes negative; the required entities are present; the declared constants stay constant; the entity glossary covers every species; every required reaction is in the model; every piece of the model holds a required entity; every fact a parameter cites is a fact the knowledge file has | a conservation-law scan; a screen for missing dynamics; the origin of each reaction (user-provided, database-supported, literature-supported, or inferred) |
| M2, candidates | every candidate is built by the `module engine` from the base model and its declared difference, so nothing else can differ; a change to a parameter the base model does not have, an undeclared symbol, and two expansions identical once built are refused by name; the M1 checks run on every candidate; a failing candidate sends the whole set back | in advisory mode an addition below the expansion floor is kept and flagged; in strict mode it is dropped after a research request has failed to raise its grade |

| module | blocking checks | advisory |
|---|---|---|
| M3, data | the rows are well formed; units and the error type are declared for every observable; every processing step is from the closed vocabulary; the held-out conditions named in the settings resolve on the data's own columns | rows the `agent` flags as suspicious; their exclusion is the user's decision |
| M4, PEtab problem | every formula compiles; every symbol resolves; every data observable and condition is explained and none is invented; the noise model is declared for every observable; every $\sigma$ reaches the model only through its own observable; the parameters are bounded and scaled with nominal values inside the bounds; PEtab's own linter passes; the pre-equilibrations reach a steady state; every piece of the model is seen by a readout | the simulated and measured magnitudes are compared; the degrees of freedom are counted |
| M5, solver | the trajectory at two tolerances agrees (otherwise tighter tolerances are proposed); every pre-equilibration settles | the cost of the gradient (otherwise adjoint sensitivities are proposed, S1.5); the parameter combinations no output depends on near the nominal values |
| M6, fit | at least three starts within 0.5 of the best objective (the plateau of the waterfall); a projected gradient norm at the optimum of at most 10; at most 30 % of the starts lost to a failed integration; a failure opens the convergence interaction point rather than a redraft | a parameter at a bound is flagged and carried into the report |
| M7, evaluation | the criterion arithmetic agrees with PEtab Select, or the run stops; the `interpretation agent`'s reply lists every computed undetermined parameter combination, credits every determined one, and names only ids the problem has, with one retry | the misfit indicators and the parameters near a bound, reported with their distance from the threshold |

#### S1.3 Interaction points

At every interaction point the `module engine` writes an interaction request with the question, the evidence, the consequences of each answer, and the answer forms; the `orchestrating agent` presents it and returns the user's answer, which is recorded with who gave it. The user may give standing instructions in advance, for example before an overnight round, and the `orchestrating agent` then answers on them and says so in the record. Unattended, a run stops after three rounds.

| interaction point | when | the user sees | answers |
|---|---|---|---|
| evidence floor | after M0, before the run | every essential reaction with its grade, source, origin, and flags; the suggested floor and its rule; which essentials leave the base model at each higher band; the facts per `agent` | the floor, the step per round, which unsupported essentials to keep or drop, the mode |

| **interaction point** | **when** | **the user sees** | **answers** |
|---|---|---|---|
| intent freeze | start of the run | the intent, the essential reactions, the shape of the data in words, the difference from the intent the knowledge module saw | freeze, or edit and stage again |
| base model | after M1, when enabled in the settings | the base model with its build report and the `agent`'s questions | accept, redraft in the user's words, or stop |
| model candidates | after M2 | each candidate as its difference from the base model, where each reaction comes from, the floor flags, the verification findings, the `agent`'s questions | which candidates to fit; redraft the expansions or the base model; stop |
| compile | after M4 | the PEtab problems and the assembly verification findings | approve, redraft, or stop; everything after this costs compute |
| budget | after the M5 probe, at the first candidate that costs anything | the time per start, the objective values and failed starts of the probe, the projected cost for the proposed starts and workers, the solver proposals | the number of starts, the solver settings (tighter tolerances, adjoint sensitivities), the optimiser; later candidates reuse the answer when their probe looks like the first one's |
| convergence | after M6, only when a fit check fails | the failed check with its numbers, the waterfall | accept with a reason that is recorded as a caveat; refit with more starts or tighter settings; compare optimisers on the same starts (once per run); stop |
| verdict | after M7 | each candidate with its criterion, difference, weight, and held-out result; the best model and why it wins; the indistinguishable set; the misfit indicators with their distance from the threshold; the `interpretation agent`'s recommendation, flagged where it disagrees with the `module engine`'s default | sign the verdict, ask for another round, ask for profiles, or stop |
| revision approval | after the `revision agent` | each revision with its parameters, the question it tests, and the data that would separate it; revisions held back by the floor | buy the revisions one at a time on the selected model, or every combination, or on another base of the user's choice; send the proposal back in the user's words; end the run |
| failure point | after three failed attempts of an `agent` | the attempts with the check each failed | grant more attempts, change the floor (strict mode), or stop |

#### S1.4 Evidence grading

The `knowledge agent` classifies each source by kind and context and never grades. The `module engine` assigns the grade from Table S1: the bands are tried from the strongest down and the first rule an entry satisfies decides. One source cited twice counts once, a source the `knowledge verification agent` classifies lower lowers the grade, and a source that contradicts the reaction marks it contested and caps its grade at medium. The suggested floor is the lowest grade among the supported essential reactions that the user did not require, and the user keeps or changes it.

Table S1: **The evidence grading rules.**

| Band or setting | Rule |
|---|---|
| high | one direct experimental source in the same system<br>one curated database entry in the same or a related system<br>two independent sources, each earning at least medium on its own |
| medium | one direct experimental source in a related system<br>one mechanistic model in the same or a related system<br>two independent reviews in the same or a related system |
| low | one review in the same or a related system<br>one direct experimental source in a different system<br>one mechanistic model in a different system<br>two independent indirect sources (any system) |
| weak | any counting source |
| counting source | an evidence item whose identifier is not unresolvable, that is inside the search fence and whose class is in the vocabulary; a source cited twice counts once |
| none | no counting source (the bands are tried strongest first; the first rule satisfied decides) |
| not applicable | offline: nothing was searched, so no grade applies |

#### S1.5 Simulation and estimation settings

**Sensitivity method and tolerances.** At the probe the `module engine` times one evaluation of the residuals and one of the residuals with their sensitivities. When the sensitivities cost more than eight times the residuals (a setting), it proposes adjoint sensitivities and states the consequence: adjoint computes the gradient without per-residual sensitivities, so the Fisher information falls back to finite differences and Fides runs without its Fisher-information Hessian. When the trajectories at two tolerances disagree, it proposes tighter tolerances. The user decides both at the budget interaction point, and with adjoint the five-start probe is run again before the budget is approved.

**Refits and the comparison of optimisers.** A refit after a failed convergence check uses one of three presets: three times the starts, a tighter solver (relative tolerance $10^{-10}$, more integration steps), or more optimiser iterations; the `module engine` recommends the order, with the tighter solver first when integrations failed. The comparison of optimisers runs the same starts with Fides,

Ipopt, and the gradient-based NLopt optimisers, writes the table into the report, and when one optimiser is conclusively better than the one that produced the fit, the candidate is refitted with it.

**Parameters, bounds, and priors.** Every estimated parameter is declared by the `agent` that introduced it with a nominal value, bounds, and a source. It is estimated on a $\log_{10}$ scale when its lower bound is positive and on a linear scale otherwise; noise parameters are always $\log_{10}$. Priors on parameters are not supported yet; they are one of the planned additions.

#### S1.6 Evaluation of the estimation results

**Model comparison.** The `module engine` computes the objective, the number of estimated parameters, and the AIC and BIC of every fitted candidate, brings candidates with different transformations of the observables onto one measurement scale first, and selects by the declared criterion. Beside the criterion it reports the differences to the best candidate, the Akaike weights, the indistinguishable set, and whether the selection is decisive (a weight of at least 0.7). Its arithmetic is cross-checked against PEtab Select, which must agree or the run stops.

**Held-out prediction.** If held-out conditions are declared, every candidate is simulated under them. Where the held-out data have replicates, the prediction error is compared with the replicates' own noise; otherwise a normalised RMSE of at most 0.3 is required. A candidate that fails cannot be the best model whatever its criterion.

**Misfit evidence.** For the best model the `module engine` tests the residuals against the assay's own repeatability, the noise standard deviation estimated from the replicates of each observable, without reference to any other model. Two indicators can open the misfit flag on their own: a $\chi^2$ test, $\chi^2 = \sum_i n_i\,(\mathrm{RMSE}_i/\sigma_i^{\mathrm{rep}})^2$ over the observables $i$ with $n_i$ training points each, on $n - n_\theta$ degrees of freedom, with $p < 0.01$ as misfit; and the ratio of the fitted $\sigma_i$ to the replicate noise $\sigma_i^{\mathrm{rep}}$, with 1.3 as the limit, which says that the error model is absorbing structure. While the flag is open, the `revision agent` may not recommend stopping, and after a selection round the `expansion agent` is asked what the model space was missing. Beside the flag, each residual series is reported with its RMSE in units of the replicate noise, its largest deviation, and its sign pattern over time, the distribution of the residuals is tested against the assumed noise model ($p < 0.01$) with the direction in which the quantile plot bends, and every parameter within 2 % of a bound is listed.

**Autocorrelation of the residuals.** For every residual series (one observable in one condition) the `module engine` computes the lag-1 autocorrelation, combines the series into one $z$-statistic weighted by the square root of the number of points per series, and tests it one-sided for positive autocorrelation at $p < 0.05$. The result is reported with the misfit evidence and corroborates it, but never opens the misfit flag alone.

**Residual forms.** Each residual series of the best model and of the base model is fitted with four forms, constant, linear, exponential, and rise and fall (the difference of two exponentials, $a\,(e^{-t/\tau_2} -$

$e^{-t/\tau_1}) + c$), and the form is chosen by small-sample AICc against the constant with a margin of 2. The amplitude is reported in units of the replicate noise and the timescales in the data's own time unit, and series whose forms agree are grouped into one sentence of the report, so that a systematic deviation shared by several readouts is read as one finding.

**Identifiability.** The Fisher information matrix at the optimum is $J^\top J$ from the residual sensitivities on the estimation scale, or finite differences when adjoint sensitivities were chosen. Its eigendirections eight or more decades below the largest eigenvalue are reported as flat, each named by the parameters that compose it. From this the `module engine` writes three lists: what the data could not distinguish (the flat combinations), what they nonetheless determined, and what is finite but poorly determined. The `interpretation agent` receives the three lists and must account for every entry. Profile likelihoods are computed on request at the verdict, and always when the inference target is parameter values.

#### S1.7 The run record

The run record is one HDF5 file per run, and it is the only state of a run: every stage reads its inputs from the record and writes its outputs to it before the next stage starts, so a run that is paused at an interaction point, or stopped, resumes from the record alone. Its groups are:

**run** the run's metadata: engine version, git state, package versions, the hashes of the settings and of the staged case, the declared policy, and the start of every session

**input** the snapshot of the staged folder taken at the intent freeze (project description, intent, settings, data files, supplied files, the knowledge file with its cached sources), with where it came from

**intent** the frozen intent

**rounds/round_k** per round: the base model (Antimony, SBML, parameter table, entity glossary, build report, the origin of every reaction), the declared expansions and the candidate set, the extracted data with the extraction report, the shared assembly, and per candidate its PEtab problem, its fit with every start, and its diagnostics; the evaluation with the selection table, the misfit evidence, and the three lists; every `agent` call with its prompt, reply, LLM version, and cost, and every attempt that was sent back; every check with its result; every interaction request with the answer and who gave it (the user, a standing instruction, or a declared policy); the decisions of the round as a readable document

**questions** the questions the `agents` raised for the user and the answers

**ledger** every event of the run in order: stage starts and ends, check results, `agent` calls, costs, interaction requests and answers

**timing** where the wall time went: sessions, waiting at interaction points, `agent` calls, compute, and the engine's own time

**fit index** the fits in the record by their key, for reuse

**outcome** how the run ended

Every entry carries a tier (core, audit, or bulk), and a copy for dissemination keeps the tiers asked for. The record is validated against a schema when it is written. A command-line reader prints any entry, HTML reports are rendered from it, and the models in SBML and PEtab can be used without the workflow. Table S3 gives the effort of each application run from its record.

#### S1.8 Settings and thresholds

Table S2 lists the settings of the workflow that a user changes, in the settings file of a run or at the interaction point named in the last column.

Table S2: **Settings of the workflow.** Every value is a default that a run may change: in the settings file of the run, or by the user at the interaction point named in the last column. The application runs used these defaults unless Table S3 says otherwise.

| setting | default | changed |
|---|---|---|
| LLM and reasoning effort of the `agent` calls | Claude Code's default model; the effort per run, optionally per `agent` (Table S3) | settings file |
| model candidates per round | 2 to 5 from the `expansion agent`, at most 8 | settings file |
| evidence floor and mode | suggested by the `module engine`, advisory mode | by the user after M0 |
| evidence floor of the expansions | the base model's floor lowered by one band per round | by the user after M0 (the step) |
| held-out conditions | none | by the user at staging |
| starts per candidate | 100, within 10 to 1000 | by the user at the budget interaction point |
| optimiser and solver settings | Fides, forward sensitivities | by the user at the budget interaction point, with the option to compare optimisers once per run |
| selection criterion | AIC, cross-checked with PEtab Select; candidates within 2 units are indistinguishable; decisive at an Akaike weight of 0.7 | settings file |
| held-out prediction | against the replicate noise with $p \geq 0.01$, or normalised RMSE at most 0.3 without replicates | settings file |
| misfit evidence | $\chi^2$ against the replicate noise with $p < 0.01$, fitted $\sigma$ over replicate noise above 1.3 | settings file |
| rounds | unlimited while the user answers the verdict, 3 unattended | by the user at the verdict, settings file |

#### S1.9 Settings of the application runs

Table S3 gives, per application, the settings of the run and its effort.

#### S1.10 Glossary

##### General

**Run** one spin-up of the workflow, which may involve many Rounds of model refinement.

**Round** one sequential execution of Modules M1 - M7 and a possible Revision. The run always starts from round 0. Controlled by the Central Engine.

**Orchestrating Agent (OA)** when using Octomod, the user first interacts with the OA to complete the Staging. Afterwards, the OA initiates a Run and a Round by starting the Central Engine. The OA combines outputs from other agents and Checks, and relays each human interaction with a summary and a recommendation.

**Module** a specific function performed in a Round, for example, parameter estimation. A Module typically consists of some task-specific programs, as well as Checks, sub-agents, and user interaction.

**Agent Output Contract** the required structure of an agent's output. The run fails if the agent fails to produce correctly structured output after 2 (default) Redraft attempts.

**Run Record** a single HDF5 file in the `runs/` folder. It holds the inputs, every artifact, every agent call, every Check and every decision of that specific run, which can contain many rounds.

**Central Engine** the deterministic Python program that controls and calls the Module Engines, calls the Revision Agent, and updates the Run Record.

**Module Engine** deterministic Python programs that call the sub-agents in each Module (except the OA and the Revision Agent), run the Checks and write the Run Record.

**Check** a deterministic program that ensures Module-specific requirements are satisfied and can stall or advise, in which case additional information is passed to the Module Agent and the user.

**Redraft** a request from the Module Engine to an agent requesting changes in the agent's output due to either a failure of satisfying the Agent Output Contract or a failure of a Check. In addition, this can be a request from the User with specific redrafting directions.

**Interaction Request** the summary the Module Engine produces for the upcoming human interaction.

**Output Attempt History** upon a Redraft, the information that the agent receives: earlier (re)draft(s) and why each failed, so it revises instead of starting over.

**Model Candidate** the Base Model plus some changes, which can either be expansions or deletions.

Table S3: **The application runs.** Settings and effort per run, from the run records. Wall time is the time from the start of the run to its end, estimation time the computation of the fits, and `agent` time the time spent in `agent` calls. The rest of the wall time is the `module engines`' own work (checks, model building, rendering) and waiting at the interaction points.

| | CTD (Results 1) | Macrophage (Results 2) | PBMC (Results 3) | Sphingolipid (Results 4) |
|---|---|---|---|---|
| run id | `ctd_blind_local_v3_opus55_20261004_022720` | `macrophage_xue_20260929_210535` (run 1), `macrophage_xue_v2_20260930_131832` (run 2) | `bodenmiller2012_crosstalk_r5_20260930_232653` | six runs, `sphingolipid_metabolism_20260929_120748` (run 1) to `sphingolipid_metabolism_v6_20261001_092043` (run 6) |
| LLM of the `agent` calls | `claude-opus-5-5` | `claude-opus-5-5` | `claude-opus-5` | `claude-opus-5-5`; the `research agent` `claude-opus-5` in runs 1, 2, 4, and 6 |
| effort | extra-high | high | extra-high (knowledge module: maximum) | maximum (knowledge module: maximum) |
| knowledge module | off | offline, advisory mode | online, advisory mode, floor low | online, advisory mode, floor low; run once, before run 1 |
| starts per model | 100 | 300 | 100 | 500 |
| optimiser | Fides, forward sensitivities | Fides, forward sensitivities | Fides, forward sensitivities | Fides, forward sensitivities |
| rounds | 2 | 5 (run 1) and 1 (run 2) | 5 | 1 per run; run 2 stopped before any fit |
| `agent` calls | 19 | 29 (run 1) and 14 (run 2), plus 1 of the knowledge module | 30, plus 3 of the knowledge module | 48 (9, 1, 8, 9, 10, and 11 per run), plus 2 of the knowledge module |
| cost (USD) | 25.40 | 17.48 (run 1) and 8.99 (run 2), plus 0.56 for the knowledge module | 119.01, plus 20.06 for the knowledge module | 111.49 (27.72, 3.45, 16.05, 16.95, 23.64, and 23.68 per run), plus 5.78 for the knowledge module |
| wall time | 21 h 55 min | 15 h 48 min (run 1) and 1 h 57 min (run 2) | 139 h 31 min | 42 h 17 min (11 h 32, 4 h 3, 4 h 3, 16 h 2, and 6 h 37 min) |
| estimation time | 2 h 0 min | 2 h 14 min (run 1) and 1 h 24 min (run 2) | 102 h 30 min | 1 h 48 min (54, 12, 21, 13, and 8 min) |
| `agent` time | 2 h 49 min | 1 h 6 min (run 1) and 21 min (run 2) | 7 h 6 min | 15 h 1 min (4 h 24, 1 h 48, 2 h 11, 4 h 5, and 2 h 33 min) |

#### Staging related

**Staging** an interactive process where the user and the OA work together to generate information in a suitable format for a Run. This includes the research question, dataset(s), other user-provided information, and data pre-processing.

**Project Description (description.md)** all information provided by the user, e.g., the experiment setup and additional literature, which can also be edited by the OA.

**Intent (intent.yaml)** the project in a structured form: scope, Required Reactions, Required Species, desired predictions, perturbations, Success Criteria, Inference Target.

**Success Criteria** the criteria the run must satisfy. If not satisfied, another round must be started to refine the models.

**Required Species** a species the user says the model must contain.

**Required Reaction** a reaction the user says the model must contain.

**Inference Target** what the run is for: `prediction`, `parameter_values` or `mechanism_discrimination`.

**Settings (settings.yaml)** workflow hyperparameters for the run, e.g., the Evidence Floor, Strict / Advisory Mode.

**Holdout Data** data that is not used for training, but used for final evaluation of a model.

#### M0 - Knowledge module

**Knowledge Module (M0)** this Module is done before the Run and checks what is known (either Online or Offline) about the system that is being modelled.

**Knowledge Agent** surveys the system for issues, classifies every information source according to reliability, and coordinates other tasks within the Knowledge Module.

**Online / Offline** whether the Knowledge Agent searches literature and databases (online) or answers from its own pre-trained knowledge (offline).

**Knowledge Verification Agent** for Online sources: re-reads each cited document and classifies the source again. If there is a disagreement, the lower grade is used.

**Essential Reaction** a reaction, determined by the Knowledge Agent, that is needed to answer the question.

**Evidence Grade** a grade (high, medium, low, weak, none, not applicable) the Knowledge Module's Module Engine gives each reaction based on a rule table.

**Evidence Floor** defined by the lowest Evidence Grade of all the Essential Reactions.

**Knowledge File (knowledge.json)** the Module's output: species, reactions, facts, grades, the Evidence Floor, and others. Written by the Knowledge Module's Module Engine.

#### Intent freeze

**Intent Freeze (human interaction)** the run's entry point. The user freezes the Intent before a Run starts; the Run will be evaluated according to how well the Intent has been achieved, and it reads only the snapshot of the staged folder taken at that moment.

#### M1 - Model building

**Advisory / Strict Mode** In Strict Mode, new reactions can only be added when their Evidence Grade is at or above the Evidence Floor. In Advisory Mode (default), new reactions can always be added, but a reaction whose Evidence Grade is below the Evidence Floor is flagged to the user for review.

**Construction Agent** writes the Base Model in an Antimony file.

**Research Request** if there is a question the Construction Agent cannot answer from the Knowledge File, a Research Request is sent to the Research Agent.

**Research Agent** conducts further literature searches and gives the results to the Construction Agent.

**Build Report** one entry per modelling decision with its rationale, alternatives, evidence, and confidence, plus the Construction Agent's questions for the user.

**Base Model** the model M1 successfully builds.

#### M2 - Model Expansion

**Model Expansion** extends the Base Model with new reactions or kinetics, to test different mechanisms. This results in 2-5 models that will be evaluated in the round.

**Expansion Agent** performs Model Expansion.

**Expansion Evidence Floor** the Evidence Floor will be lowered by one level every time an attempt to build a Model Expansion is made. This Expansion Evidence Floor is used for the specific Model Expansion. In strict mode, the reactions a Model Expansion adds must have an Evidence Grade at or above the Expansion Evidence Floor.

**Construction Verification Agent** an independent agent that reads the Model Report and checks it against the Intent and the Project Description, and whether the decisions are justified.

#### M3 - Data Extraction

**Experimental Design Summary** a summary of each experimental condition and the replicates per condition.

**Data Extraction Agent** interprets what each data column means for the Module Engine, which then moves the data into the corresponding files. For instance, the time points are moved to the time column of the `measurements.tsv` file.

#### M4 - PEtab assembly

**PEtab Problem** the standard files that a fit needs: SBML model `model.xml`, `observables.tsv`, `conditions.tsv`, `parameters.tsv`, `measurements.tsv`, and a connecting YAML file `model.yaml`. One per Model Candidate.

**Observation Agent** writes the `observables.tsv` file containing the observation model of each measured quantity (formula on model species, transformation, noise model, scaling and offset).

**Condition Encoding** how each experimental condition is structured in the model, and whether the model first runs to steady state (pre-equilibration).

**Condition Agent** writes the Condition Encoding.

**Assembly** the Module Engine combines the observables and the Condition Encoding. Written once per round, shared by all its Model Candidates.

**Assembly Verification Agent** audits the PEtab tables against the experiment (noise against technique, conditions against the description, units, what the assay measures).

**Compile** the user approves the PEtab Problems before any compute is spent.

#### M5 - Compile and probe

**Solver Checks** the Module Engine checks the AMICI simulation for tolerance agreement, sensitivity, steady states, and Structural Nonidentifiabilities.

**Structural Nonidentifiability** a parameter combination no output depends on. Found by a local check near the nominal parameter values: a zero singular value of the sensitivity matrix, i.e. a zero eigenvalue of the Fisher Information Matrix (FIM).

**Probe** five optimisation starts. Measures the time per start and what fails, before the full estimation.

**Budget** the user approves the number of starts and the solver settings, from the Probe results.

#### M6 - Estimation

**Multistart Optimisation** optimisation from different randomly initialized starts (100 by default, fixed seed, Fides) to avoid local minima.

**Waterfall Plot** a plot showing the sorted final objective values of the starts. A plateau at the bottom means the best fit was found repeatedly and the optimisation has converged.

**Ensemble** the starts within the likelihood-ratio threshold (1.92) of the best objective. The threshold is that of a 95% confidence interval with one degree of freedom, based on Wilks' theorem.

**Convergence** when a Post-fit Check fails, the user can accept with a reason, refit, compare optimisers, or stop.

#### M7 - Evaluation

**Selection Criterion** AIC by default. Ranks models by fit and parsimony.

**Best Model** the best Model Candidate by the Selection Criterion that also passes the Holdout Data prediction.

**Indistinguishable Set** the Model Candidates within 2 criterion points of the Best Model, meaning the data cannot tell them apart.

**Misfit Evidence** whether the Best Model's simulation falls within the standard deviation of the data, i.e. the replicates' own noise. It is quantified by a $\chi^2$ goodness-of-fit test at that noise level (misfit when $p < 0.01$) and by the ratio of the fitted noise to the replicates' noise (misfit above 1.3). Misfit is the default reason for another round.

**Autocorrelated Data Check** the Module Engine's check on the residuals for autocorrelation: each residual series is fitted with a form (constant, linear, exponential, rise-and-fall).

**Interpretation Agent** reads the evaluation and the results of the Autocorrelated Data Check; writes the diagnosis and the answer to each Success Criterion in the Intent.

**Verdict** the user can decide to approve the selection, do another round, compute profiles, or stop.

#### Revision

**Revision** the step after a Verdict if the user approves: proposes model revisions, and the user can approve them. This can either be a reaction change, an observation or condition change, or a Parameter Split or Combination.

**Revision Agent** reads the fits, residuals and Misfit Evidence, says what the Model Candidates from the last round were missing, and proposes model revisions. Called by the Central Engine.

**Parameter Split or Combination** one parameter takes a different value in each group (e.g. per cell population). If the fitted values do not differ, the copies should be combined.

**Revision Approval** the user approves the revisions and therefore starts a new Round, picks another Base Model, asks for a Redraft. Otherwise, the Run ends.

### S2 Results 1: CTD phosphorylation without user guidance

#### S2.1 Inputs to the run

The workflow received three inputs: the data file (36 relative intensities, nine phosphorylation counts at 0, 1, 2, and 16 h), the project description, and the intent, both given below in full. The knowledge module was switched off, and no agent had web access. The description was written from the Methods of the published paper only. It names the hypotheses the experimenters had in mind and the earlier conclusion, drawn by eye, that the kinase acts distributively, and it contains no result of the published analysis.

##### S2.1.1 The project description, as given

**The system.** RNA polymerase II carries a long, flexible tail on its largest subunit called the C-terminal domain, or CTD. The CTD is not a folded domain: it is a tandem array of short repeats, each seven amino acids long, with the sequence Tyr–Ser–Pro–Thr–Ser–Pro–Ser. In human Pol II there are 52 of these heptarepeats. During transcription the CTD is progressively decorated with phosphate groups, and the resulting pattern of marks is what recruits and releases the RNA-processing and elongation factors that act on the polymerase. Which marks are present, and where, is therefore functionally important. The kinase we are studying here is P-TEFb (the CDK9–cyclin T1 complex), which acts during transcriptional elongation.

**The experiment.** This is an in vitro kinase assay, not a cellular experiment. There is no transcription, no DNA, no polymerase, only a purified CTD substrate, purified kinase, and ATP in buffer. Substrate: a recombinant CTD construct containing 8 heptarepeats, not the full 52, present at 100 µM. Kinase: P-TEFb, catalytic; it is not consumed and its concentration does not change over the experiment. ATP: 3 mM at the start; each phosphorylation consumes one ATP. Sampling: four time points, 0, 1 h, 2 h and 16 h (0, 60, 120 and 960 minutes). Replication: one measurement per species per time point; there are no replicate measurements in this dataset.

**What the readout is, and what it cannot see.** The readout is mass spectrometry on the intact CTD construct. MS separates the substrate molecules by mass, and since each phosphate adds a fixed mass, it resolves how many of the eight repeats carry a phosphate, a number between 0 and 8. It does not resolve which repeats those are. A molecule phosphorylated on repeats 1 and 2 and a molecule phosphorylated on repeats 1 and 7 are the same mass and appear in the same peak. Everything downstream has to live with that: the data are nine numbers per time point, one for each phosphorylation count, and the positional arrangement behind those numbers is not observed. A second point about the chemistry, established previously for this kinase and these conditions: within a single heptarepeat, phosphorylation happens on either Ser2 or Ser5, but not both at once. So for the purposes of counting, each repeat is either modified or unmodified, one bit per repeat, and the distinction between the two serine positions does not need to be tracked.

**The data file, and a caveat you must handle.** The data file has one row per time point and nine columns, `phos0_rel` through `phos8_rel`: the relative MS signal for CTD molecules carrying 0, 1, . . . , 8 phosphates. These are relative signal intensities, not concentrations, and they are not comparable between time points as they stand. Mass spectrometry reports intensities on an arbitrary scale that drifts between injections. You can see this directly in the file: the nine numbers sum to 1.00 at $t = 0$, to 4.07 at 1 h, to 3.79 at 2 h and to 0.85 at 16 h, even though the total amount of substrate in the tube is conserved throughout. Any model of this data needs a way to map predicted amounts onto measured intensities, and that mapping is not the same at every time point. The measurement error is believed to be additive and normally distributed, with a single standard deviation for the whole experiment, the same for every peak, that scales from one time point to the next with that spectrum's overall intensity, the same factor that maps amounts onto intensities at that time point.

**What is already believed, and what is not settled.** Earlier work on this kinase, using this kind of hyperphosphorylation assay in human and in *Drosophila*, concluded that P-TEFb works distributively rather than processively: that it modifies a repeat, lets go, and rebinds, rather than holding on and running along the substrate. That conclusion came from looking at the shape of MS distributions by eye, and was never checked against a quantitative model. Beyond that, the mechanism is genuinely open. We do not know from these data what governs the rate at which any individual repeat gets modified, and that is what we would like a model to tell us. One specific possibility was put to us and we want it tested rather than assumed either way: that the modification of one repeat changes the rate at which its immediate neighbours in the array are modified, as opposed to the rate depending only on how many repeats of the molecule are already modified (cooperativity between repeats, which was also raised as a possibility to look at), or on nothing at all.

**What can be computed here.** Any model that simulates is acceptable, whether it tracks only how many repeats are modified or which repeats are modified (with eight repeats, 256 configurations). Models of a few hundred states and a thousand reactions simulate in seconds on the machine this runs on, so size alone is not a reason to prefer one description over another; choose the description the question needs. If a possibility can only be represented by tracking which repeats are modified, we would rather see that model fitted than the question settled by argument from the counts alone. Such a model can be written out in full, every state and every reaction listed one by one, and we would rather it were written out than avoided. The simulator reserves the single letters x, k, p, y, w, h and t, so no species or parameter may be named with one of them.

**The question.** Build a model of this time course that explains the measured distributions, and tell us what it implies about how P-TEFb modifies the CTD. Be explicit about what these data can and cannot establish; with eight repeats, four time points and no positional information, we expect there to be things the data simply cannot decide, and we would rather know which those are than have them glossed over. We would also like the earlier conclusion, drawn by eye, that the kinase acts distributively rather than processively, to be tested against a model and not assumed.

##### S2.1.2 The intent, as given

**Question.** Build a model of this CTD phosphorylation time course that explains the measured distributions, and say what it implies about how P-TEFb modifies the substrate. Report explicitly what these data can and cannot establish about the mechanism. In particular, test whether the modification of a repeat changes the rate at which its immediate neighbours in the array are modified, against the alternatives that the rate depends only on how many repeats are already modified (cooperativity between repeats) or on nothing at all, and say which of these the data can and cannot separate. Also test the earlier conclusion, drawn by eye, that the kinase acts distributively rather than processively.

**System.** RNA polymerase II CTD (8 heptarepeats, in vitro), phosphorylated by P-TEFb (CDK9/cyclin T1); mass-spectrometry time course. **Required entities.** CTD, P-TEFb, ATP. **Required predictions.** The mechanism of phosphorylation that best explains the measured distributions, identified and defended; a statement of which mechanistic questions these data cannot answer. **Perturbations.** None, a single in vitro condition sampled over time. **Timescale.** Minutes, horizon 960 min. **Scope exclusions.** Transcription, DNA and the polymerase itself; the distinction between Ser2 and Ser5 within a repeat. **Inference target.** Mechanism discrimination.

**Success criteria.** (1) The selected mechanism is reported with its information criterion, its Akaike weight and the indistinguishable set, honestly, whatever they turn out to be. (2) The observation model is stated explicitly, including how predicted amounts are mapped onto relative MS intensities at each time point. (3) Any mechanistic question the data cannot resolve is named as unresolved rather than decided.

###### S2.1.3 Settings and standing answers

The run used the default settings: 100 optimisation starts per model candidate with Fides and forward sensitivities, no held-out condition, and no cap on the number of rounds. Our answers at the interaction points were fixed before the run started: the intent as staged, all five model candidates as built, the default estimation settings, one revision round when the `interpretation agent` recommended it, and no further round after that. No information was added at any interaction point.

##### S2.2 The model candidates

**Base model (M1).** The `construction agent` wrote the base model in terms of the phosphorylation count: species $\mathrm{CTD}_n$, $n = 0, \ldots, 8$, for the CTD molecules with $n$ of the 8 repeats phosphorylated, ATP, ADP, and the kinase. The substrate starts at 100 µM in $\mathrm{CTD}_0$ and ATP at 3 mM, and each phosphorylation consumes one ATP. The rate at which one unmodified repeat of a molecule in state $\mathrm{CTD}_n$ is modified is

$$v_n = k_{\mathrm{mod}}\,[\mathrm{PTEFb}]\,\frac{[\mathrm{ATP}]}{K_{\mathrm{ATP}} + [\mathrm{ATP}]}\,c_p^{\,n}, \tag{S1}$$

and the transition $\mathrm{CTD}_n \to \mathrm{CTD}_{n+1}$ proceeds at $(8 - n)\,v_n$. Here $k_{\mathrm{mod}}$ is the modification rate per repeat, $K_{\mathrm{ATP}}$ the ATP affinity, and $c_p$ a factor applied once per phosphate already on the molecule, so that $c_p > 1$ accelerates and $c_p < 1$ slows the kinase as the molecule fills up. $K_{\mathrm{ATP}} = 50$ µM was fixed, so with 3 mM ATP the kinase is saturated with ATP, whereas in the published models the rate is proportional to the ATP concentration. The base model also carries processive encounters, in which the kinase modifies $j$ repeats in one binding event with continuation probability $q$ (weight $q^{j-1}(1 - q)$), and with $q = 0$ every encounter modifies one repeat.

**Candidates (M2).** The `expansion agent` wrote the five candidates of round 0 as settings and extensions of the base model:

- **Uniform distributive**: $c_p = 1$, $q = 0$ (6 estimated parameters: $k_{\mathrm{mod}}$, four scaling parameters, and $\sigma$).
- **Global effect**: $c_p$ estimated (7 parameters). The published Global-effect model applies one factor once any repeat is phosphorylated.

- **Processive**: $q$ estimated (7 parameters).
- **Neighbouring effect**: the rate of a repeat is multiplied by $a^m$, with $m \in \{0, 1, 2\}$ the number of phosphorylated immediate neighbours (7 parameters). The published Neighbouring-effect model applies the factor once, whether one or both neighbours are phosphorylated. This candidate depends on the positions of the phosphates and is written on the 74 classes described below.
- **Neighbouring + global effect**: $a$ and $c_p$ estimated (8 parameters), the combination the published analysis did not consider.

**The 74 classes.** A molecule with 8 repeats has $2^8 = 256$ phosphorylation patterns. Under the neighbour rule the rate of an unmodified repeat depends only on how many phosphates the molecule carries and on whether its two neighbours are phosphorylated, so patterns with the same multiset of gaps (stretches of unmodified repeats, each at a chain end or between two phosphates) evolve identically and can be summed into one class without approximation. The `expansion agent` wrote the two candidates with a neighbouring effect on 74 class states instead of 256 pattern states (86 species and 293 reactions, against 12 species and 36 reactions for the count-based candidates), and the published analysis simulated all 256 patterns.

**Revision (round 1).** The `revision agent` proposed an inhibition of the kinase by its product: free kinase binds the phosphorylated repeats, $\mathrm{PTEFb} \to \mathrm{PTEFb_I}$ at rate $k_{\mathrm{on,I}}[\mathrm{PTEFb}] \sum_n n\,[\mathrm{CTD}_n]$, and returns with $k_{\mathrm{off,I}} = 1\,\mathrm{min}^{-1}$ (fixed). Added to the round-0 winner this gives the selected model with 9 estimated parameters, fitted at $k_{\mathrm{mod}} = 5.14{\times}10^{-3}\,\mathrm{min}^{-1}$ per repeat, $c_p = 1.63$ per phosphate, $a = 0.311$ per phosphorylated neighbour, and $k_{\mathrm{on,I}} = 9.09 \times 10^{-4}\,\mathrm{M}^{-1}\,\mathrm{min}^{-1}$.

#### S2.3 Estimation and model selection

Table S4 lists every model of both analyses with its number of estimated parameters, negative log-likelihood, AIC, and BIC on all 36 measurements. All run models were estimated with 100 starts. In round 0 the model combining the neighbouring and the global effect was selected (AIC −166.4, Akaike weight 0.997 among the five candidates), and in round 1 the product-inhibition model was selected over it (AIC −172.8).

#### S2.4 Comparison with the published analysis

The observation and noise models of the two analyses are identical, so their likelihoods and AICs are directly comparable. Three differences in the model forms explain the different AIC of the same hypothesis in the two analyses (panel E of the CTD figure in the main text): the ATP dependence (saturated in the run, first order in the published models), the neighbour factor (per phosphorylated neighbour in the run, once in the published model), and the global factor (per phosphate in the run, once in the published model). Table S5 crosses these forms, re-implemented and refitted on all 36 points. The ATP form is worth 4.4 AIC units on the published Neighbouring-effect rule and

Table S4: **All models of both analyses on the 36 CTD measurements.** Number of estimated parameters $n_\theta$, negative log-likelihood, AIC, and BIC. Published values from Callenbach et al. (2026) and its S1 Text, run values from the run record. $\mathrm{BIC} = 2\,\mathrm{NLLH} + n_\theta \ln 36$.

| Analysis | Model | $n_\theta$ | NLLH | AIC | BIC |
|---|---|---|---|---|---|
| Published analysis | Uniform distributive | 6 | −58.478 | −104.96 | −95.45 |
| | Processive | 7 | −65.466 | −116.93 | −105.85 |
| | Global-effect | 7 | −72.757 | −131.51 | −120.43 |
| | Neighbouring-effect | 7 | −89.997 | −165.99 | −154.91 |
| | Directional | 9 | −90.912 | −163.82 | −149.57 |
| Octomod | Uniform distributive | 6 | −61.358 | −110.72 | −101.21 |
| | Processive | 7 | −80.952 | −147.90 | −136.82 |
| | Global effect | 7 | −81.641 | −149.28 | −138.20 |
| | Neighbouring effect | 7 | −84.253 | −154.51 | −143.42 |
| | Neighbouring + global effect | 8 | −91.202 | −166.40 | −153.74 |
| | + product inhibition (round 1) | 9 | −95.401 | −172.80 | −158.55 |

less than one unit on the selected model. The per-neighbour rule combined with the count factor is 7.8 units better than the published once-rule with the count factor. On the 18 points at 1 and 2 h alone, where the two analyses differ in form, the selected model refitted on these points has an AIC of −43.4 against −41.7 for the published model. The published Processive and Directional models were not re-implemented, because their exact definitions differ from the run's candidates, and their values are taken from the paper.

Table S5: **The model forms crossed, all 36 points.** ATP dependence (first order or saturated), neighbour rule (none, the published rule "once, whether one or both neighbours are phosphorylated", or the run's rule "a factor per phosphorylated neighbour"), and count factor (none or a factor per phosphate), each refitted from 8 starts. Fitted factors: $a$ neighbour, $c_p$ count.

| ATP | neighbour rule | count factor | $n_\theta$ | NLLH | AIC | fitted factors |
|---|---|---|---|---|---|---|
| first order | none | no | 6 | −58.5 | −105.0 | |
| first order | once (published) | no | 7 | −90.0 | −166.0 | $a = 2.26$ (the published model) |
| first order | per neighbour | no | 7 | −86.0 | −158.0 | $a = 1.90$ |
| first order | none | yes | 7 | −82.9 | −151.8 | $c_p = 1.15$ |
| first order | once (published) | yes | 8 | −90.1 | −164.3 | $a = 2.45$, $c_p = 0.99$ |
| first order | per neighbour | yes | 8 | −94.0 | −172.0 | $a = 0.31$, $c_p = 1.61$ (4 of 8 starts) |
| saturated | none | no | 6 | −61.4 | −110.7 | (run: Uniform distributive) |
| saturated | once (published) | no | 7 | −87.8 | −161.6 | $a = 2.14$ |
| saturated | none | yes | 7 | −81.6 | −149.3 | $c_p = 1.14$ (run: Global effect) |
| saturated | once (published) | yes | 8 | −88.2 | −160.4 | $a = 2.50$, $c_p = 0.97$ |

#### S2.5 Where the phosphates sit

The predicted positions of the phosphates come from a re-implementation of both selected models on all 256 phosphorylation patterns, simulated at the fitted parameters. For molecules with $m$ phosphates at a given time we count the adjacent phosphate pairs per molecule and divide by the expectation under random placement of $m$ phosphates on 8 repeats, which for $m = 2$ is $7/28 = 0.25$ of the molecules. Table S6 gives the ratios at 1 and 2 h. Among molecules with two phosphates at 1 h, the published Neighbouring-effect model places them on adjacent repeats in 46.5 % of the molecules and the Octomod model in 7.7 % (panel D of the CTD figure in the main text), and the clustering vanishes as the molecule fills up.

Table S6: **Adjacent phosphate pairs per molecule relative to random placement**, by number of phosphates on the molecule, at 1 and 2 h. A ratio of 1 means random placement, above 1 clustered, below 1 spread out.

| model | time | 2 | 3 | 4 | 5 | 6 | 7 |
|---|---|---|---|---|---|---|---|
| published Neighbouring-effect | 1 h | 1.86 | 1.47 | 1.26 | 1.13 | 1.06 | 1.01 |
| | 2 h | 1.99 | 1.54 | 1.30 | 1.15 | 1.06 | 1.01 |
| Octomod selected | 1 h | 0.31 | 0.38 | 0.51 | 0.71 | 0.87 | 0.97 |
| | 2 h | 0.27 | 0.33 | 0.46 | 0.67 | 0.86 | 0.96 |

### S3 Results 2: Octomod facilitates knowledge-guided model refinement

This section documents the two Octomod runs on the macrophage transcript data of Xue et al. (2014) behind the main text: the data and the noise model (S3.1), the inputs to the runs (S3.2), the five rounds of the first run with every model candidate (S3.3), the equations and fitted values of the models (S3.4), and the second run, which recalibrated the final model, with its profile likelihoods (S3.5). Every number comes from the run records `macrophage_xue_20260929_210535` (run 1) and `macrophage_xue_v2_20260930_131832` (run 2). Models are numbered 0 to 5 as in the main text: model 0 is the base model and model $k$ the model taken forward from round $k$. Rounds are counted from 1, as in the main text and its figure; since a run starts from round 0 (S1.10), the record stores round $k$ under `rounds/round_`$(k-1)$.

#### S3.1 Data and noise model

**Source.** The data are part of the human macrophage compendium of Xue et al. (2014) (GEO SubSeries GSE46903 of SuperSeries GSE47189). CD14$^+$ monocytes from buffy coats of healthy donors were differentiated into macrophages for 72 h with GM-CSF and then stimulated with IFN-$\gamma$ (200 IU/ml) or kept in culture without a stimulus, and lysed at the stated time; the 0 h arrays are macrophages lysed at the end of differentiation. Expression was measured on Illumina HumanHT-12 V3 BeadChips (GEO platform GPL6947), one array per sample. We used the deposited values, which are $\log_2$-transformed and quantile-normalised across the 384 arrays of the study, without a second logarithm or any renormalisation, and the one probe per transcript the platform carries: IRF1 `ILMN_1708375`, SOCS1 `ILMN_1774733`, CXCL9 `ILMN_1745356` and CXCL10 `ILMN_1791759`. The study computed an array background of 6.747 $\log_2$ units. By the detection p-values of the deposit, IRF1 is detected on every eligible array, while SOCS1 is not detected on 11, CXCL9 on 47 and CXCL10 on 28 of the 85 eligible arrays, almost all of them untreated arrays at 6.1 to 6.6 $\log_2$ units, the array's floor.

**Selection.** 85 arrays of the deposit match the selection (platform GPL6947, differentiation with GM-CSF, IFN-$\gamma$ or untreated). Four arrays of donor BC974 were excluded because their values contradict their labels: its 0 h array looks stimulated (IRF1 10.0, CXCL10 8.4), its 0.5 and 1 h arrays look like 4 h or more of IFN-$\gamma$, and its 2 h array looks unstimulated; its 4, 6 and 24 h arrays agree with the other donors and were kept. The two deposited files disagree on two sections of chip 4676163034: the SOFT file labels section A as the 48 h untreated array of donor BC59 and section F as an array of another stimulus, and the series matrix swaps the two. We followed the series matrix (section F kept as the BC59 control, section A excluded), as section A looks unlike any untreated array (SOCS1 8.2, IRF1 8.3). This leaves 81 arrays, 35 stimulated and 46 untreated, from 35 donor labels on 24 chips, and $4 \times 81 = 324$ measurements (Table S7, Figure S1). The exclusions were proposed by the `orchestrating agent` from its audit of the deposit and decided by us during staging; they are judgements on the values, not confirmed labelling errors, and their effect on the results was not tested.

**Design.** The design is unbalanced. The early stimulated arrays (0.5 to 2 h) come from two donors, BC964 and BC985; untreated arrays exist at 0 h and from 6 h on; the time-matched untreated series at 6, 12, 24 and 48 h come from three donors, BC59, BC85-1 and BC85-2; 72 h holds seven pairs of a stimulated and an untreated array of the same donor; and the 27 arrays at 0 h come from 24 donor labels, 20 of which appear nowhere else. There are no technical replicates; the donors are the replicates.

Table S7: **Arrays per time point and arm** in the fitted data, identical in both runs. Each array contributes four measurements.

| time (h) | 0 | 0.5 | 1 | 2 | 4 | 6 | 12 | 24 | 48 | 72 |
|---|---|---|---|---|---|---|---|---|---|---|
| IFN-$\gamma$ | – | 2 | 2 | 2 | 3 | 6 | 3 | 10 | – | 7 |
| untreated | 27 | – | – | – | – | 3 | 3 | 3 | 3 | 7 |

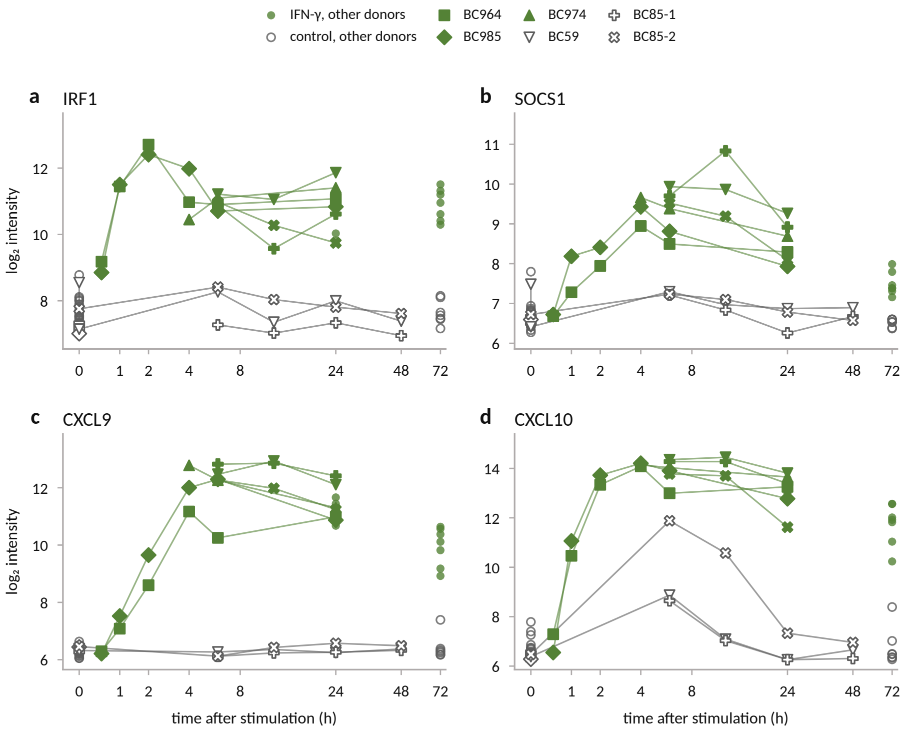

Figure S1: **The fitted data.** The 81 arrays, four transcripts, $\log_2$ quantile-normalised intensity against time; stimulated arrays green, untreated grey. Donors with a time course carry their own marker and are connected (BC964, BC985 and BC974 in the early series; BC59, BC85-1 and BC85-2 in the mid series); the arrays of the other donors are small circles. Untreated CXCL10 rises at 6 and 12 h in the three donors of the mid series. Time axis $\log(1 + t/2\,\mathrm{h})$.

**Observation and noise model.** Every transcript $i$ is observed as

$$y_{ir} = \log_2 m_i(t_r; u_r) + o_i + \varepsilon_{ir}, \qquad \varepsilon_{ir} \sim \mathcal{N}(0, \sigma_i^2), \tag{S2}$$

for array $r$ at time $t_r$, with $u_r = 1$ under IFN-$\gamma$ and $u_r = 0$ untreated, the modelled transcript level $m_i$ in units of its basal level ($m_i(0) = 1$), an offset $o_i$ for the probe's baseline and one residual standard deviation $\sigma_i$ per transcript, shared by both arms and constant in time. The offsets and $\sigma_i$ are estimated with the kinetic parameters and counted in the number of estimated parameters $n_\theta$. The `observation agent` first proposed a constant $\sigma$ for IRF1 and CXCL10 and one that depends on the predicted level for SOCS1 and CXCL9; at the first interaction point on the PEtab problems we sent it back with "use the same noise structure for all four transcripts: one constant sigma per transcript on the $\log_2$ scale; the noise must for the first run not depend on the model's predicted level". A detection or censoring term had been excluded during staging. At the maximum-likelihood estimate a free $\sigma_i$ equals the root-mean-square residual of transcript $i$ over all 81 arrays, so the $\pm 2\sigma$ bands of the main figure show what a model leaves unexplained: assay noise, donor-to-donor differences (no donor parameters were fitted) and misfit. Because one $\sigma_i$ serves both arms, it is a compromise: in model 5 the root-mean-square residual of CXCL9 is 0.63 in the stimulated and 0.20 in the untreated arm, against $\sigma = 0.44$.

**Replicate floor.** The engine's model-free reference for the noise is the replicate floor, the pooled standard deviation of the arrays that share stimulus and time point (14 such cells, 67 degrees of freedom per transcript in run 1): 0.49 (IRF1), 0.36 (SOCS1), 0.46 (CXCL9) and 0.70 (CXCL10) $\log_2$ units. Since the arrays of a cell come from different donors, the floor contains the donor-to-donor spread.

#### S3.2 Inputs to the runs

The first run received the data file, the project description and the intent, which we wrote with the `orchestrating agent` from our notes, the deposit and the Methods of the paper. The description is a lab note on the cells and the stimulation, the measurement and the excluded arrays, the measured curves of both arms, and our requests for the model: one effective activity downstream of IFN-$\gamma$ that drives the four transcripts, each with basal and activity-dependent production and first-order degradation and without freely estimated Hill exponents; kinetic parameters shared across donors; and no change in untreated expression attributed to stimulation-induced feedback. The only literature was three claims we wrote from the paper (the dose, the differentiation and the processing of the values); no agent searched online, and the knowledge module ran offline, so no evidence floor applied, and we kept its two reactions for the activity $A$ in the base model. No condition was held out, and every candidate was estimated with 300 starts (Table S3). The intent, in its relevant parts:

> **Question.** Can one shared transcriptional activity driven by IFN-$\gamma$, together with gene-specific basal production, activity-dependent production and first-order mRNA degradation, explain the IRF1, SOCS1, CXCL9 and CXCL10 time courses of GM-CSF-differentiated human macrophages over 0.5–72 h, or is an attenuation of that activity needed, and does a single inhibitory feedback variable (for instance one produced from the SOCS1 transcript) account for it? Which of the activation, turnover and feedback rates can these arrays actually pin down?
>
> **Required reactions.** For each transcript $X$ of IRF1, SOCS1, CXCL9 and CXCL10: $\varnothing \to X$, basal and $A$-dependent production with $A$ as modifier, linear or simply saturating in $A$, no free

Hill exponent; and $X \to \varnothing$, first-order degradation. The activation and loss of $A$ are left open on purpose.

**Success criteria.** (1) The no-feedback base (shared activity; per transcript a basal rate, an activity-dependent rate and a degradation rate) and each alternative are compared by information criteria on all 81 arrays; a winner is reported with Akaike weight $\geq 0.7$, otherwise the indistinguishable set is reported as such. (2) The selected model reproduces each of the four IFN-$\gamma$ time courses within the donor-to-donor scatter, with no systematic time-dependent residual for any transcript. (3) The initial state is constrained by the pretreatment arrays and the untreated time courses are fitted within the donor-to-donor scatter; movement in the untreated cultures is never used as evidence for or against stimulation-induced feedback, and any control excursion the model does not reproduce is reported as unexplained rather than absorbed by a feedback parameter. (4) If attenuation is selected, its timescale and the four degradation rates are reported with profile-likelihood intervals; every rate the arrays cannot resolve is named as such. (5) Kinetic parameters are shared across donors; any donor-specific baseline or scaling parameter is declared and counted against the model.

**Inference target.** Mechanism discrimination.

Before run 1, two launches stopped before any fit. In the second, the `construction agent` proposed a transient after handling that acts through $A$, to reproduce the rise in the untreated cultures, which the draft of criterion 3 asked to be reproduced. When we asked which criterion forbade attributing this movement to feedback, the `orchestrating agent` proposed the wording of criterion 3 given above, which we adopted.

#### S3.3 The first run: five rounds and 26 fitted models

Table S8 lists every model candidate of run 1 in build order with the values the run recorded, and Figure S2 the standardised residuals of the six models of the main text. All models are fitted to the same 324 measurements with the same observation model, so their likelihoods and information criteria are comparable across rounds.

Table S8: **The model candidates of run 1, in build order, and the recalibration of run 2.** Round, candidate with what it adds to the base of its round, estimated parameters $n_\theta$, negative log-likelihood, AIC and BIC ($= 2\,\mathrm{NLLH} + n_\theta \ln 324$), $\Delta$AIC to the round's criterion winner and Akaike weight $w$ within the round, and the number of starts (of those with a finite final value) within 0.1 of the best. From round 2 on, a round's base is the previous round's model with its fit reused. The criterion winner of each round is the model of the main text, except in round 4, where it is the CXCL9 $\sigma$ split (italic) and model 4 was taken forward instead. Run 2 has no selection; its AIC and BIC are computed from its likelihood.

| round | candidate | $n_\theta$ | NLLH | AIC | BIC | $\Delta$AIC | $w$ | starts |
|---|---|---|---|---|---|---|---|---|
| 1 | no feedback: the base model (**model 0**) | 17 | 369.63 | 773.27 | 837.54 | 113.40 | <0.001 | 300/300 |
| 1 | generic attenuation: $R$, induced by $A$, divides the generation of $A$ (**model 1**) | 19 | 310.93 | 659.86 | 731.70 | 0.00 | >0.999 | 300/300 |
| 1 | stimulus loss: first-order loss of the stimulus | 18 | 320.73 | 677.46 | 745.51 | 17.60 | <0.001 | 231/300 |

| round | candidate | $n_\theta$ | NLLH | AIC | BIC | ΔAIC | $w$ | starts |
|---|---|---|---|---|---|---|---|---|
| 1 | SOCS1-transcript feedback: an inhibitor made from the SOCS1 transcript divides the generation of $A$ | 19 | 325.70 | 689.41 | 761.24 | 29.54 | <0.001 | 300/300 |
| 2 | base: model 1, fit reused | 19 | 310.93 | 659.86 | 731.70 | 43.51 | <0.001 | reused |
| 2 | IRF1 saturation and onset lag $A_2$ (**model 2**) | 22 | 286.18 | 616.35 | 699.53 | 0.00 | 0.881 | 298/299 |
| 2 | IRF1 saturation, onset lag and fast second attenuator | 24 | 286.18 | 620.36 | 711.09 | 4.00 | 0.119 | 299/300 |
| 2 | onset lag: the chemokines driven by a delayed copy $A_2$ of $A$ | 21 | 297.06 | 636.11 | 715.51 | 19.76 | <0.001 | 297/299 |
| 2 | IRF1 saturation: IRF1 production saturating in $A$ | 20 | 299.49 | 638.97 | 714.59 | 22.62 | <0.001 | 300/300 |
| 2 | onset lag and fast second attenuator | 23 | 297.06 | 640.12 | 727.07 | 23.76 | <0.001 | 299/300 |
| 2 | IRF1 saturation and fast second attenuator | 22 | 299.49 | 642.97 | 726.15 | 26.62 | <0.001 | 300/300 |
| 2 | fast second attenuator | 21 | 310.93 | 663.86 | 743.26 | 47.51 | <0.001 | 300/300 |
| 3 | base: model 2, fit reused | 22 | 286.18 | 616.35 | 699.53 | 41.09 | <0.001 | reused |
| 3 | culture component: a transient $H$ after handling drives CXCL10 production in both arms (**model 3**) | 24 | 263.63 | 575.26 | 666.00 | 0.00 | 0.990 | 299/299 |
| 3 | delayed-onset attenuator: $R$ produced from a delayed copy of $A$ | 24 | 268.75 | 585.50 | 676.24 | 10.24 | 0.006 | 169/300 |
| 3 | SOCS1-transcript feedback: an inhibitor made from the SOCS1 transcript removes $A$ | 24 | 269.12 | 586.25 | 676.98 | 10.98 | 0.004 | 178/300 |
| 3 | IRF1 negative autoregulation: an IRF1-induced factor destabilises IRF1 mRNA | 24 | 275.55 | 599.10 | 689.84 | 23.84 | <0.001 | 299/299 |
| 4 | base: model 3, fit reused | 24 | 263.63 | 575.26 | 666.00 | 55.39 | <0.001 | reused |
| 4 | *CXCL9 $\sigma$ per arm (observation model)* | 25 | 234.94 | 519.88 | 614.40 | 0.00 | >0.999 | 293/299 |
| 4 | activity-dependent signal loss: the availability $L$ is consumed in proportion to $A$ (**model 4**) | 25 | 245.46 | 540.91 | 635.43 | 21.04 | <0.001 | 295/300 |
| 4 | delayed-onset attenuator, added back by us | 26 | 245.90 | 543.81 | 642.11 | 23.93 | <0.001 | 275/299 |
| 4 | SOCS1-protein feedback: a SOCS1 protein state made from the transcript removes $A$ | 26 | 245.93 | 543.86 | 642.16 | 23.98 | <0.001 | 183/300 |
| 4 | IRF1 destabiliser: an $A$-induced factor shortens the IRF1 mRNA's life | 26 | 248.87 | 549.74 | 648.04 | 29.86 | <0.001 | 297/299 |
| 4 | SOCS1 $\sigma$ per arm (observation model) | 25 | 257.47 | 564.93 | 659.45 | 45.06 | <0.001 | 299/299 |
| 5 | base: model 4, fit reused | 25 | 245.46 | 540.91 | 635.43 | 1.91 | 0.222 | reused |
| 5 | upstream stage: a first-order stage $S$ between the stimulus and $A$ (**model 5**) | 27 | 242.50 | 539.00 | 641.08 | 0.00 | 0.577 | 134/300 |
| 5 | cumulative signal loss, re-proposed (a second rate of the same loss) | 26 | 245.46 | 542.91 | 641.21 | 3.91 | 0.082 | 288/300 |
| 5 | delayed-onset attenuator | 27 | 244.87 | 543.74 | 645.82 | 4.74 | 0.054 | 17/300 |
| 5 | SOCS1-protein deactivation of $A$ | 27 | 245.27 | 544.54 | 646.62 | 5.54 | 0.036 | 128/300 |
| 5 | induced fast deactivator of $A$ | 27 | 245.45 | 544.89 | 646.97 | 5.89 | 0.030 | 296/300 |

| round | candidate | $n_\theta$ | NLLH | AIC | BIC | ΔAIC | $w$ | starts |
|---|---|---|---|---|---|---|---|---|
| run 2 | model 5, recalibrated (S3.5) | 25 | 241.75 | 533.50 | 628.02 | – | – | 79/300 |

**Round 1.** The `construction agent` wrote the base model with our eight required reactions and the two essential reactions of the activity $A$ (eight species and 15 reactions in its model text, with the terms of the alternatives switched off; model 0 has five coupled states, S3.4) and suggested STAT1 as a possible counterpart of $A$ without naming $A$ as STAT1. We approved the base. The `expansion agent` wrote three alternatives as settings of the base's switches: an inhibitor made from the SOCS1 transcript, a generic attenuator $R$ induced by $A$, and first-order loss of the stimulus. The generic attenuator won decisively (model 1); no attenuation was 113.4 AIC units behind, the SOCS1-transcript feedback 29.5 and stimulus loss 17.6. Of model 1's gain of 58.7 log-likelihood units over model 0, 37.3 came from the stimulated arm; IRF1 was the exception, fitted worse than by model 0 (by 6.7 units in the stimulated and 2.9 in the untreated arm). Three of six misfit indicators fired: $\chi^2$/dof 1.82 against the replicate floor, and the fitted $\sigma$ of IRF1 and CXCL10 at 1.36 and 1.45 times the floor. The attenuator ended at a corner of its box, its gain $g_R$ at the upper and its relaxation rate $k_R$ at the lower bound, with the gradient pointing outward, so that $R$ acted as an integrator of $A$; the IRF1 degradation rate was at its upper bound. The residuals showed IRF1 below the model between 4 and 12 h and the chemokines starting later than the model (CXCL10 $-3.2\sigma$ at 0.5 h). The `interpretation agent` named IRF1 the prime suspect and recommended a saturating dependence of IRF1 on $A$, with an onset delay second. The `revision agent` proposed IRF1 production saturating in $A$, a fast second attenuator, a delayed copy $A_2$ of $A$ driving the chemokines, and a $\sigma$ per arm for CXCL9 and for CXCL10 (its first reply named a reaction the model does not have and was returned; the redraft repaired it). We asked for every combination of the changes rather than one at a time; under the ceiling of eight models this admits three changes, so the two noise changes were left out.

**Round 2.** All eight combinations of the three changes were fitted on model 1. IRF1 saturation with the onset lag won (model 2, weight 0.88), 24.8 log-likelihood units ahead of model 1, 16.7 of them in the stimulated arm (IRF1 8.1, CXCL9 5.3, CXCL10 3.3); either change alone was worse, by 19.8 AIC units without the saturation and 22.6 without the lag. The fast second attenuator came out degenerate, both of its parameters at their lower bounds, and gained nothing. Two indicators still fired ($\chi^2$/dof 1.59; the $\sigma$ of CXCL10 at 1.40 times the floor). The IRF1 residual had become a transient excess near 1.5 h ($1.9\sigma$), and untreated CXCL10 showed a transient near 6.6 h ($3.8\sigma$) that the flat untreated trajectories of models 0 to 2 cannot follow; the attenuator stayed at its corner in all eight models. The `interpretation agent` proposed freeing the attenuator from its box. The `revision agent` read the corner as the fit moving towards an integrating feedback, but since a revision cannot change a bound, it flagged this to us, and we kept it for a later run (S3.5). It proposed four changes: SOCS1-transcript feedback on model 2, an attenuator with a delayed onset, a transient after handling at $t = 0$ that drives CXCL10 production only and never $A$, aimed at untreated CXCL10, and an IRF1-induced factor for the excess at 2 h; a fifth, a second delay stage for the chemokines, named a reaction the model does not have and was withdrawn in the redraft.

We bought all four, fitted one at a time.

**Round 3.** The culture component won (model 3, weight 0.99): 21.3 of its 22.5 log-likelihood units over model 2 came from untreated CXCL10. The delayed-onset attenuator and the SOCS1-transcript feedback gained 11.7 and 11.5 units in the stimulated arm, against 1.3 for the culture component, and lost in the one cell they cannot fit without it: in untreated CXCL10 each trailed by about 21.5 units, more than the about 16 by which each led in all other cells together. The IRF1 change improved IRF1 only in a degenerate corner of its parameters. One indicator still fired ($\chi^2$/dof 1.37). The `interpretation agent` wrote that the selected model "wins on the untreated cultures, not on the IFN-$\gamma$ response", and that its weight said nothing about attenuation or feedback, which had not been combined with the culture component. The `revision agent` read the attenuator pinned at its corner as a mechanism in itself, a cumulative activity-dependent loss of signalling availability, such as receptor down-regulation or ligand consumption, with one parameter, and proposed it beside SOCS1-protein feedback, an IRF1 destabiliser driven by $A$, and a $\sigma$ per arm for CXCL9 (the change the ceiling had left out after round 1) and for SOCS1. It did not propose the delayed-onset attenuator again, whose gain it read as the same late decline; we added it back, so that the attenuation forms were compared on model 3.

**Round 4.** The engine first dropped the delayed-onset attenuator we had added back, as already carried by the base, because model 3's model text contains it switched off. We stopped at the next interaction point and had the engine's rule corrected (a change that the base's parameter table holds at its neutral value now counts as absent), and the round was re-entered at M1 with all seven models; the run record carries the code state of the correction. The CXCL9 $\sigma$ split won (weight >0.999), with 22.4 of its 28.7 log-likelihood units over model 3 from untreated CXCL9, where the arrays sit at the array's floor: the split gave the untreated arm a $\sigma$ of 0.20, 0.43 times the replicate floor, and the stimulated arm 0.71. The gain came from the smaller $\sigma$ of the untreated arm, not from a better curve, and the `assembly verification agent` had warned that the split makes the values the lab note calls the array's floor the most precise CXCL9 data in the fit. The four mechanisms each gained 11.8 to 12.3 units in the stimulated arm, without the split, and the three attenuation forms (activity-dependent signal loss, delayed-onset attenuator, SOCS1-protein feedback) lay within 0.5 units of each other; signal loss was the first candidate of the run whose attenuator parameters both left their bounds. The `interpretation agent` wrote that the winner "is not a mechanism" but "a detection-floor artefact already named in the lab note", and that the ranking below it reflected the search order (one change at a time on each round's base, 18 of 4096 possible models), and recommended continuing from the winner. The `orchestrating agent` proposed instead to take signal loss, model 4, as the base of round 5, with the other attenuation forms, an onset delay and the $\sigma$ split as changes, then to compute profiles and close the run. We authorised this plan, and the `orchestrating agent` entered the remaining answers of run 1 under it.

**Round 5.** The `revision agent` treated the $\sigma$ split as bookkeeping and found its prime suspect where no round had looked: in every model the stimulus had entered the generation of $A$ as an instantaneous step, and it read the residuals at 0.5 to 2 h as about 0.4 h of dead time followed by a steep rise. It proposed an upstream stimulus stage, re-proposed cumulative signal loss, and added a SOCS1-protein deactivation of $A$ and an induced fast deactivator of $A$ for its early peak; under the plan, the delayed-onset attenuator was added back once more. The engine did not take up the $\sigma$ split as a change, because it had been selected on the round-4 winner, so all six models of round 5 have one $\sigma$ for CXCL9. On model 4 the other attenuation forms added 0.0 to 0.6 log-likelihood units, and the re-proposed signal loss, model 4's own mechanism under a new parameter name, reproduced model 4 exactly. The upstream stage gained 3.0 units, 2.7 of them in stimulated CXCL10 at a cost of 1.7 in stimulated SOCS1: weight 0.58, 1.91 AIC units ahead of model 4 and indistinguishable from it; BIC prefers model 4 (635.4 against 641.1). At the verdict one indicator still fired ($\chi^2$/dof 1.22 against the replicate floor, $p = 0.005$), every fitted $\sigma$ lay at 0.96 to 1.17 times the floor, and the IRF1 residual was the same transient excess near 1.5 h that models 2 to 4 had shown (Figure S2); four proposals had aimed at it, and none removed it. Profiles of model 5 left the attenuator's relaxation rate open at its lower bound and its gain across the whole box. The `interpretation agent` concluded that "the data therefore demand attenuation but give no evidence that the attenuating agent is SOCS1" and recommended one more change, saturating chemokine production with a fast deactivator of $A$; the `orchestrating agent` closed the run under the plan we had authorised in round 4.

#### S3.4 The models

**Conventions.** Time is in hours, $t \in [0, 72]$, and all states are dimensionless. The input $u$ is 1 in the stimulated and 0 in the untreated condition, which also holds the 27 arrays at 0 h. Each transcript $m_i$, $i \in \{\mathrm{IRF1}, \mathrm{SOCS1}, \mathrm{CXCL9}, \mathrm{CXCL10}\}$, is in units of its basal level. The initial state is the unstimulated steady state, the same in both arms: $m_i(0) = 1$, the availability $L(0) = 1$, the first handling state $H_1(0) = 1$, and every other state 0. Within a round all candidates are compiled from one model text that holds the reactions of every change of the round; a candidate is a row of its parameter table, which fixes the gain of each mechanism it lacks at its neutral value 0, so the model texts in the record must be read together with the parameter tables. Kinetic parameters are estimated on a $\log_{10}$ scale except the fractions $w_{\mathrm{lag}}$ and $w_S$ (linear, in $[0, 1]$); the offsets $o_i \in [-10^3, 10^3]$ (linear) and $\sigma_i \in [10^{-4}, 10^3]$ ($\log_{10}$) enter through eq. (S2). Table S9 gives the fitted values.

**Model 0, the base model** (round 1, `M0_no_feedback`). With the alternatives switched off, the generation of $A$ reduces to $k_A u$, and

$$\dot{A} = k_A u - k_A A, \tag{S3}$$

$$\dot{m}_i = d_i\,(1 + f_i A) - d_i\,m_i, \tag{S4}$$

with $A(0) = 0$: five coupled states and 17 estimated parameters ($k_A$, the four $d_i$ and $f_i$, the four $o_i$ and $\sigma_i$). Equal production and loss coefficients fix the scale of $A$, whose unattenuated stimulated steady state is 1, so the $f_i$ are folds of basal production per unit $A$. Eq. (S3) holds the two essential reactions of the knowledge module, eq. (S4) our eight required ones.

**Models 1 to 5: the equations that change.** Each model is given as its difference from the previous one; all other equations are unchanged.

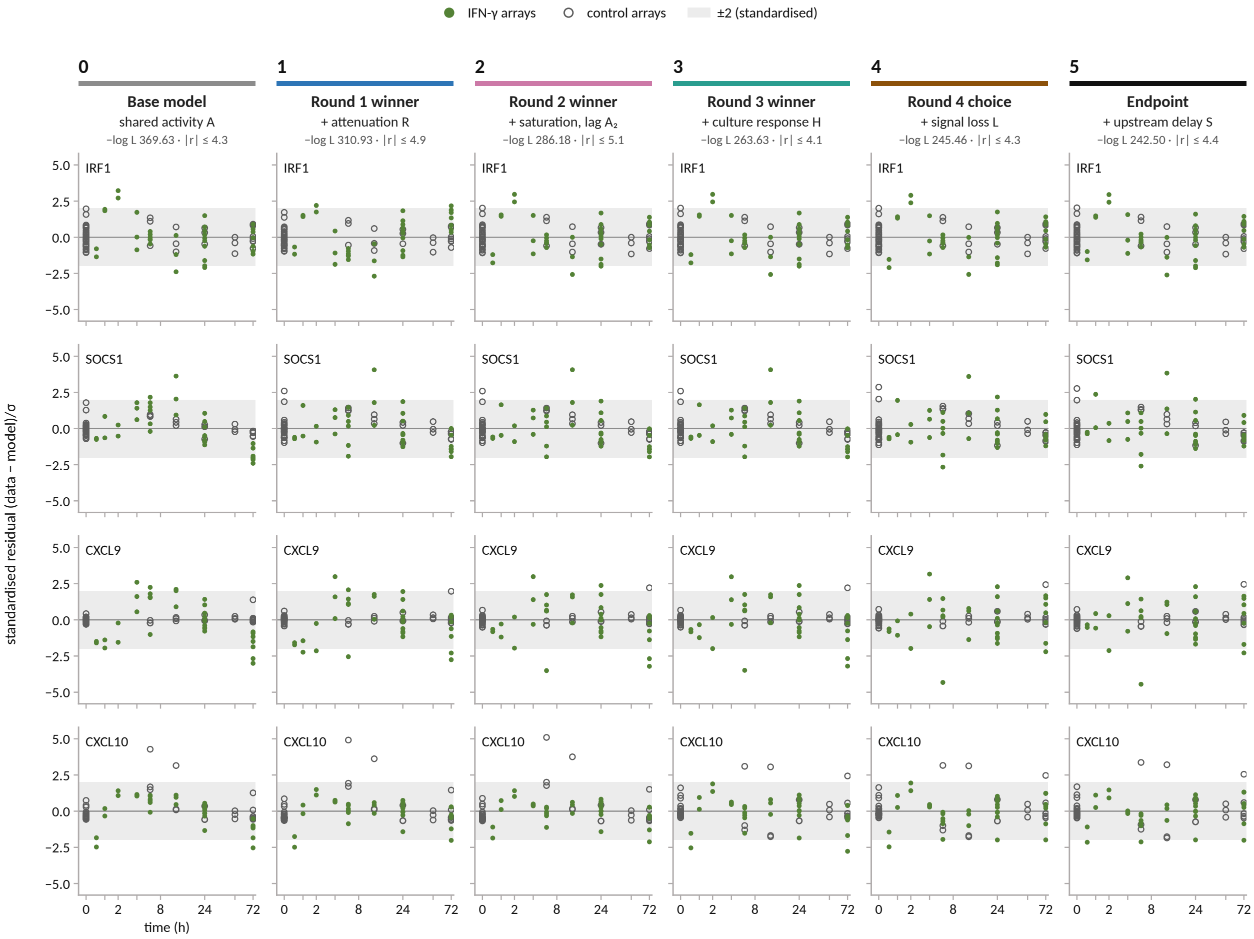

Figure S2: **Standardised residuals of models 0 to 5.** (data − model)$/\sigma_i$ at the measured times, with each model's fitted $\sigma_i$; one column per model of the main text, headed by its number, the round it came from, what it added, its negative log-likelihood and its largest absolute residual; stimulated arrays filled, untreated open; grey band $\pm 2$. The systematic misfit under IFN-$\gamma$ shrinks with the attenuation (model 1) and with the saturation and lag (model 2), and the untreated CXCL10 misfit at 6 and 12 h with the culture component (model 3); the IRF1 excess at 1 and 2 h remains in every column. The recalibration of model 5 in run 2 changes no standardised residual by more than 0.07. Time axis $\log(1 + t/2\,\mathrm{h})$.

**Model 1** (round 1, `M2_generic_A_induced_attenuation`). The attenuator $R$, induced by $A$, divides the generation of $A$:

$$\dot{A} = k_A \frac{u}{1 + g_R R} - k_A A, \qquad \dot{R} = k_R A - k_R R, \quad R(0) = 0. \tag{S5}$$

Added: $g_R \in [0.01, 100]$ and $k_R \in [0.01, 10]\,\mathrm{h}^{-1}$. Both ended at the corner, $g_R = 100$ and $k_R = 0.01\,\mathrm{h}^{-1}$, with $d_{\mathrm{IRF1}} = 10\,\mathrm{h}^{-1}$ at its upper bound, all three with the gradient pointing outward. Then $g_R R = g_R k_R \int_0^t e^{-k_R(t-s)} A(s)\,\mathrm{d}s$, which to first order in $k_R t$ ($\leq 0.72$ over the 72 h) depends on $g_R$ and $k_R$ only through $g_R k_R = 1.0\,\mathrm{h}^{-1}$: the integrator limit.

**Model 2** (round 2). IRF1 production saturates in $A$, and the chemokines are driven by a mixture of $A$ and a delayed copy $A_2$:

$$d_{\mathrm{IRF1}}\,(1 + f_{\mathrm{IRF1}} A) \;\to\; d_{\mathrm{IRF1}}\Big(1 + \frac{f_{\mathrm{IRF1}}\,A}{1 + h_{\mathrm{IRF1}} A}\Big), \tag{S6}$$

$$d_i\,(1 + f_i A) \;\to\; d_i\big(1 + f_i\,[(1 - w_{\mathrm{lag}})A + w_{\mathrm{lag}} A_2]\big), \qquad i \in \{\mathrm{CXCL9}, \mathrm{CXCL10}\}, \tag{S7}$$

$$\dot{A}_2 = k_{\mathrm{lag}} A - k_{\mathrm{lag}} A_2, \quad A_2(0) = 0. \tag{S8}$$

Added: $h_{\mathrm{IRF1}} \in [10^{-3}, 10^3]$, $w_{\mathrm{lag}} \in [0, 1]$ and $k_{\mathrm{lag}} \in [0.05, 20]\,\mathrm{h}^{-1}$. $h_{\mathrm{IRF1}} = 15.9$ and $k_{\mathrm{lag}} = 0.754\,\mathrm{h}^{-1}$ ended inside their boxes; $w_{\mathrm{lag}} = 1$ at its upper bound, so the chemokines follow $A_2$ alone; $g_R$ and $k_R$ stayed at the corner, and $d_{\mathrm{IRF1}} = 2.33\,\mathrm{h}^{-1}$ left its bound.

**Model 3** (round 3, `handling_pulse_cxcl10`). A two-stage transient, started at $t = 0$ in both arms and independent of $u$, adds to the production of CXCL10 only:

$$\dot{m}_{\mathrm{CXCL10}} = d_{\mathrm{CXCL10}}\big(1 + f_{\mathrm{CXCL10}}\,[(1 - w_{\mathrm{lag}})A + w_{\mathrm{lag}} A_2] + f_H H_2\big) - d_{\mathrm{CXCL10}}\, m_{\mathrm{CXCL10}},$$

$$\dot{H}_1 = -k_H H_1, \qquad \dot{H}_2 = k_H H_1 - k_H H_2, \qquad H_1(0) = 1,\; H_2(0) = 0, \tag{S9}$$

so that $H_2(t) = k_H t\, e^{-k_H t}$ peaks at $t = 1/k_H$. Added: $f_H \in [0.01, 10^3]$ and $k_H \in [0.02, 2]\,\mathrm{h}^{-1}$, which ended inside: $f_H = 22.4$, $k_H = 0.317\,\mathrm{h}^{-1}$ (peak at 3.2 h). $g_R$, $k_R$ and $w_{\mathrm{lag}}$ remained at their bounds.

**Model 4** (round 4, `activity_dependent_signal_loss`). The availability $L$, constant until now, is consumed in proportion to the activity:

$$\dot{L} = -k_{LA}\, A L, \quad L(0) = 1, \qquad \dot{A} = k_A \frac{u\,L}{1 + g_R R} - k_A A, \tag{S10}$$

so that $L(t) = \exp\big(-k_{LA} \int_0^t A\,\mathrm{d}s\big)$. Added: $k_{LA} \in [10^{-4}, 1]$ per unit $A$ and hour, which ended at 0.888, near its upper bound. In model 4 both attenuator parameters left their bounds, $g_R = 86.9$ and $k_R = 0.0509\,\mathrm{h}^{-1}$; $w_{\mathrm{lag}} = 1$ remained at its bound.

**Model 5** (round 5, `upstream_stimulus_delay`). A first-order stage $S$ (`S_in` in the record) is put between the input and $A$:

$$\dot{S} = k_{\mathrm{Sin}}\, u - k_{\mathrm{Sin}} S, \quad S(0) = 0,$$

$$k_A \frac{u\,L}{1 + g_R R} \;\to\; k_A \frac{[(1 - w_S)\,u + w_S S]\,L}{1 + g_R R}. \tag{S11}$$

Added: $w_S \in [0, 1]$ and $k_{\mathrm{Sin}} \in [0.3, 30]\,\mathrm{h}^{-1}$. $w_S = 1$ ended at its upper bound, so $A$ is driven by $S$ alone; $k_{\mathrm{Sin}} = 1.67\,\mathrm{h}^{-1}$ inside. In model 5, $g_R = 100$, $k_{LA} = 1.00$, $w_{\mathrm{lag}} = 1$ and $w_S = 1$ are at their upper bounds, $k_A = 0.102\,\mathrm{h}^{-1}$ lies within 0.3 % of its box above the lower bound, and $k_R = 0.120\,\mathrm{h}^{-1}$ is inside.

**Model 5 in full** (run 1; the candidate's model text with its parameter table):

$$
\begin{aligned}
\dot S &= k_{\mathrm{Sin}}\,(u - S),\\
\dot A &= k_A\,\frac{[(1-w_S)\,u + w_S S]\,L}{1+g_R R} - k_A A,\\
\dot L &= -k_{LA}\,AL,\\
\dot R &= k_R\,(A - R),\\
\dot A_2 &= k_{\mathrm{lag}}\,(A - A_2),\\
\dot m_{\mathrm{IRF1}} &= d_{\mathrm{IRF1}}\Big(1 + \frac{f_{\mathrm{IRF1}}\,A}{1+h_{\mathrm{IRF1}}A} - m_{\mathrm{IRF1}}\Big),\\
\dot m_{\mathrm{SOCS1}} &= d_{\mathrm{SOCS1}}\big(1 + f_{\mathrm{SOCS1}}A - m_{\mathrm{SOCS1}}\big),\\
\dot m_{\mathrm{CXCL9}} &= d_{\mathrm{CXCL9}}\big(1 + f_{\mathrm{CXCL9}}\,[(1-w_{\mathrm{lag}})A + w_{\mathrm{lag}}A_2] - m_{\mathrm{CXCL9}}\big),\\
\dot m_{\mathrm{CXCL10}} &= d_{\mathrm{CXCL10}}\big(1 + f_{\mathrm{CXCL10}}\,[(1-w_{\mathrm{lag}})A + w_{\mathrm{lag}}A_2] + f_H H_2 - m_{\mathrm{CXCL10}}\big),\\
\dot H_1 &= -k_H H_1, \qquad \dot H_2 = k_H\,(H_1 - H_2),
\end{aligned}
\tag{S12}
$$

with $S = A = R = A_2 = H_2 = 0$, $L = H_1 = 1$ and $m_i = 1$ at $t = 0$: 11 coupled states, with 19 kinetic and 8 observation parameters estimated (27). The model text of round 5 holds 20 species and 45 reactions; the other nine species are inert in model 5, because every rate through which one of them could act has its gain fixed at 0, and no observable depends on them.

#### S3.5 The second run: recalibration of model 5 and its profiles

**Why a second run.** From round 1 on, the attenuator sat at a corner of its box ($g_R$ at the upper, $k_R$ at the lower bound), and in model 5 $g_R$ and $k_{LA}$ were still at their upper bounds. A revision cannot change a bound, so the `revision agent` flagged it to us after round 2, and we kept it for a later run. After run 1 we asked the `orchestrating agent` what a new run should do, and whether one was worth doing at all. It recommended one focused run that measures model 5 properly, with the attenuator reparameterised and the boxes widened, rather than extending it, and we agreed.

**What was supplied.** Model 5's reactions were supplied as the model, without the switched-off branches (11 species, 20 reactions), and changed in three ways: (i) $R \to g_R R$, a rescaling of the state, so that $\dot R = G_R A - k_R R$ and the inhibition is $1/(1+R)$, with $G_R = g_R k_R$; (ii) $F_{\mathrm{IRF1}} = f_{\mathrm{IRF1}}/h_{\mathrm{IRF1}}$, so that IRF1 production tends to $d_{\mathrm{IRF1}}(1 + F_{\mathrm{IRF1}})$ at saturating $A$; (iii) $w_S = w_{\mathrm{lag}} = 1$, their fitted values in model 5, fixed and removed. (i) and (ii) are exact reparameterisations; (iii) and the

Table S9: **Fitted values of models 0 to 5 (run 1) and of model 5 recalibrated (run 2).** Rounded best estimates; $^\dagger$ at a bound; $^\ddagger$ within 5 % of the box edge on the parameter's scale; $[x]$ fixed; – absent. Bracketed entries of the $G_R$ and $F_{\mathrm{IRF1}}$ rows in run 1 are the combinations $g_R k_R$ and $f_{\mathrm{IRF1}}/h_{\mathrm{IRF1}}$, which run 2 estimates directly (S3.5). Rates in $\mathrm{h}^{-1}$; $k_{LA}$ and $G_R$ per unit $A$ and hour; gains dimensionless; $o_i$ and $\sigma_i$ in $\log_2$ units. Run 2's AIC is computed from its likelihood.

| model | 0 | 1 | 2 | 3 | 4 | 5 | 5, recalibrated |
|---|---|---|---|---|---|---|---|
| $k_{\mathrm{Sin}}$ | – | – | – | – | – | 1.669 | $0.300^\dagger$ |
| $k_A$ | 1.107 | 0.193 | 0.215 | 0.215 | 0.191 | $0.102^\ddagger$ | 0.107 |
| $k_{LA}$ | – | – | – | – | $0.888^\ddagger$ | $1.00^\dagger$ | 4.238 |
| $g_R$ | [0] | $100^\dagger$ | $100^\dagger$ | $100^\dagger$ | $86.91^\ddagger$ | $100^\dagger$ | – |
| $k_R$ | [0.3] | $0.0100^\dagger$ | $0.0100^\dagger$ | $0.0100^\dagger$ | 0.0509 | 0.120 | 0.0557 |
| $G_R$ $[g_R k_R]$ | – | (1.000) | (1.000) | (1.000) | (4.426) | (12.03) | 209.1 |
| $k_{\mathrm{lag}}$ | – | – | 0.754 | 0.862 | 0.192 | 1.200 | 1.090 |
| $w_{\mathrm{lag}}$ | – | – | $1^\dagger$ | $1^\dagger$ | $1^\dagger$ | $1^\dagger$ | [1] |
| $w_S$ | – | – | – | – | – | $1^\dagger$ | [1] |
| $d_{\mathrm{IRF1}}$ | $7.777^\ddagger$ | $10^\dagger$ | 2.330 | 2.330 | 1.439 | 2.635 | 2.643 |
| $d_{\mathrm{SOCS1}}$ | 1.350 | 0.823 | 0.687 | 0.686 | 0.176 | 0.569 | 0.562 |
| $d_{\mathrm{CXCL9}}$ | 0.373 | 0.245 | 0.541 | 0.491 | 0.488 | 0.371 | 0.383 |
| $d_{\mathrm{CXCL10}}$ | 0.589 | 0.433 | 1.945 | 1.275 | 1.341 | 1.457 | 1.508 |
| $f_{\mathrm{IRF1}}$ | 9.291 | 40.41 | 189.6 | 189.5 | 1357 | 1393 | – |
| $h_{\mathrm{IRF1}}$ | – | – | 15.91 | 15.90 | 126.0 | 130.1 | 581.0 |
| $F_{\mathrm{IRF1}}$ $[f_{\mathrm{IRF1}}/h_{\mathrm{IRF1}}]$ | – | – | (11.92) | (11.92) | (10.76) | (10.71) | 10.79 |
| $f_{\mathrm{SOCS1}}$ | 2.767 | 13.39 | 13.45 | 13.45 | 48.12 | 50.93 | 240.4 |
| $f_{\mathrm{CXCL9}}$ | 30.95 | 153.7 | 156.8 | 156.6 | 597.5 | 627.8 | 2958 |
| $f_{\mathrm{CXCL10}}$ | 74.73 | 362.7 | 370.0 | 456.2 | 1775 | 1830 | $8639^\ddagger$ |
| $f_H$ | – | – | – | 22.43 | 22.30 | 20.59 | 20.44 |
| $k_H$ | – | – | – | 0.317 | 0.318 | 0.304 | 0.302 |
| $o_{\mathrm{IRF1}}$ | 7.618 | 7.636 | 7.616 | 7.616 | 7.616 | 7.616 | 7.616 |
| $o_{\mathrm{SOCS1}}$ | 6.707 | 6.690 | 6.691 | 6.691 | 6.705 | 6.714 | 6.713 |
| $o_{\mathrm{CXCL9}}$ | 6.295 | 6.292 | 6.304 | 6.304 | 6.310 | 6.317 | 6.318 |
| $o_{\mathrm{CXCL10}}$ | 6.926 | 6.923 | 6.921 | 6.602 | 6.604 | 6.603 | 6.603 |
| $\sigma_{\mathrm{IRF1}}$ | 0.590 | 0.665 | 0.576 | 0.576 | 0.580 | 0.571 | 0.570 |
| $\sigma_{\mathrm{SOCS1}}$ | 0.610 | 0.427 | 0.428 | 0.428 | 0.382 | 0.392 | 0.389 |
| $\sigma_{\mathrm{CXCL9}}$ | 0.789 | 0.556 | 0.489 | 0.490 | 0.443 | 0.437 | 0.437 |
| $\sigma_{\mathrm{CXCL10}}$ | 1.157 | 1.008 | 0.973 | 0.735 | 0.723 | 0.700 | 0.698 |
| $n_\theta$ | 17 | 19 | 22 | 24 | 25 | 27 | 25 |
| NLLH | 369.63 | 310.93 | 286.18 | 263.63 | 245.46 | 242.50 | 241.75 |
| AIC | 773.27 | 659.86 | 616.35 | 575.26 | 540.91 | 539.00 | 533.50 |

widened boxes (Table S10) change the estimation problem. The equations are

$$
\begin{aligned}
\dot S &= k_{\mathrm{Sin}}\, u - k_{\mathrm{Sin}} S,\\
\dot A &= k_A\, \frac{S\,L}{1+R} - k_A A,\\
\dot L &= -k_{LA}\, AL,\\
\dot R &= G_R A - k_R R,\\
\dot A_2 &= k_{\mathrm{lag}} A - k_{\mathrm{lag}} A_2,\\
\dot m_{\mathrm{IRF1}} &= d_{\mathrm{IRF1}}\Big(1 + F_{\mathrm{IRF1}}\, \frac{h_{\mathrm{IRF1}} A}{1+h_{\mathrm{IRF1}} A}\Big) - d_{\mathrm{IRF1}}\, m_{\mathrm{IRF1}},\\
\dot m_{\mathrm{SOCS1}} &= d_{\mathrm{SOCS1}}\,(1 + f_{\mathrm{SOCS1}} A) - d_{\mathrm{SOCS1}}\, m_{\mathrm{SOCS1}},\\
\dot m_{\mathrm{CXCL9}} &= d_{\mathrm{CXCL9}}\,(1 + f_{\mathrm{CXCL9}} A_2) - d_{\mathrm{CXCL9}}\, m_{\mathrm{CXCL9}},\\
\dot m_{\mathrm{CXCL10}} &= d_{\mathrm{CXCL10}}\,(1 + f_{\mathrm{CXCL10}} A_2) + d_{\mathrm{CXCL10}} f_H H_2 - d_{\mathrm{CXCL10}}\, m_{\mathrm{CXCL10}},\\
\dot H_1 &= -k_H H_1, \qquad \dot H_2 = k_H H_1 - k_H H_2,
\end{aligned}
\tag{S13}
$$

with the initial state of model 5 and 25 estimated parameters (17 kinetic, the four $o_i$ and the four $\sigma_i$). The lab note gained a paragraph on the first run and the aim of this one; the data, the knowledge file and the required reactions were unchanged. A first launch with the original target was refused at its start, because mechanism discrimination with a single supplied model is contradictory. The target was set to parameter values, for which profiles of all parameters are computed, and criterion 1 was reworded to compare the refit with the first run; the `orchestrating agent` made these changes under the plan we had authorised.

Table S10: **The boxes that changed between model 5 (run 1) and its recalibration (run 2)**, all on $\log_{10}$ scale except $w_S$ and $w_{\mathrm{lag}}$ (linear). Unchanged: $k_{\mathrm{Sin}} \in [0.3, 30]$, $k_{\mathrm{lag}} \in [0.05, 20]$, $d_i \in [0.01, 10]$, $f_{\mathrm{SOCS1}}$, $f_{\mathrm{CXCL9}}$, $f_{\mathrm{CXCL10}} \in [0.1, 10^4]$, $f_H \in [0.01, 10^3]$, $k_H \in [0.02, 2]$, $o_i$ and $\sigma_i$.

| parameter | model 5 (run 1) | model 5, recalibrated (run 2) | run-2 estimate |
|---|---|---|---|
| $k_A$ | $[0.1, 100]$ | $[0.01, 100]$ | 0.107 |
| $k_{LA}$ | $[10^{-4}, 1]$ | $[10^{-4}, 100]$ | 4.24 |
| $g_R \to G_R = g_R k_R$ | $g_R \in [0.01, 100]$ | $G_R \in [10^{-3}, 10^3]$ | 209.1 |
| $k_R$ | $[0.01, 10]$ | $[10^{-4}, 10]$ | 0.0557 |
| $f_{\mathrm{IRF1}} \to F_{\mathrm{IRF1}} = f_{\mathrm{IRF1}}/h_{\mathrm{IRF1}}$ | $f_{\mathrm{IRF1}} \in [0.1, 10^4]$ | $F_{\mathrm{IRF1}} \in [0.1, 10^3]$ | 10.79 |
| $h_{\mathrm{IRF1}}$ | $[10^{-3}, 10^3]$ | $[10^{-3}, 10^4]$ | 581.0 |
| $w_S$, $w_{\mathrm{lag}}$ | $[0, 1]$ | fixed at 1, removed | – |

**The fit.** Run 2 reached a negative log-likelihood of 241.75 with 25 estimated parameters (AIC 533.50, BIC 628.02), against model 5's 242.50 with 27 (AIC 539.00, BIC 641.08), at an optimum outside run 1's box: in run 1's parameters, $g_R = G_R/k_R = 3.75 \times 10^3$ and $k_{LA} = 4.24$, against $g_R \le 100$ and $k_{LA} \le 1$ in run 1. $k_{\mathrm{Sin}} = 0.300\,\mathrm{h}^{-1}$ is at its lower bound and $f_{\mathrm{CXCL10}} = 8639$ near its upper bound. The recalibration changes no standardised residual of model 5 by more than 0.07 (mean 0.009), so Figure S2 holds for it as well. Run 2 is not a selection step: the difference from model 5 reflects the two fractions fixed at their bound and the widened boxes, not a new mechanism.

**Profiles.** All 25 parameters were profiled (941 points, 25 min on 19 workers; Figure S3, Table S11). A 95 % interval is the set where the negative log-likelihood lies within 1.92 of its minimum; an interval that reaches its search bound is open on that side. 17 intervals close and 8 are open. The gain $G_R$ closes but spans 4.6 decades, and the relaxation rate $k_R$ is open below ($k_R \leq 0.74\,\mathrm{h}^{-1}$, a relaxation time of at least 1.35 h). The signal-loss rate $k_{LA}$ is open down to $10^{-4}$ at a cost of only 0.84 in the negative log-likelihood, with $k_R$ then at its own lower bound: once the attenuator is free, the arrays require a cumulative attenuation but do not separate the availability $L$ from an integrating $R$. Of the degradation rates only CXCL9's closes (half-life 0.52 to 3.9 h); those of IRF1, SOCS1 and CXCL10 are open at the ceiling of $10\,\mathrm{h}^{-1}$, so only upper limits on their half-lives follow. The IRF1 saturation amplitude $F_{\mathrm{IRF1}}$ closes at 8.4 to 14.3, and the upstream rate $k_{\mathrm{Sin}}$ is open at its lower bound. The upper ends of the $G_R$, $k_{LA}$, $f_{\mathrm{SOCS1}}$ and $f_{\mathrm{CXCL9}}$ profiles are reached with $f_{\mathrm{CXCL10}}$ at its upper bound of $10^4$, a box the recalibration did not widen, so these upper limits are conditional on it. Run 1's profiles of model 5 (27 parameters, 826 points) had left $k_R$ open below ($[0.01, 3.97]\,\mathrm{h}^{-1}$) and $g_R$ across its whole box, with $g_R$, $k_{LA}$, $w_S$ and $w_{\mathrm{lag}}$ at their upper bounds at the ends of most profiles; their intervals are conditional on run 1's box.

Table S11: **Profile-likelihood intervals (95 %) of model 5 recalibrated (run 2).** Status: closed, or open at the lower or upper search bound. $G_R = g_R k_R$; $F_{\mathrm{IRF1}} = f_{\mathrm{IRF1}}/h_{\mathrm{IRF1}}$ is the induced increment of IRF1 production at saturation. Rates in $\mathrm{h}^{-1}$; offsets and $\sigma$ in $\log_2$ units.

| parameter | scale | estimate | 95 % interval | search box | status |
|---|---|---|---|---|---|
| $k_{\mathrm{Sin}}$ | log10 | 0.3 | [0.3, 10.78] | [0.3, 30] | lower-open |
| $k_A$ | log10 | 0.1071 | [0.07665, 3.324] | [0.01, 100] | closed |
| $k_{LA}$ | log10 | 4.238 | $[1 \times 10^{-4}$, 8.731] | $[1 \times 10^{-4}$, 100] | lower-open |
| $G_R$ | log10 | 209.1 | [0.01375, 563.4] | [0.001, 1000] | closed |
| $k_R$ | log10 | 0.05574 | $[1 \times 10^{-4}$, 0.7413] | $[1 \times 10^{-4}$, 10] | lower-open |
| $k_{\mathrm{lag}}$ | log10 | 1.09 | [0.07594, 15.71] | [0.05, 20] | closed |
| $d_{\mathrm{IRF1}}$ | log10 | 2.643 | [0.9711, 10] | [0.01, 10] | upper-open |
| $d_{\mathrm{SOCS1}}$ | log10 | 0.5615 | [0.08316, 10] | [0.01, 10] | upper-open |
| $d_{\mathrm{CXCL9}}$ | log10 | 0.3827 | [0.1777, 1.34] | [0.01, 10] | closed |
| $d_{\mathrm{CXCL10}}$ | log10 | 1.508 | [0.3855, 10] | [0.01, 10] | upper-open |
| $F_{\mathrm{IRF1}}$ | log10 | 10.79 | [8.437, 14.26] | [0.1, 1000] | closed |
| $f_{\mathrm{SOCS1}}$ | log10 | 240.4 | [9.434, 371.5] | [0.1, $1 \times 10^4$] | closed |
| $f_{\mathrm{CXCL9}}$ | log10 | 2958 | [118.7, 4319] | [0.1, $1 \times 10^4$] | closed |
| $f_{\mathrm{CXCL10}}$ | log10 | 8639 | [337.2, $1 \times 10^4$] | [0.1, $1 \times 10^4$] | upper-open |
| $h_{\mathrm{IRF1}}$ | log10 | 581 | [23.92, $1 \times 10^4$] | [0.001, $1 \times 10^4$] | upper-open |
| $f_H$ | log10 | 20.44 | [10.51, 41.52] | [0.01, 1000] | closed |
| $k_H$ | log10 | 0.3016 | [0.2061, 0.5328] | [0.02, 2] | closed |
| $o_{\mathrm{IRF1}}$ | lin | 7.616 | [7.468, 7.764] | [−1000, 1000] | closed |
| $o_{\mathrm{SOCS1}}$ | lin | 6.713 | [6.611, 6.814] | [−1000, 1000] | closed |
| $o_{\mathrm{CXCL9}}$ | lin | 6.318 | [6.204, 6.431] | [−1000, 1000] | closed |
| $o_{\mathrm{CXCL10}}$ | lin | 6.603 | [6.382, 6.783] | [−1000, 1000] | closed |
| $\sigma_{\mathrm{IRF1}}$ | log10 | 0.5703 | [0.4949, 0.6668] | $[1 \times 10^{-4}$, 1000] | closed |
| $\sigma_{\mathrm{SOCS1}}$ | log10 | 0.3895 | [0.3396, 0.4552] | $[1 \times 10^{-4}$, 1000] | closed |
| $\sigma_{\mathrm{CXCL9}}$ | log10 | 0.4373 | [0.3795, 0.5114] | $[1 \times 10^{-4}$, 1000] | closed |
| $\sigma_{\mathrm{CXCL10}}$ | log10 | 0.698 | [0.6058, 0.8157] | $[1 \times 10^{-4}$, 1000] | closed |

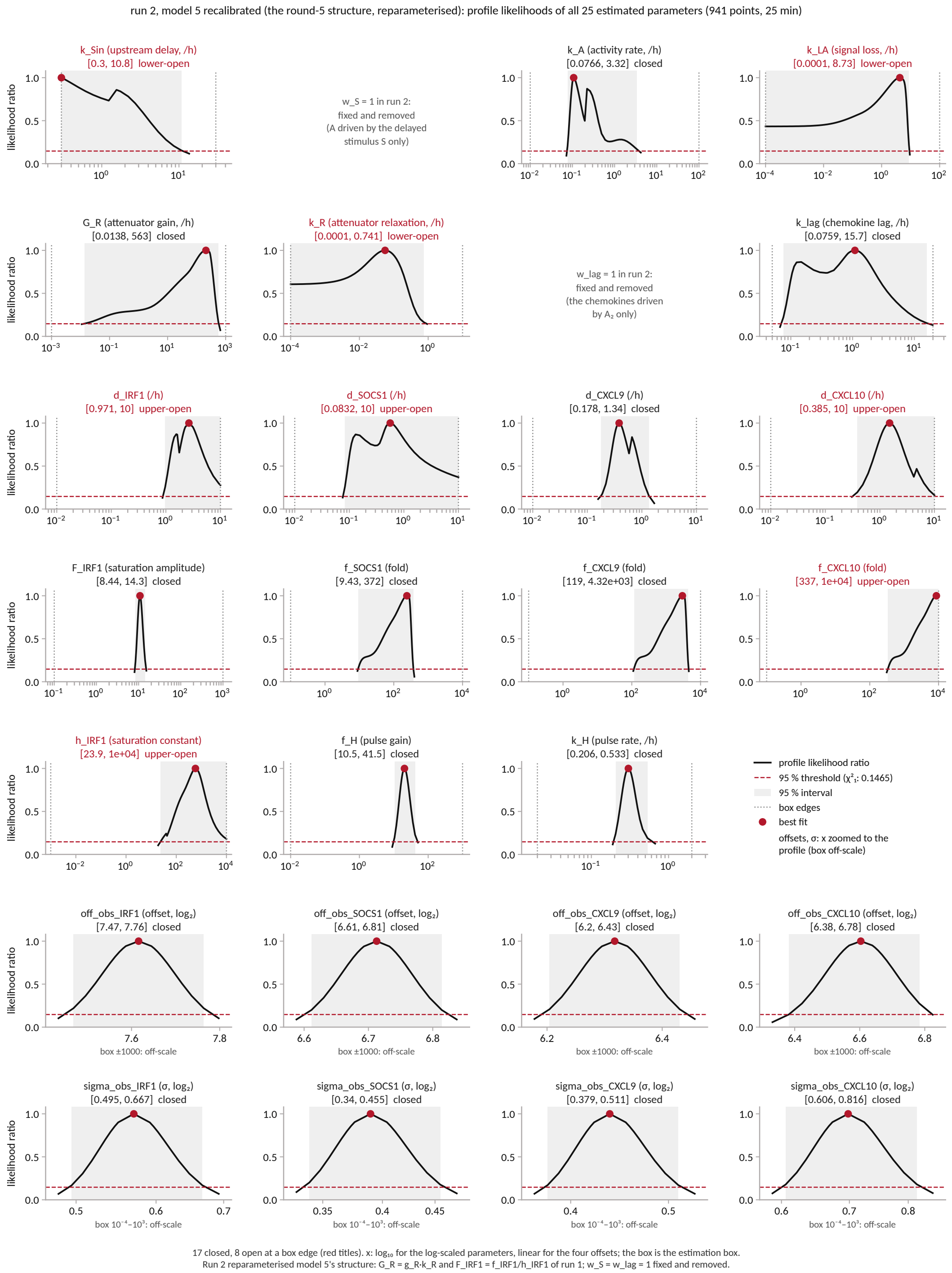

Figure S3: **Profile likelihoods of model 5 recalibrated (run 2).** All 25 estimated parameters on their estimation scale: profile likelihood ratio (black), 95 % threshold (dashed), 95 % interval (shaded), search bounds (dotted) and best fit (dot); red titles mark intervals open at a bound; offsets and $\sigma$ are drawn on the range of their profile. $w_S$ and $w_{\text{lag}}$ are fixed at 1 in run 2.

### S4 Results 3: Communication between immune cell types (PBMC)

#### S4.1 Data and noise model

**Data.** The deposited single-cell files of Bodenmiller et al. (2012) (Cytobank experiment 15711) hold one 96-well plate: column = stimulus, row = fixation time (0, 1, 5, 15, 30, 60, 120, and 240 min). All wells were barcoded, pooled, stained, and measured together, so instrument conditions are identical for all wells and cell types. We used the columns IL-3, unstimulated reference, G-CSF, GM-CSF, IFN-$\gamma$, IFN-$\alpha$, and LPS. Every channel was transformed as $\mathrm{arcsinh}(x/5)$. All values in the data are on this scale. The five cell types were gated along the authors' gating tree, each cut-off at the density minimum of the pooled data within its parent gate: $CD4^+$ T cells (26.6 % of the cells, about 2100 per well), $CD4^-$ T cells (13.4 %, the authors call this gate $CD8^+$ T cells, but the panel has no CD8 antibody), B cells (10.2 %), NK cells (24.5 %), and monocytes (14.1 %, about 1700 per well at 0 min and 76 to 520 at 240 min). The per-well median of pSTAT1, pSTAT3, and pSTAT5 in each cell type gives $7 \times 5 \times 3 \times 8 = 840$ measurements.

**Noise model.** The noise is additive and normal, with a fixed standard deviation per measurement that is the standard error of the per-well median combined with the well-to-well reproducibility; no $\sigma$ is estimated, and the cell-to-cell spread within a well is not part of it. For every row, $\sigma^2 = \sigma_{\mathrm{bw}}^2 + \mathrm{se}^2$, with $\sigma_{\mathrm{bw}}$ the well-to-well standard deviation of the median for that readout, estimated from the eleven unstimulated 0-min wells (the only replicates on the plate), and $\mathrm{se} = 1.2533 \cdot \mathrm{SD}/\sqrt{n}$ the sampling error of the median from the $n$ cells of the well. On the 42 monocyte rows at 120 and 240 min, $\sigma$ is doubled, because the monocyte gate loses most of its cells over the four hours and faster under stimulation, so the remaining monocytes are a selected subset.

#### S4.2 The knowledge file

The `knowledge agent` ran online in two calls (46 and 8 min) and the `knowledge verification agent` in one call: the `module engine` resolved 44 distinct sources (DOI, PubMed id, or URL) and fetched and cached four documents in full, and the verification agent re-classified all 123 evidence items and disagreed with the knowledge agent on one. The knowledge file contains 73 reactions (Table S12) and 25 facts (Table S13): 44 reactions graded high, 7 medium, 21 low, and 1 without evidence. The suggested floor, "low", was set by the lowest-graded supported essential reactions, the inactivation of the type-I interferon receptor in each cell type, for which the only evidence came from human cell lines (Kumar et al., 2003). We accepted it, with the module in advisory mode, in which a reaction below the floor is flagged but not refused, so that the early rounds could explore mechanisms with little evidence behind them.

Table S12: **Reactions of the knowledge file, grouped.** Number of reactions in the group, whether the `knowledge agent` marked them essential for a base model, and their grades. Grades are assigned by the `module engine` from the kind and context of the evidence (Supplementary Note S1.4).

| group | $n$ | essential | grades |
|---|---|---|---|
| IFN-$\alpha$ binding to IFNAR, one per cell type | 5 | yes | 5 high |
| IFNAR inactivation, one per cell type | 5 | yes | 5 low |
| STAT1, STAT3, and STAT5 phosphorylation by the active receptors, per cell type | 15 | yes | 14 high, 1 medium |
| pSTAT dephosphorylation, per STAT and cell type | 15 | yes | 15 low |
| IFN-$\gamma$ binding to IFNGR and inactivation (monocytes, B cells, $CD4^+$ T cells) | 6 | yes | 6 high |
| LPS binding to TLR4 and inactivation (monocytes) | 2 | yes | 2 high |
| GM-CSF, IL-3, and G-CSF binding to their receptors and inactivation (monocytes) | 6 | yes | 5 high, 1 medium |
| Type-I interferon: production driven by TLR4, clearance, binding to IFNAR in each cell type | 7 | no | 4 high, 3 medium |
| IL-6: production driven by TLR4 and the CSF receptors, clearance, IL-6R binding and inactivation ($CD4^+$ and $CD4^-$ T cells, monocytes) | 8 | no | 6 high, 2 medium |
| IFN-$\gamma$ secretion by NK cells driven by IFNAR | 1 | no | high |
| LPS binding to TLR4 on B cells | 1 | no | low |
| IL-10 and IL-27 secretion driven by TLR4 | 2 | no | 1 high, 1 none |
| all | 73 | 54 | 44 high, 7 medium, 21 low, 1 none |

Table S13: **Facts of the knowledge file.** Id, fact (shortened to its first clause, the full statement is on Zenodo), the agent role it is addressed to, grade.

| ID | Fact (shortened) | For | Grade |
|---|---|---|---|
| fact_1 | Among purified human leukocyte populations TLR4 (with TLR2 and TLR5) is restricted to myelomonocytic cells, while T, B and NK cells express TLR1 but not TLR4 | construction, condition encoding | high |
| fact_2 | The G-CSF receptor is detectable on human neutrophils and monocytes but not lymphocytes | construction, condition encoding | medium |
| fact_3 | High-affinity GM-CSF binding sites are present on human myeloid cells at 50-500 per cell and absent from human T-lineage cell lines | construction, condition encoding | medium |
| fact_4 | GM-CSF binds its receptor with Kd about 15-30 pM and half-maximal biological activity at 10-20 pM | condition encoding, construction | medium |
| fact_5 | The IFN-gamma transducing chain IFNGR2 is high on myeloid cells, moderate on B cells and poor on T cells, and its surface level determines how fast and how strongly STAT1 is activated | construction, condition encoding | high |
| fact_6 | IFNAR2 is present on essentially all human blood monocytes, B cells, CD4+ and CD8+ T cells, and all express STAT1 | construction | high |
| fact_7 | After type I interferon, STAT3 and STAT5 phosphorylation in human blood leukocyte subsets is maximal near 45 min and declines gradually, and STAT1 in CD8 T cells and monocytes is also optimal at 45 min | construction | high |
| fact_8 | Once receptor signalling stops, phosphorylated STAT1 is completely cleared within about 60 min, and under continuous stimulation nuclear pY-STAT1 peaks at 20-30 min and is gone by 4 h | construction | low |
| fact_9 | Bulk monocyte cytokine release after LPS is slower than the lymphocyte onset seen here | construction, expansion | none |
| fact_10 | LPS induces a strong IFN-beta mRNA response in human macrophages that subsides by 8 h, while in mouse macrophages the LPS-driven ISGF3 response is far greater at 4 h than 1 h because endogenous IFN-beta protein takes hours to accumulate | construction, expansion | medium |
| fact_11 | In primary human PBMC, LPS induces IFN-beta transcription and secretion with STAT1 phosphorylation and downstream ISG expression, and in human myeloid cells the late arm of that response is abolished by an IFNAR2-neutralising antibody | construction, expansion | high |
| fact_12 | GM-CSF and IL-3 induce IL-6, IL-1beta, IL-8, TNF-alpha and IL-1RA gene expression in purified human monocytes with different kinetics from LPS, while IFN-gamma alone induced none of the monocyte cytokine genes tested | construction, expansion | high |
| fact_13 | Most resting human CD4 T cells carry surface IL-6R, at higher levels than CD8 T cells, while only a small and variable proportion of CD8 T and B cells are positive | construction | high |
| fact_14 | IL-6 elicits both pSTAT3 and pSTAT1 in human T cells and monocytes within 10-30 min | construction, expansion | high |

| ID | Fact (shortened) | For | Grade |
|---|---|---|---|
| fact_15 | LPS pretreatment of primary human macrophages blocks IL-6-induced STAT3 activation rapidly and independently of transcription, while IL-10-induced STAT3 is largely unaffected | construction, expansion | medium |
| fact_16 | Human monocytes carry about 3,000-7,000 IFN-gamma binding sites per cell (Ka 4-12 x $10^8$ $M^{-1}$) and degrade about 5,000 IFN-gamma molecules per cell per hour at 37 C | construction | high |
| fact_17 | A relay through a sensing population is documented in human PBMC | construction, expansion | medium |
| fact_18 | Paracrine type I interferon from a small fraction of early responders coordinates the antiviral response of the whole population in mouse dendritic cells | expansion | low |
| fact_19 | A secreted factor in a 200 ul well is not automatically uniform | construction | none |
| fact_20 | Arcsinh with cofactor 5 is the standard transformation for mass cytometry ion counts, mapping channels to a comparable range | observation model | low |
| fact_21 | Mass-tag cellular barcoding pools all 96 wells into one sample stained and acquired together, which is why any well-to-well difference must have arisen in the well before fixation | observation model | high |
| fact_22 | The authors of the source study read the same plate as inter-cellular communication | construction, expansion | high |
| fact_23 | A modern single-cell study of human PBMC stimulated with LPS reported that only monocytes were transcriptionally activated, with no distinct interferon-stimulated-gene response in T, B or NK cells | construction, expansion | none |
| fact_24 | Handling and plating alone induce monokine mRNA in human monocytes within 30-90 min, but secretion of those mediators required a second, LPS-derived signal | construction, observation model | none |
| fact_25 | Quantitative proteomes of 28 primary human immune cell populations in resting and activated states, including secreted proteins, provide per-population copy numbers for receptors and ligands where a model needs initial values (immprot.org) | construction | high |

#### S4.3 The drift term

In the unstimulated reference well the readouts rise slowly over the four hours, mostly after 60 min, in every cell type and most in the monocytes (Figure S4). All wells were stained and measured together, so the rise happened in the wells. Other channels drift as well: pErk, pPlcg2, and pSlp76 rise and pS6 falls in every cell type. In the initial project description we decided how the drift enters every model. The modelled median count of a readout $r$ is background plus pSTAT plus drift, observed on the arcsinh scale. Before time zero the cells rest without drift, and at time zero the drift starts in every condition, the reference included, so it is the change since time zero as the data define it. The `construction agent` implemented it as one clock $C$ shared by all readouts and one drift state $D_r$ per readout,

$$\begin{aligned} \dot{C} &= k_C\,(u_D - C), && C(0) = 0, \\ \dot{D}_r &= k_r\,C - \mu\,D_r, && D_r(0) = 0, \\ y_r &= \operatorname{arcsinh}\big([\,b_r + s_r\,\mathrm{pSTAT}_r + D_r\,]/5\big), && \end{aligned} \tag{S14}$$

in which $u_D$ is 0 during pre-equilibration and 1 from time zero in every condition, so that $C$ rises from 0 towards 1 with the time constant $1/k_C$ (fitted: 60 min). The rate $k_r$ is one per readout (15), the removal rate $\mu$ is shared (fitted: $1/\mu = 500\,\text{min}$), $b_r$ is the background and $s_r$ the readout scale per phospho-site. The drift state is added to the count and does not act on the signalling. The alternative offered in round 0, basal secretion of the two factors in place of the drift, was worse when compared to models with the drift (Table S14). The fitted drift is a monotone rise. It does not follow the early rise and fall of T-cell pSTAT1 in the reference well, and it does not reach the full late rise in the monocytes, where the stimulated wells can even end below the reference at 240 min and $\sigma$ is doubled.

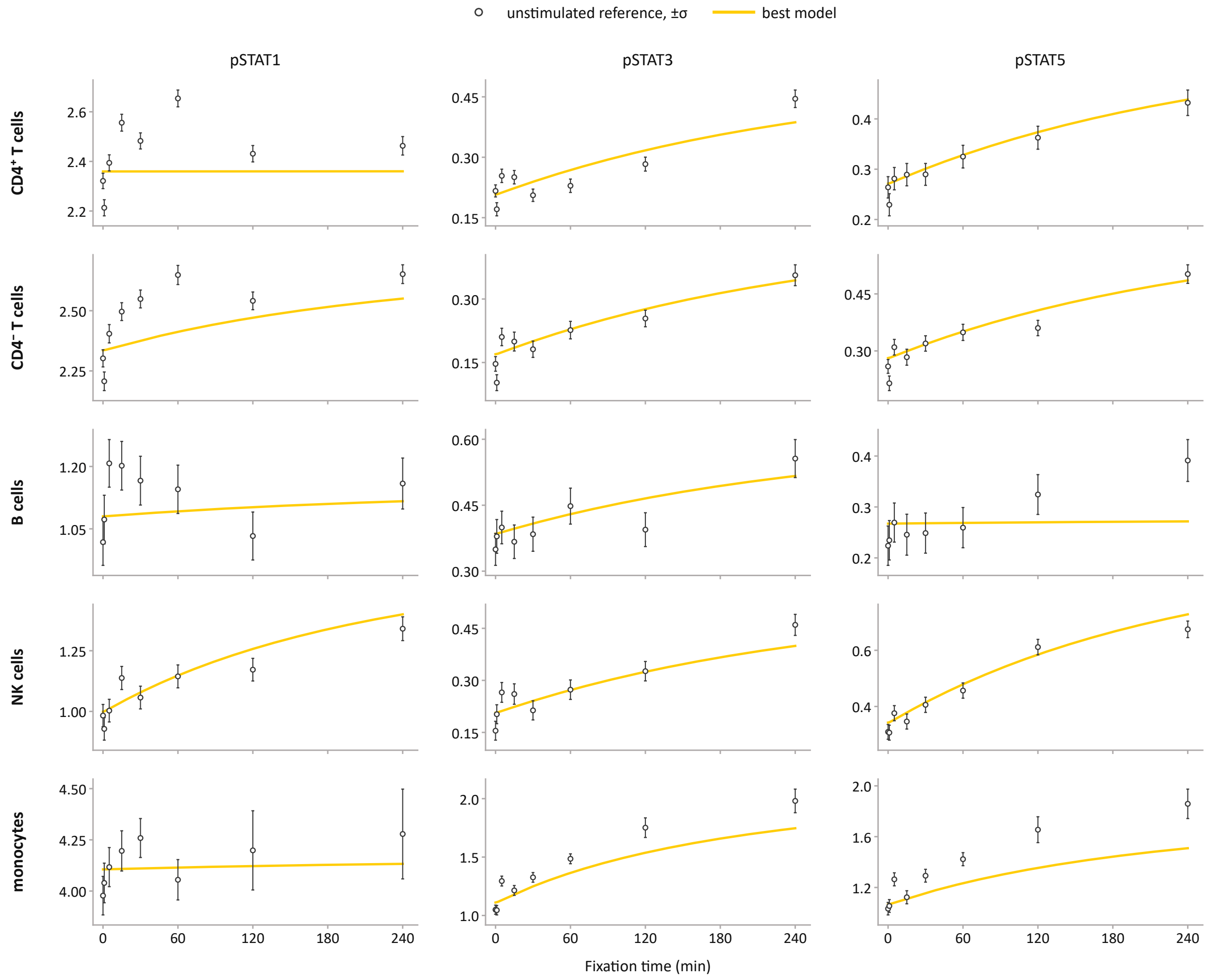

Figure S4: **The drift in the unstimulated reference well.** The 15 readouts of the reference well (open circles, $\pm\sigma$) and the best model (yellow). Rows are the five cell types, columns the three phospho-STATs, and the y range is set per panel. Without its drift term the model is flat in this well at the 0-min level.

#### S4.4 The run: five rounds and 17 models

**Staging.** The gating, the choice of stimuli, the choice of the per-well median as the fitted quantity, and the noise model were worked out by us in conversation with the `orchestrating agent`, which proposed the options and checked them against the data. The decisions are ours. In the same conversations we wrote the project description and the intent. The description gives the system, the plate, and the processing of S4.1, and what the data show: the lymphocytes respond to IFN-$\alpha$ within minutes and to LPS only after about an hour, and the reference is not flat. It names five candidate explanations for the late LPS response (a type-I interferon and IL-6 secreted by monocytes, IFN-$\gamma$ from NK or T cells, a delayed response inside the lymphocytes without a secreted factor, and combinations) and eleven decisions of ours that every model takes as given, among them the observation model of S4.3, the fixed $\sigma$, no held-out condition, the joint simulation of the five cell types of a well, and the pre-equilibration of every condition with stimulus, drift, and factor production off. The intent asks which communication between the cell types the data require, permit, or exclude, and requires a verdict for each candidate explanation.

**Inputs.** The run received the data file (840 rows with $\sigma$), the description, and the intent. It used the knowledge module online in advisory mode, a literature search before each expansion, 100 starts per candidate with Fides and forward sensitivities.

**The rounds.** Table S14 lists the 17 models in build order, with the values the run recorded and, for the seven models of the main text, the re-estimated values the main text uses. The two differ because a bug in the workflow had given every fit of the run a $\sigma$ that differed from the declared one, so those seven models were re-estimated under the declared $\sigma$ (S4.4). The four models the run selected in turn keep their order under the declared $\sigma$, so the run's decisions stand. In round 0 the `construction agent` wrote the base model and the `expansion agent` the four expansions of the main text, and the model with both factors won. The `interpretation agent` reported that IFN-$\gamma$ secreted by NK or T cells had no verdict, since no candidate had fitted it, and the `orchestrating agent` bought a further round on our standing instruction. In round 1 the `revision agent` proposed seven mechanisms. We chose five, each added on its own to the round-0 winner and fitted, and the T-cell shut-off won. The `orchestrating agent` signed that verdict and bought the next round, again on our standing instruction. In round 2 the `revision agent` proposed four mechanisms, each fitted on its own on the round-1 winner, and the direct route won over the B-cell IL-3 route. Its gain lay in the early B-cell pSTAT5 transient under IL-3 and in offsets of T-cell pSTAT1, not in the late LPS response, which is why we did not read it as a mechanism. Rounds 3 and 4 held, at our request, the control and the control with its own LPS route. The seventeenth model, the base model with the direct route, is the delayed response inside the lymphocytes without a secreted factor. It is not a candidate of the run, because no round fitted the direct route without the secreted factors. We built it from the base model and the route of round 2 and fitted it only in the re-estimation (S4.4). The run had 41 interaction points, answered by us in the conversation or by standing instructions we gave before overnight rounds. The effort is given in Table S3.

Table S14: **The 17 models in build order.** Round, candidate, and what it adds to the base of its round. "Run": estimated parameters $n_\theta$, negative log-likelihood, and AIC as recorded by the run, under the collapsed $\sigma$ of the bug described below. "Re-estimated": the values under the declared $\sigma$ for the seven models of the main text, which the main text and its figures use. In rounds 1 to 3 every candidate is the previous round's winner plus one mechanism, and the round's candidates share one supermodel encoding that carries the reactions of every mechanism of the round, switched off by parameters fixed at zero, so $n_\theta$ as counted in the run can exceed that of the exported model (the control: 147 in the run, 132 exported). The base model with the direct route is not a candidate of the run.

| | | | run | | | re-estimated | | |
|---|---|---|---|---|---|---|---|---|
| round | candidate | adds | $n_\theta$ | NLLH | AIC | $n_\theta$ | AIC | BIC |
| 0 | base model | no communication | 109 | 1268.1 | 2754.3 | 109 | 5591.4 | 6107.4 |
| 0 | type-I interferon | secreted by monocytes | 114 | 190.7 | 609.5 | – | – | – |
| 0 | IL-6 | secreted by monocytes | 126 | 22.8 | 297.7 | – | – | – |
| 0 | both factors | both secreted factors (selected) | 131 | −348.0 | −433.9 | 131 | 537.0 | 1157.1 |
| 0 | basal secretion | both factors, basal secretion in place of the drift | 116 | 105.9 | 443.9 | – | – | – |
| 1 | basal type-I interferon | basal secretion of the interferon beside the drift | 132 | −375.8 | −487.7 | – | – | – |
| 1 | IFN-$\gamma$ from NK cells | IFN-$\gamma$ secreted by NK cells | 135 | −402.7 | −535.4 | – | – | – |
| 1 | IL-6 from IFNAR | IL-6 production also driven by the interferon receptor | 132 | −445.4 | −626.7 | – | – | – |
| 1 | T-cell shut-off | STAT-specific shut-off of the response in T cells (selected) | 135 | −474.3 | −678.6 | 135 | 34.6 | 673.7 |
| 1 | SOCS3 brake | a gp130-driven SOCS3 brake | 134 | −348.0 | −427.9 | – | – | – |
| 2 | direct route | LPS and the CSFs act on the lymphocytes directly (selected, the best model) | 154 | −685.0 | −1061.9 | 154 | −347.9 | 381.0 |
| 2 | B-cell IL-3 route | a direct response of B cells to IL-3 | 138 | −658.7 | −1041.4 | – | – | – |
| 2 | monocyte shut-off | the STAT-specific shut-off also in monocytes | 138 | −474.3 | −672.6 | – | – | – |
| 2 | IL-27-like factor | a third secreted factor | 148 | −558.1 | −820.3 | – | – | – |
| 3 | control | the best model with the production of both secreted factors switched off | 147 | −127.8 | 38.5 | 132 | 1692.5 | 2317.3 |
| 4 | control + own LPS route | the control with its own direct route for LPS in each lymphocyte cell type | 164 | −281.5 | −234.9 | 149 | 1093.6 | 1798.9 |
| – | base + direct route | the base model with the direct route, no secreted factor | – | – | – | 128 | 1961.0 | 2566.9 |

**The noise-model bug and the re-estimation.** A bug in the assembly code of M4, fixed since, wrote the per-row $\sigma$ column of the measurement table as text, which the PEtab import maps to one noise parameter per observable and condition, so that every fit of the run used the 240-min $\sigma$ of each time series for all its eight time points. We therefore re-estimated the seven models of the main text outside the workflow under the declared $\sigma$, with the same software as the run and 200 starts per model. Table S14 gives the results, and the waterfall plots, gradients, and parameters at bounds of every re-estimated model are on Zenodo.

#### S4.5 The best model and the control models under LPS

**The best model.** The best model is the base model with both secreted factors, the T-cell shut-off, and the direct route, as drawn in the models figure of the main text: 93 species, 180 reactions, and 154 estimated parameters. Two facts of the fitted model are not visible in the drawing. The production of the type-I interferon in monocytes is driven by TLR4 only at the lower bound of its rate, and by the monocytes' own IFNAR at a rate of 0.21, so the interferon amplifies itself through the monocytes, from about $10^{-7}$ at 15 min to 0.2 at 150 min, and this loop is what delays the lymphocyte response. The direct route is fast: its transcript and protein turnover rates sit at the upper bound of $1\,\mathrm{min}^{-1}$, so DR plateaus within 5 min, and in $CD4^+$ T cells it feeds pSTAT1 only.

**The control models.** The control is the best model with the two secretion reactions of the monocytes replaced by zero rates and the 22 parameters that then act on nothing fixed, leaving 132 estimated parameters. In the control, LPS and the CSFs use one shared route with the same turnover rates and the same STAT couplings, so the LPS response cannot have a timing of its own. The control with its own LPS route adds to it, in each lymphocyte cell type, a hypothetical LPS receptor with binding and unbinding, a transcript it induces, the protein made from it, and that protein's phosphorylation of STAT1, STAT3, and STAT5, in 36 reactions with shared rates and one coupling per STAT and cell type, for 149 estimated parameters. The workflow built both at our request in rounds 3 and 4, each as the only candidate of its round, and Figure S5 shows their fits under LPS beside the best model.

#### S4.6 Comparison with the agent-only analyses

**Setup.** The PBMC problem was analysed two more times without Octomod, once with Claude Code and once with Codex, by co-authors not involved in the Octomod run, on separate machines and accounts (Methods). All three analyses received the same description and data file. For each agent-only analysis, an execution prompt was first generated by a higher-tier model of the same framework from the description, the data, and an initial prompt-generating prompt, instructing the agent to act as the orchestrator of a multi-agent process with separate instances for model building, estimation, testing, verification, and critical review, to keep records of every decision, and to run without input from a user. Both analyses had web access, and neither the model format nor the estimation tools were prescribed. Table S15 summarises the three analyses.

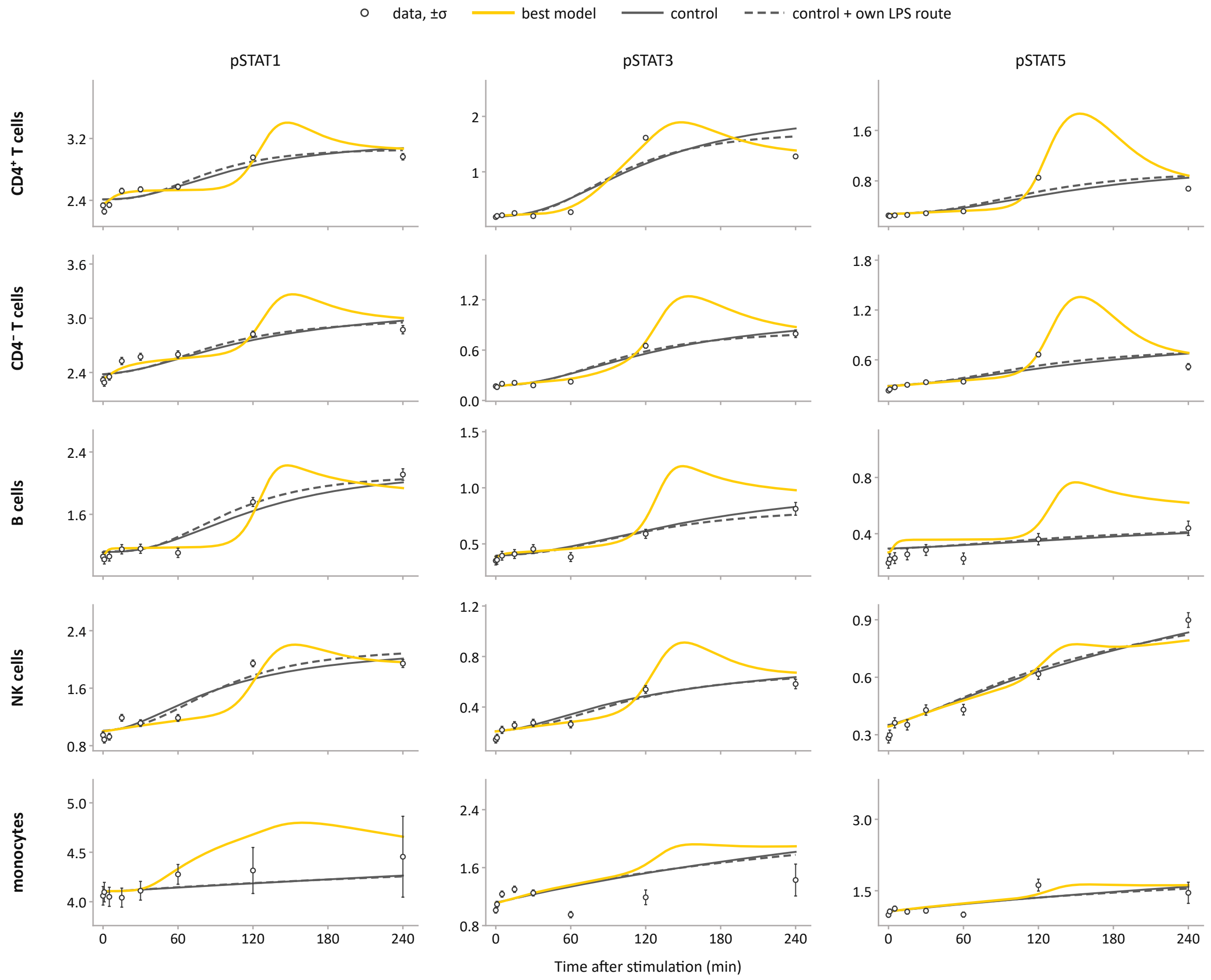

Figure S5: **Fits under LPS of the best model and the two control models.** Data as open circles with $\pm\sigma$, the best model in yellow, the control in grey, and the control with its own LPS route in dashed grey. Rows are the five cell types, columns the three phospho-STATs, and the y ranges are those of the fits figure of the main text.

Table S15: **Setups of Octomod and two agent-only control runs applied to the Bodenmiller et al. (2012) use case.**

| | Octomod | Claude Code only | Codex only |
|---|---|---|---|
| Structure given by | `central engine` and `module engines` (a program) | execution prompt | execution prompt |
| LLM | Claude Opus 5 (`claude-opus-5`) | Claude Opus 5.5 (prompt generated by Claude Fable 5.1) | GPT6-Sol, medium effort (prompt generated by GPT6-Astra, ultra effort) |
| Language and tools | Python with SBML, PEtab, AMICI, and pyPESTO | Julia, own code | Python, own code with SciPy |
| Models fitted | 17 (16 in five rounds, one outside the run) | 32 (57 specified) | none converged |
| States | 64 to 93 species | 99 (selected model) | 38 |
| Estimated parameters | 109 to 154 | 101 to 190 | 122 to 156 |
| STAT species in the model | yes | no | yes (fractions) |
| ODE solver | CVODES (AMICI) | OrdinaryDiffEq.jl | SciPy (BDF) |
| Optimiser | Fides | Levenberg–Marquardt, own multi-start | trust-region least squares |
| Starts per model | 100 | 10 to 40 | 1 |
| Derivatives | forward sensitivities | forward-mode automatic differentiation | finite differences |
| Limit per start | 1000 iterations | 3000 iterations or 30 min | 20 evaluations or 30 min |
| Output format | SBML, PEtab, run record | own files | own files |
| User during the run | at the interaction points | none | none |
| Wall time | 140 h (103 h computation) | 15 h on 24 CPUs | 5 h, single process |

**Verification.** For the Claude Code analysis, we recomputed the objective value of all 32 fitted models from their stored predictions and the original data file, and every value agrees with the reported one. We re-implemented its selected model in Python from the Julia source and simulated it with the stored parameters: the predictions agree to $2 \times 10^{-10}$ and reproduce the reported $\chi^2$ of 2516.8. For the Codex analysis, we recomputed the reported objective value from the stored predictions. In both analyses, our decisions on the observation model, $\sigma$, the joint simulation of the five cell types, and the resting state are implemented as specified.

**Models.** The Octomod models are reaction networks with named species, including the STATs, in SBML and PEtab. The Claude Code models have no STAT species: every state is the active or internalised fraction of a receptor complex or a stage of a transit chain, and each pSTAT readout is a weighted sum of such states. Its selected model (Figure S6) has, in each lymphocyte cell type, one six-stage chain for LPS and one for the CSFs with a free input strength per stimulus and cell type, and each pSTAT readout has a free amplitude per readout and cell type, 99 states and 172 parameters in all. The analysis's own caveat and the knowledge file of the Octomod run say the same thing about it: a delayed response with a free amplitude per cell type and readout is equivalent to a mediator with free effects. The Codex model without a secreted factor has 38 states and 129 parameters, with a direct route of two first-order stages.

**Estimation.** All three analyses use the same data and $\sigma$, so their objective values are on one scale. In the Claude Code analysis, each model was fitted from 10 to 40 starts, each limited to 3000 iterations or 30 minutes, and 57 % of the starts ended at a limit. Its own convergence criterion, three starts within 0.1 of the best, was met for 9 of the 32 models. The analysis selected `M-null-lean-basal-drift` by BIC, and its lowest-AIC model is `M-null-basal-drift` (190 parameters, AIC $-1167.9$). Figure S7 places both beside the Octomod models, and Figure S8 shows the fits of the selected model beside the Octomod best model. In the Codex analysis, no parameter estimation converged: seven runs with one start each ended at a self-imposed evaluation or time limit, and the best point of its model without a secreted factor has $\chi^2 = 7105$. The AIC and BIC of all 32 Claude Code models are on Zenodo.

**Conclusions of the two analyses.** Table S16 sets the verdicts of the Octomod run and the Claude Code analysis side by side on the questions both addressed. The Codex analysis reports observations on the data only and draws no conclusion from a model.

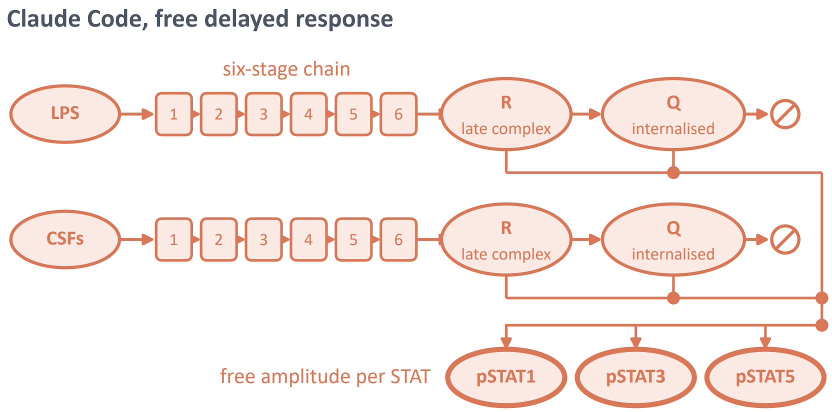

Figure S6: **Claude Code's selected model (`M-null-lean-basal-drift`), drawn in the style of the models figure of the main text.** In each lymphocyte cell type, a six-stage chain for LPS and one for the CSFs with a free input strength $w$ per stimulus and cell type. The last stage activates a late complex $R$, which is internalised to $Q$ (rate $k_{\mathrm{int}}$) and recycled to the free pool ($k_{\mathrm{rec}}$), and each pSTAT readout is $B\,(R + \varphi Q)$ with a free amplitude $B$ per readout and cell type. The model has no STAT species and no secreted factor. Not drawn: the receptor modules for IFN-$\alpha$ and IFN-$\gamma$ in every cell type, TLR4 and the CSF receptors on monocytes, a basal chain in monocytes, and a drift on every readout.

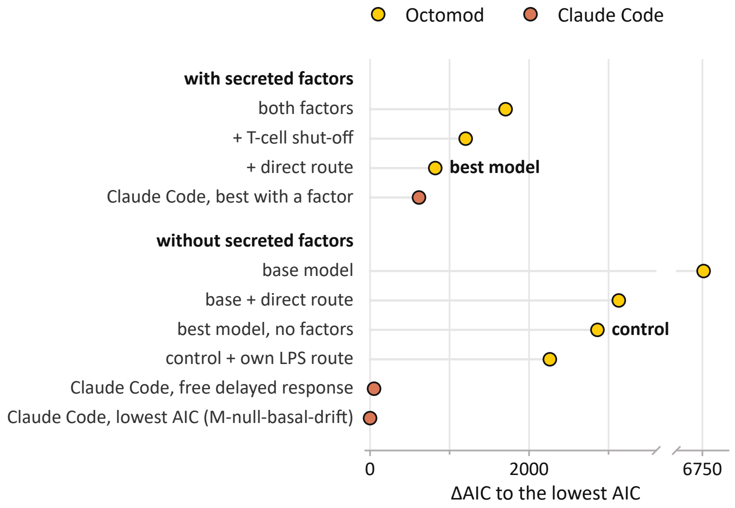

Figure S7: **ΔAIC of the Octomod models and the Claude Code models.** The seven Octomod models of the main text, Claude Code's best model with a secreted factor, its selected model (the free delayed response), and its lowest-AIC model (`M-null-basal-drift`), each relative to the lowest AIC of all (−1167.9). The models are grouped by whether they contain secreted factors, as in the models figure of the main text, and the AIC is on the likelihood scale of Table S14.

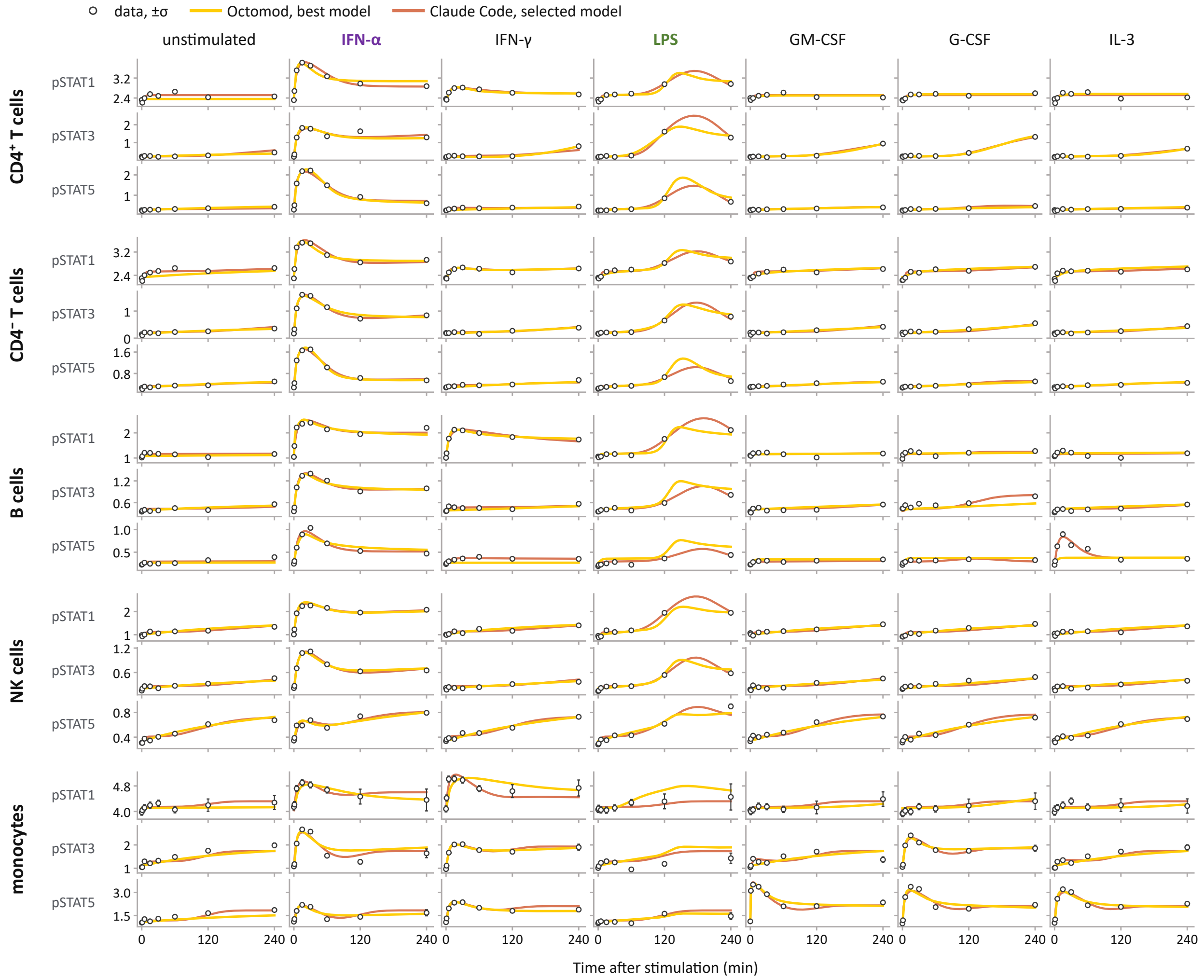

Figure S8: **All 840 measurements with the Octomod best model (yellow) and Claude Code's selected model (orange).** Columns are the seven conditions, rows the five cell types and the three phospho-STATs, and the data are open circles with $\pm\sigma$.

Table S16: **Verdicts of the Octomod run and the Claude Code analysis.**

| Question | Octomod | Claude Code |
|---|---|---|
| Is a late response of the lymphocytes needed at all? | yes, the base model without one is 5900 AIC behind the best model | not tested |
| Which secreted factors does the best model with communication contain? | a type-I interferon and IL-6 from monocytes | the same two |
| Is one secreted factor enough? | no, the second factor is required | no |
| Does the type-I interferon trigger the production of IL-6? | yes, proposed by the `revision agent`, it improves the fit, but it was not selected and is not in the best model | yes, the only mechanism explaining the second wave under IFN-$\alpha$ |
| Is the drift in the unstimulated wells caused by basal secretion of the factors? | no | no |
| Do B cells respond to IL-3 directly? | proposed by the `revision agent` and fitted, not in the best model | yes, part of every model |
| Do NK cells secrete IFN-$\gamma$ that acts on the other cell types? | improves the fit, not selected | no improvement, excluded as the only signal |
| Is communication required? | yes, within the models of the run every model without secreted factors is at least 1441 AIC behind | cannot be decided, its selected model has no factor and fits best |

### S5 Results 4: Hypothesis testing on unpublished sphingolipid data

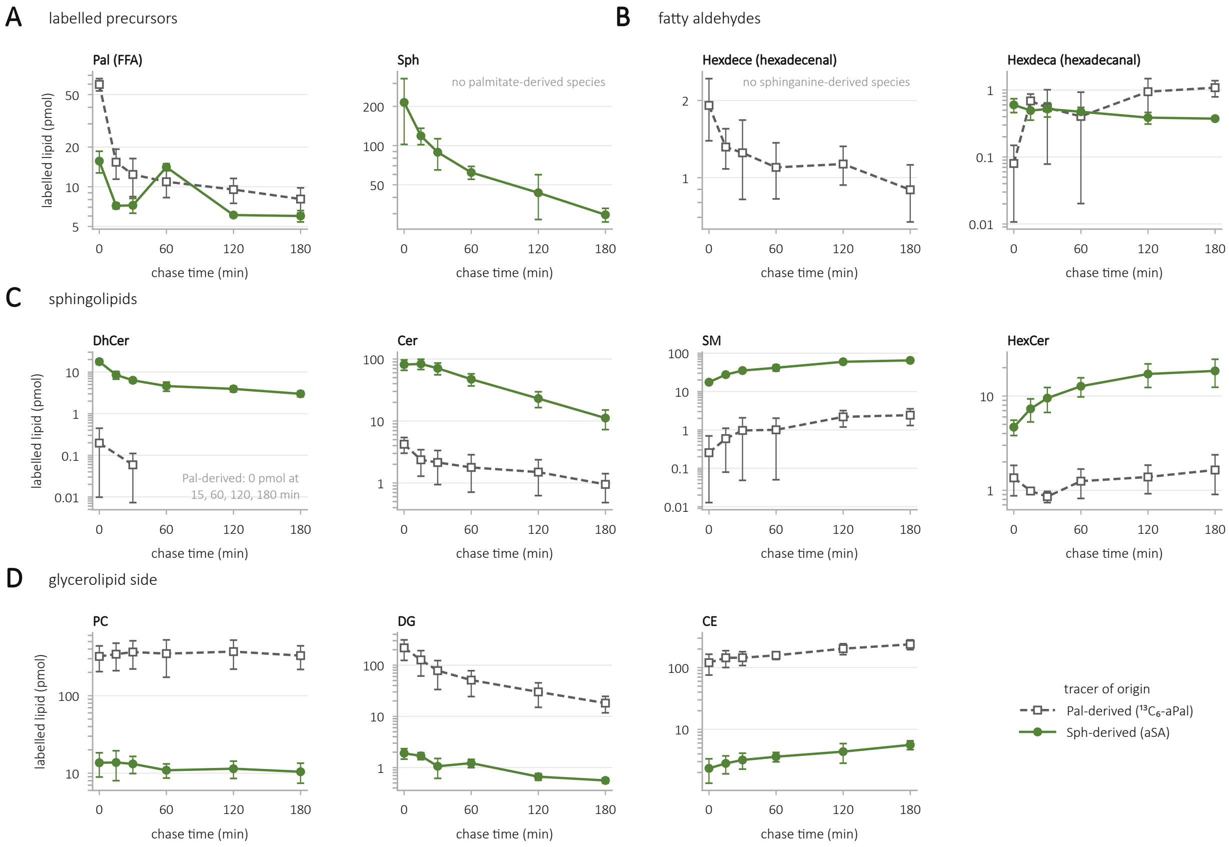

Figure S9: **Labelled lipid classes of the sphingolipid data by tracer of origin.** Basal condition, pmol per sample. (A) Labelled precursors: free fatty acids (Pal; all chain lengths) and sphinganine (Sph). (B) Fatty aldehydes: hexadecenal (Hexdece) and hexadecanal (Hexdeca). (C) Sphingolipids: DhCer, Cer, SM and HexCer. (D) Glycerolipids: PC, DG and CE. Species are assigned to Pal-derived ($^{13}C_6$-alkyne palmitate; squares, dashed) or Sph-derived (alkyne sphinganine; circles, solid) label and summed per class and sample. The median over three technical replicates gives one value per biological replicate; points show the mean $\pm$ SD over three biological replicates. Values are not normalised to the total label at $t = 0$ and therefore differ from the fitted fractions (Fig. 7C of the main text). The models were fitted to the sum of both tracers, not to this split. Logarithmic y-axes; lower error bars are truncated at 5 % of the mean, and means of zero are listed in the panel.

### References

Bodenmiller, B., Zunder, E. R., Finck, R., Chen, T. J., Savig, E. S., Bruggner, R. V., Simonds, E. F., Bendall, S. C., Sachs, K., Krutzik, P. O., and Nolan, G. P. (2012). Multiplexed mass cytometry profiling of cellular states perturbed by small-molecule regulators. *Nature Biotechnology*, 30(9):858–867.

Callenbach, A., Dorešić, D., Düster, R., Nakonecnij, V., Dudkin, E., Geyer, M., and Hasenauer, J. (2026). Quantitative modelling of P-TEFb mediated CTD phosphorylation identifies local cooperativity. *PLOS Computational Biology*, 22(7):e1014531.

Kumar, K. G. S., Tang, W., Ravindranath, A. K., Clark, W. A., Croze, E., and Fuchs, S. Y. (2003). $SCF^{HOS}$ ubiquitin ligase mediates the ligand-induced down-regulation of the interferon-$\alpha$ receptor. *The EMBO Journal*, 22(20):5480–5490.

Xue, J., Schmidt, S. V., Sander, J., Draffehn, A., Krebs, W., Quester, I., De Nardo, D., Gohel, T. D., Emde, M., Schmidleithner, L., Ganesan, H., Nino-Castro, A., Mallmann, M. R., Labzin, L., Theis, H., Kraut, M., Beyer, M., Latz, E., Freeman, T. C., Ulas, T., and Schultze, J. L. (2014). Transcriptome-based network analysis reveals a spectrum model of human macrophage activation. *Immunity*, 40(2):274–288.
